# Glycoengineering of nematode antigens using insect cells: a promising approach for producing bioactive vaccine antigens of the barber’s pole worm *Haemonchus contortus*

**DOI:** 10.1101/2025.04.11.646772

**Authors:** Isabella Adduci, Floriana Sajovitz-Grohmann, Licha N Wortha, Zuzanna Dutkiewicz, Hugo Weidinger, Sonja B Rohrer, Anja Joachim, Thomas Wittek, Dirk Werling, Iain BH Wilson, Katharina Lichtmannsperger, Shi Yan

## Abstract

The H11 antigens, located on the intestinal microvilli of *Haemonchus contortus*, comprise a group of homologous aminopeptidases essential for the parasite’s digestion of blood meals. Native H11 proteins are promising vaccine antigens, capable of eliciting robust protective immunity against *H. contortus* in sheep and goats. However, recombinant forms of H11, produced either in conventional expression systems or in transgenic *Caenorhabditis elegans*, failed to replicate the protective efficacy of the native form, most likely due to two critical factors: improper glycosylation and protein misfolding. To address these limitations, we developed a novel strategy to produce recombinant *Haemonchus* antigens in glycoengineered insect cells. By introducing three *C. elegans* genes that alter the native N-glycosylation pathways of Hi5 insect cells we successfully expressed soluble H11 and GA1 antigens featuring nematode-specific glycan epitopes, including tri-fucosylated structures and the Galβ1,4Fuc motif. The glycoengineered H11 proteins retained aminopeptidase activity and stimulated cytokine secretion from ovine peripheral blood mononuclear cells *in vitro*. These findings establish a platform for producing bioactive vaccine antigens against the parasitic nematode *H. contortus*.

## Introduction

*Haemonchus contortus*, a hematophagous parasitic nematode, significantly impacts small ruminant production leading to substantial economic losses and compromised animal welfare worldwide. Current control strategies rely heavily on anthelmintic drugs. However, these are associated with significant drawbacks, including ecotoxic effects, mandatory withdrawal periods, and the rapidly increasing prevalence of resistance to all classes of anthelmintic compounds [1,2].

Vaccination is considered an effective strategy for parasite control. Since the late 1980s, several vaccine candidates have been validated as effective against *H. contortus* infection [3], and among them, two antigens lectin affinity-purified from the intestinal fraction of the adult parasites, namely H11 and H-gal-GP, displayed superior efficacy. H11 primarily contains a family of membrane-bound aminopeptidases [4], and H-gal-GP is a multi-enzyme complex consisting of a group of aspartyl proteases, metalloproteases, and cysteine proteinases [5]. When tested separately as vaccine candidates, each antigen induced a strong protective immune response in sheep and goats, as judged by significant reduction of parasite egg shedding and worm burden [4,6]. In 2014, the Barbervax^®^ vaccine, containing both H11 and H-gal-GP, was commercialised and licensed in Australia for the use in sheep [7]. However, these native antigens must be isolated from adult worms derived from donor sheep upon slaughter, as adult *H. contortus* cannot be obtained by *in vitro* cultivation. Therefore, this approach does not only raise biosecurity concerns but can also impact on animal welfare due to the necessity to experimentally infect animals for large-scale antigen production.

The major isoforms of *H. contortus* H11 aminopeptidases, including H11, H11-1, H11-4 and H11-5, have previously been produced recombinantly using different expression hosts, including bacteria, Sf9 and Sf21 insect cells, and the free-living nematode *Caenorhabditis elegans* [8–11]. However, none of the recombinant H11 variants could sufficiently reduce worm burden and faecal egg excretion in challenged animals compared to the native form. A potential explanation is the N-glycosylation of native H11 antigens, which results in unusual structures with up to three fucose residues and the formation of a stage-specific tri-fucosylated core [12,13]; in adult worms, two of the three core fucose residues can be further galactosylated, forming two types of Gal-Fuc epitopes [14]. Such core modifications have been observed in some nematode species [15], but they are absent in mammals (**Fig. 1A**). Thus, we hypothesised that the core tri-fucosylated N-glycans are part of the conformational epitopes of intact H11 antigens. A lack of such parasite-specific post-translational glycan modifications and suboptimal protein folding may be the major cause of the recombinant H11s’ inadequate efficacy alongside with other possible reasons. Another limitation in previous attempts to develop effective recombinant H11-based vaccines may be the exclusion of essential H11 isoforms. Previous studies included a maximum of two major isoforms in vaccine formulations [9–11], which might result in an inefficient antigenic targeting.

**Fig 1.**
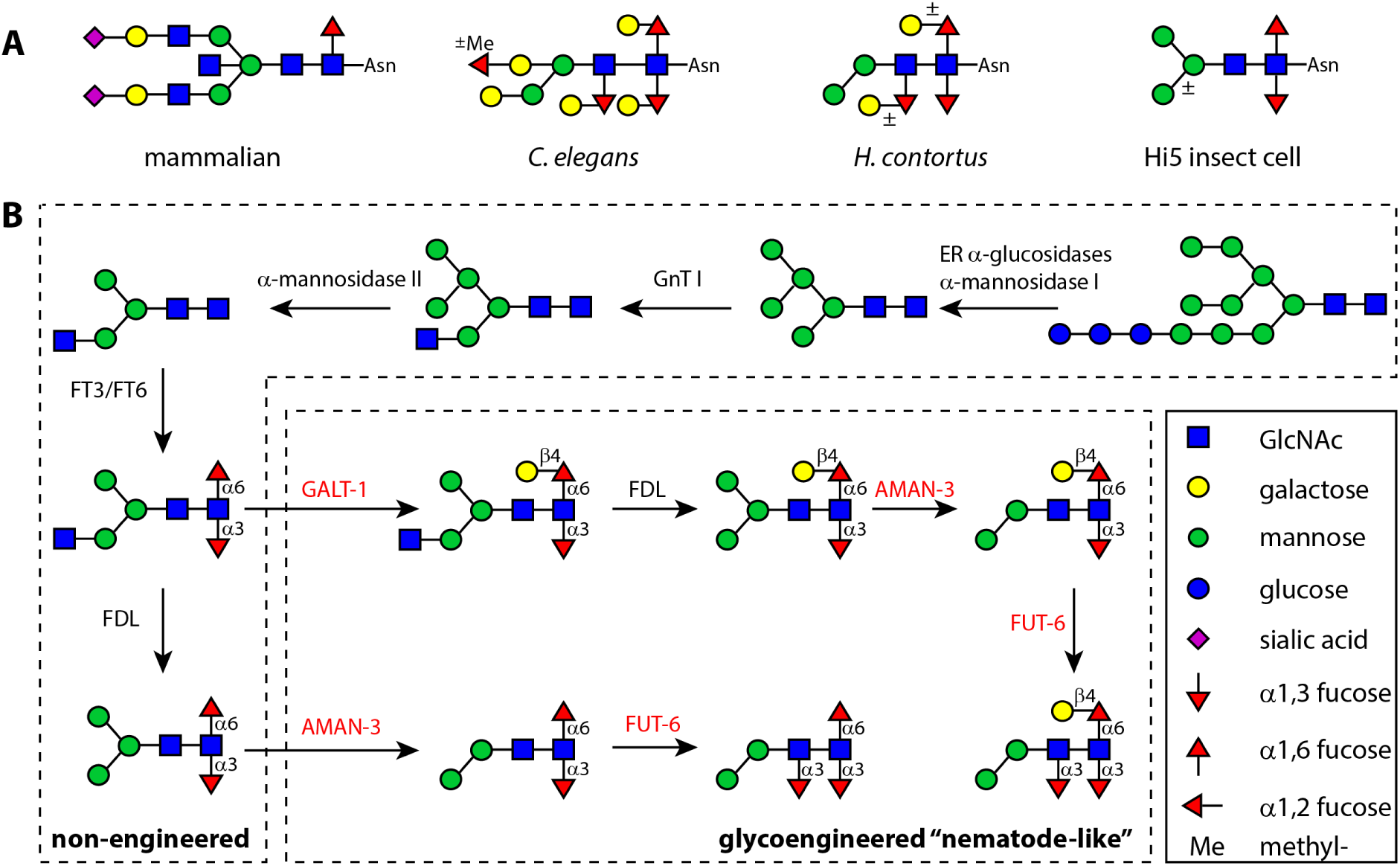
Distinct N-glycan core modifications and the concept of N-glycosylation engineering. (A) A comparison of typical N-glycan core modifications of mammalian, *C. elegans*, *H. contortus* and insect glycoproteins. (B) Major biosynthetic pathways of N-glycans in non-engineered and glycoengineered Hi5 cells. Foreign glycoenzymes of nematodes are shown in red whereas the native insect enzymes are in black. Glycan structures are drawn in symbol nomenclature for glycans (SNFG) [57] and the monosaccharides are annotated in a solid box.

To address these points, we redesigned the antigen production strategy and used a novel and advanced glycoengineering approach, employing the *Trichoplusia ni*-derived Hi5 insect cells as expression host (**Fig. 1B**). The native Hi5 cell line is capable of producing secreted glycoproteins carrying abundant di-fucosylated N-glycans [16,17], which is the basis for introducing novel N-glycan biosynthetic pathways. Previous studies have shown that three essential enzymes of the model nematode *C. elegans*, the Golgi mannosidase III AMAN-3 [18], the core α1,3-fucosyltransferase FUT-6 [19] and the β1,4-galactosyltransferase GALT-1 [20], participate the biosynthesis of tri-fucosylated glycans. We therefore sought to introduce the three relevant genes (*aman-3*, *fut-6* and *galt-1*) into Hi5 cells to enable nematode-type glycan modifications on recombinant *Haemonchus* antigens.

In this study, we investigated the expression pattern of native H11 antigens in a local *H. contortus* strain (KB strain from southern Germany) using a custom antibody and molecularly identified seven H11 homologs in this strain. Using Hi5 insect cells, we produced two sets of recombinant *H. contortus* antigens, including the glycoengineered variants modified with the desired nematode-type N-glycans, and evaluated their bioactivities. This work established the basis for producing H11-based glycoengineered antigens and facilitating efficacy assessments against *H. contortus* in a controlled animal trial. It paves the way of developing glycoengineered vaccines against a broader spectrum of metazoan pathogens.

## Results

### Expression of H11 antigens is stage-specific

To better understand the expression patterns of native H11 antigens in *H. contortus*, we sought to identify conserved sequences for generating H11-specific polyclonal antibodies. Sequence alignment initially revealed five conserved motifs among the H11 variants, ranging from 6 to 22 amino acids (**S1 Fig**). Using AlphaFold-generated 3D-models, the predicted locations of these motifs were visualised, and two motifs were selected for antibody targeting: an exopeptidase motif (GAMENWGLITYRE) located within the molecule and a surface-exposed motif (EPEKYRHPKYGFKWDVPLWYQE) (**Fig 2A and 2B**). Two H11-specific polyclonal antibodies obtained from rabbit sera, targeting either a 16-aa (C+EPEKYRHPT/KYGFKWD) or a 14-aa (C+GAMENWGLITYRE) synthetic oligopeptide, were evaluated using Western blot. Only the antibody targeting the 16-aa motif recognised the intact form of H11 antigens of 110 kDa in size. This signal was detectable exclusively in adult-stage worms and was absent in infectious L3 larvae (**Fig 2C**). Further analysis using lysates of other nematodes (*C. elegans*, *Ascaris suum* and *Oesophagostomum dentatum*) indicated that the binding was specific to *H. contortus* (**S2 Fig**). Therefore, the 16-aa targeting antibody was considered reliable and specific for detecting H11, and was used subsequently for probing recombinant H11 products in insect cell cultures.

**Fig 2.**
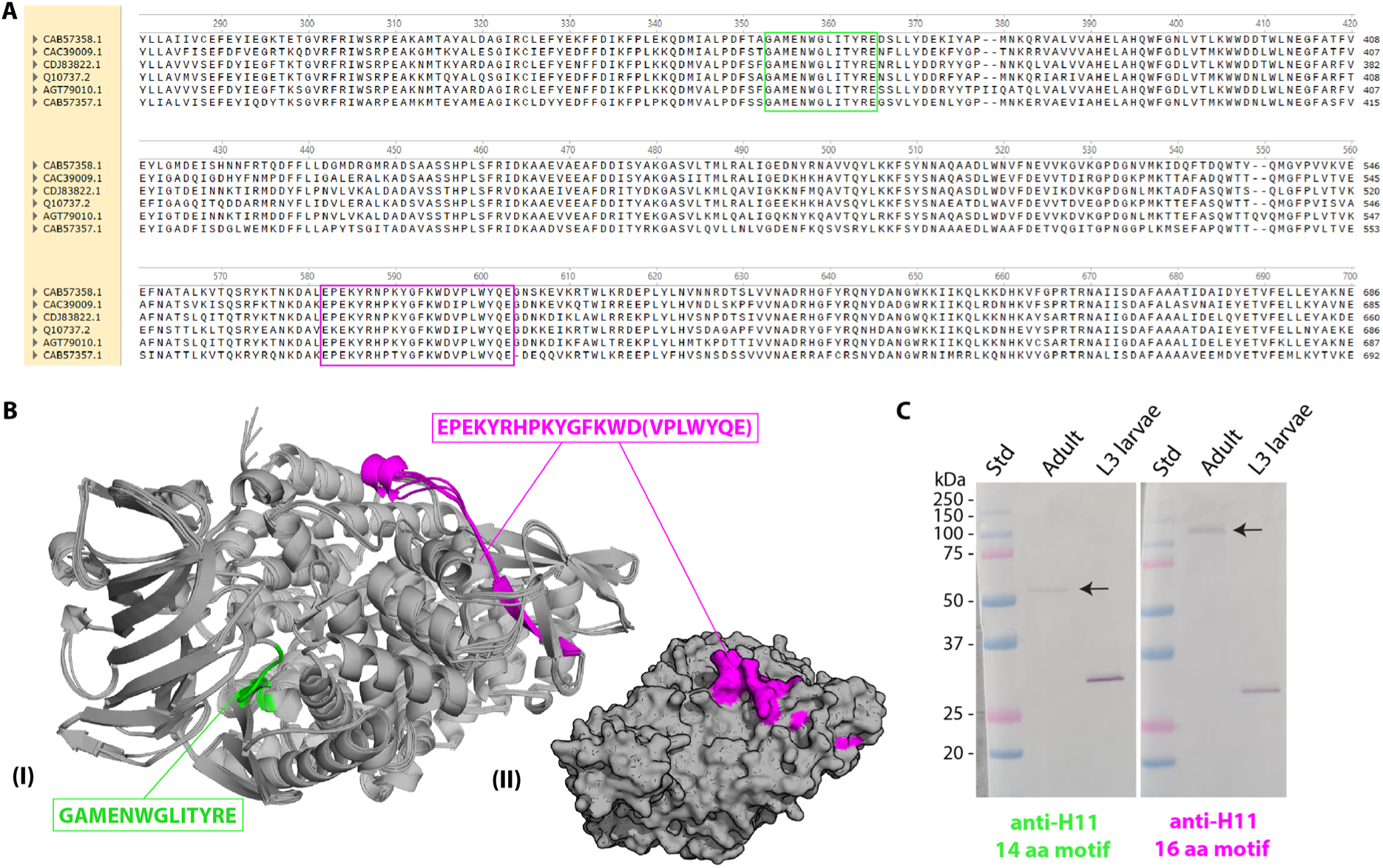
Identification of two conserved H11 motifs for polyclonal antibody preparation. (A) Multiple sequence alignment of H11 isoforms using published protein sequences (Genbank accession nos. CAB57357.1, CDJ83822.1, AGT79010.1, CAB57358.1, CAC39009.1, Q10737.2). The two conserved regions used for polyclonal antibody generation are highlighted in green (GAMENWGLITYRE) and magenta (EPEKYRHPKYGFKWD(VPLWYQE)). (B) The 3D-structure model of 6 H11 proteins, as predicted using AlphaFold; (I) representation of the structure, with the conserved motifs highlighted in green and magenta, showing their location within the protein chain; (II) surface representation of the H11 protein, with the magenta motif used for anti-H11 polyclonal antibody generation. (C) Western blot showing the reactivity of anti-H11 polyclonal antibodies generated against the two conserved motifs, using native antigens from *H. contortus* adults and third-stage (L3) larvae.

### Seven H11-encoding genes are transcribed in adult H. contortus

In order to identify major isoforms of the H11 ‘hidden’ antigens in the adult worm, we re-investigated the published genome data of *H. contortus* (Project: PRJEB506; MHco3(ISE) strain) [21]. Sequence alignment suggested a cluster of seven genes on chromosome 5, predicted to encode H11 aminopeptidase (**Fig 3A**; gene IDs: HCON_00156230, HCON_00156240, HCON_00156250, HCON_00156260, HCON_00156270, HCON_00156280 and HCON_00156285). Genes encoding five homologs out of the seven isoforms, namely H11 (GenBank accession number: X94187.1), H11-1 (AJ249941.1), H11-2 (AJ249942.2), H11-4 (AJ311316.1) and H11-5 (KF381362.1) were reported to be transcribed during parasite development [8,9]. To verify that all seven homologues are also present in the used *H. contortus* strain, we PCR-amplified the coding regions using cDNAs of the KB strain as template and performed sequence analyses. Amino acid sequences of putative H11 isoforms were determined based on DNA sequencing results of either PCR amplicons or plasmid DNAs obtained by TOPO cloning (**S1 Text**). In total, seven H11 isoforms were determined, namely KB H11, KB H11-1, KB H11-2, KB H11-4, KB H11-5a, KB H11-5b and KB H11-5c, which displayed high sequence identities (91.83% - 99.79%) compared to the ones in the MHco3(ISE) strain (**Fig 3B**). Phylogenetic analysis suggested the presence of five major branches, with branches a, b and c of H11-5 being highly homologous (**Fig 3C**).

**Fig 3.**
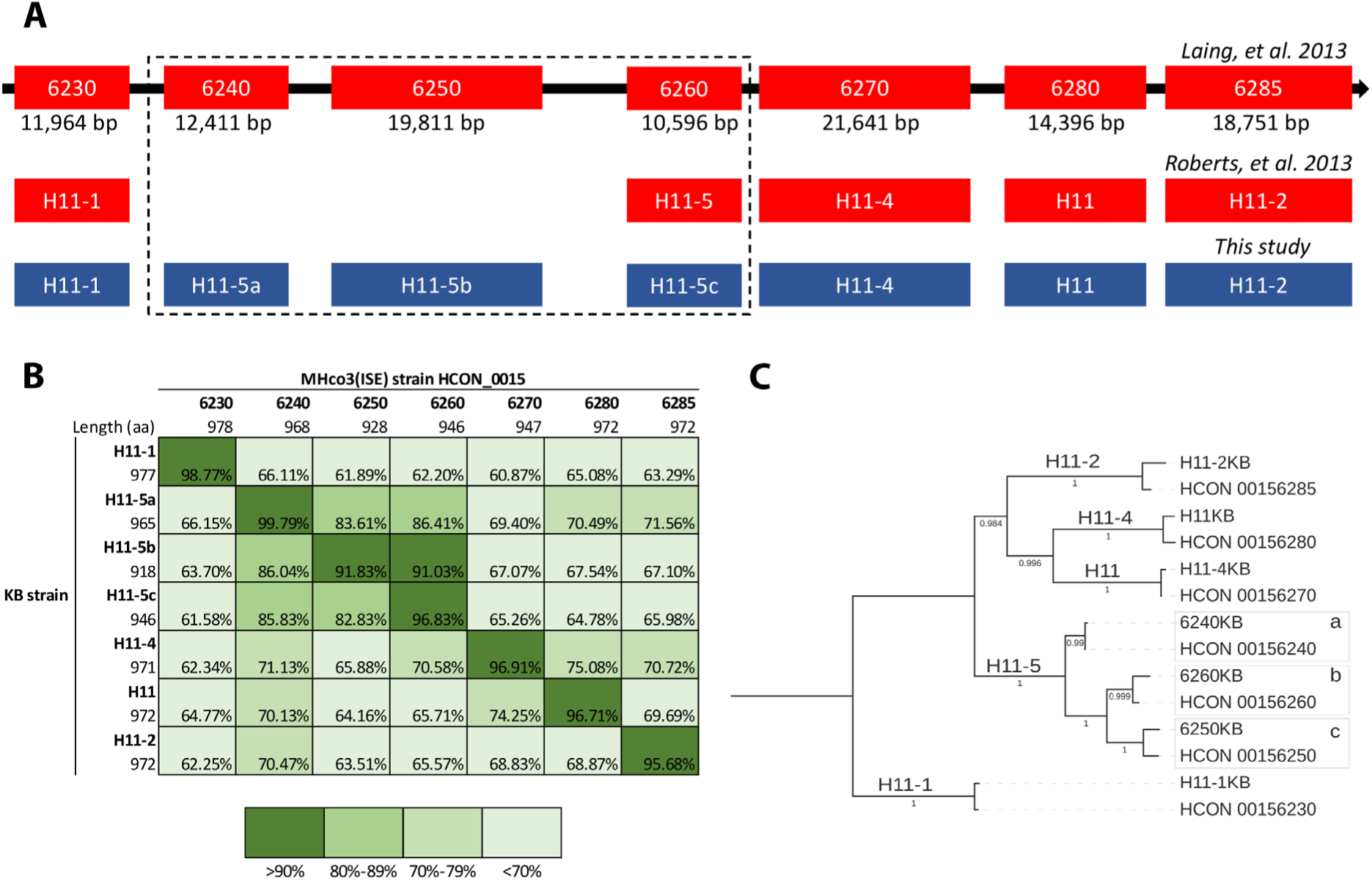
Identification of *h11* gene loci in the genome of *H. contortus* and sequence alignment of H11 isoforms. (A) Seven *h11* gene loci were determined on chromosome 5 of *H. contortus* genome (GenBank accession no. LS997566.1, 38,739,194 – 38,876,568). (B) Pairwise comparisons of H11 protein sequences determined in the KB and MHco3(ISE) strains. (C) A phylogenetic tree constructed with FastTree demonstrated the homology of H11 sequences used in this study.

### Recombinant baculoviruses were successfully constructed

The molecular engineering of *Haemonchus* antigens was performed *in silico*, involving the removal of predicted transmembrane domains (TMDs) from native protein sequences and the insertion of functional elements. These included a melittin signal peptide, a HisFLAG duo-affinity tag, and a thrombin cleavage site, all incorporated at the N-terminus of TMD-truncated antigen sequences. The protein sequences of the designed recombinant antigens are provided in the **S1 Text**. Based on protein sequences, codon-optimised DNA fragments encoding all seven isoforms of H11 (KB strain) were synthesised and subcloned into the pACEBac1 acceptor vector (**Fig 4**). To strengthen the potential efficacy of a multivalent vaccine, we also included an additional antigen as a complementary target - the apical gut membrane polyprotein GA1 (*H. contortus* US strain, GenBank AAB01192.1), another ‘hidden’ antigen of *H. contortus* previously proposed as a promising vaccine candidate [22].

**Fig 4.**
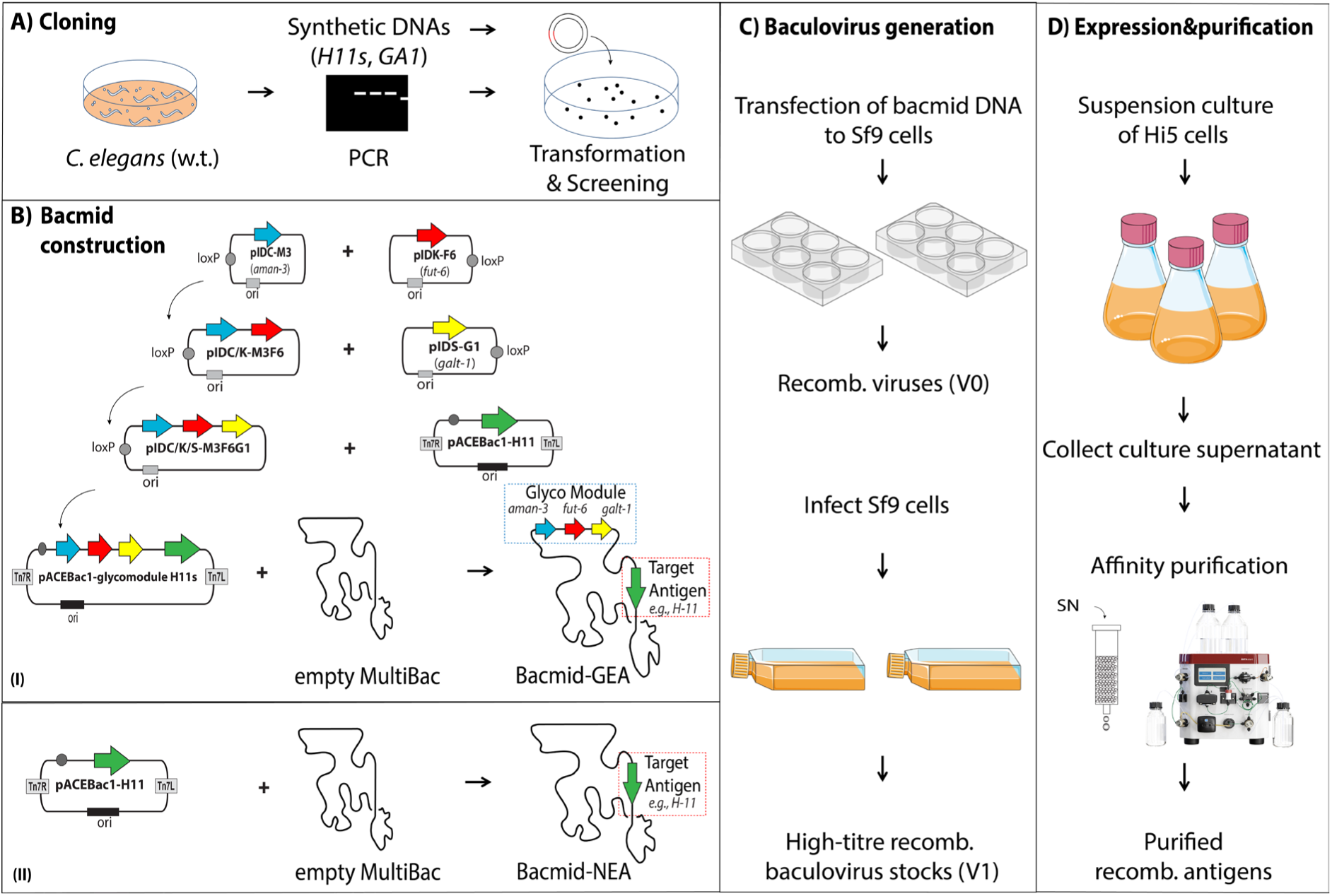
Schematic workflow of the preparation of glycoengineered recombinant antigens. (A) Molecular cloning of synthetic DNAs and glyco-genes from wild type *C. elegans* and (B) construction of bacmid DNAs containing antigen-encoding DNAs with or without the ‘glyco-module’. (C) Preparation of high-titre recombinant baculovirus stocks using Sf9 insect cells. (D) Recombinant expression and purification of soluble antigens of *H. contortus*.

DNA constructs obtained at each cloning step were verified by PCR and DNA sequencing. In parallel, coding sequences (CDSs) of the three corresponding *C. elegans* glyco-genes (*aman-3*, *fut-6* and *galt-1*) were first cloned using the MultiBac^TM^ kit into donor vectors separately (pIDC, pIDK or pIDS) and then integrated together to form a large donor vector that hosts all three CDSs as the glyco-module (**Fig 4B and Fig 5A**). Nanopore DNA sequencing data confirmed the successful integration of the glyco-module and one of the eight antigen-encoding genes (**S3 Fig**). At the end, in total 16 recombinant baculoviruses, either with or without the glyco-module, were successfully prepared; they were able to be propagated in Sf9 cells and to infect Hi5 cells, as judged by the appearance of YFP signals (**S4 Fig**).

**Fig 5.**
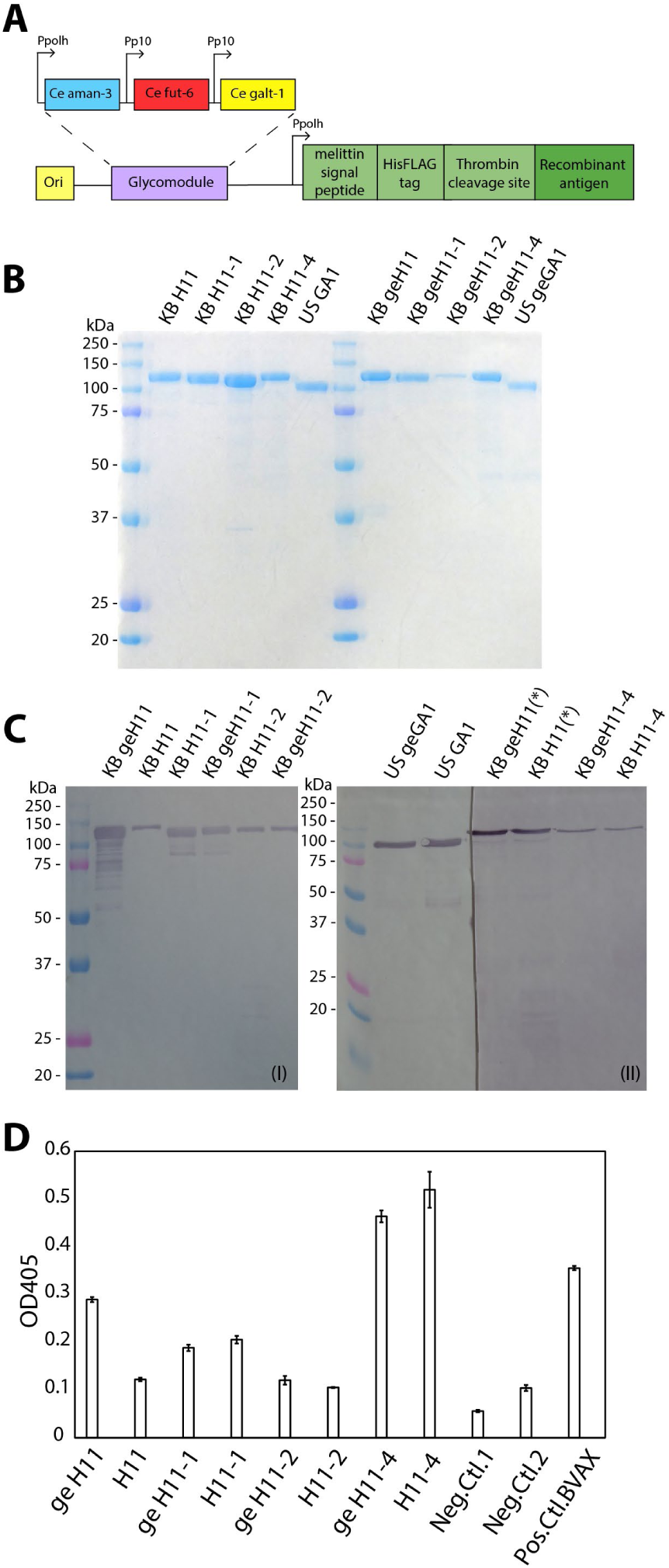
Analysis of recombinant H11s and GA1 antigens. (A) Schematic presentation of DNA constructs carrying the glyco-module and an engineered antigen-encoding gene controlled by separate promoters. (B) Coomassie blue staining of purified recombinant H11 and GA1 antigens. (C) Western blot assessment of purified recombinant H11 and GA1 antigens, using either a rabbit anti-H11 polyclonal IgG (I) or an anti-FLAG IgG monoclonal antibody (II); asterisks indicate antigens probed by both antibodies. (D) Aminopeptidase activity of recombinant H11 antigens at pH 7.0. Absorptions at optical density OD_450_ were measured after incubation for 24 hours at 37°C. Reaction mixtures containing either no substrate (Neg.Ctl.1) or no recombinant antigen (Neg.Ctl.2) were used as negative controls. Barbervax^®^ was used as a positive control (Pos.Ctl.BVAX). ge, glycoengineered; KB, an isolate from Katharinenhof in Bavaria, Southern Germany; US, an isolate from Beltsville maintained by the United States Department of Agriculture.

### Yield, purity and enzymatic activity of recombinant antigens

Purified recombinant antigens were quantified by Bradford protein assay. Five out of the eight designed antigens were successful expressed recombinantly with a satisfactory yield (1-71 mg/L); the expression of H11-5 isoforms (a, b and c) did not result in a sufficient yield. Among the five non-glycoengineered antigens (NEA), H11-1 (71 mg/L) and GA1 (12 mg/L) showed the highest level of antigen production, followed by H11 (7 mg/L), H11-2 and H11-4 (3 and 2 mg/L, respectively). For the glycoengineered antigens (GEA), geH11 (9 mg/L) showed the highest antigen yield, followed by geH11-1 and geGA1 (both 7 mg/L); geH11-2 and geH11-4 showed the lowest level of expression (1 and 2 mg/L, respectively). Moreover, antigen purity was assessed by Coomassie blue staining post SDS-PAGE. All recombinant proteins were detected as a single band with approximal molecular weights at 110 kDa (H11 isoforms) or 100 kDa (GA1) (**Fig 5B**). Purified antigens were also visualised by Western blot using two sets of antibody combinations (**Fig 5C I and II**). Using the glycoengineered H11-1 antigen as an example, the N-terminal affinity tag was removed by overnight incubation with thrombin and the tag-free H11-1 could be detected using the H11(16-aa motif)-specific polyclonal antibody (**S5 Fig**). In addition, protein sequences of all recombinant products were verified by LC-MS/MS analysis, which indicated high peptide coverages except for H11-5a (**S1 Table**).

After confirming the successful expression and purification of intact H11 isoforms, we next assessed the enzymatic activity of the recombinant proteins. As H11 aminopeptidases belong to the M1 peptidase family, *in vitro* assays were carried out using *p*-nitroanilide *(p*NP)-leucine as a substrate and zinc as a cofactor. Data demonstrated that all recombinant H11s, both in non-engineered and glycoengineered forms, were able to hydrolyse the substrate. Among the isoforms, H11-4 and geH11-4 displayed the highest activity towards the given substrate at pH 7 (**Fig 5D**). Enzymatic activities were also measured at pH 6 and pH 8 (**S6 Fig**), which indicated a similar activity pattern across pH conditions.

### Glycoengineered antigens possess nematode-type N-glycans

MALDI-TOF MS spectra of N-glycans of NEA and GEA showed that apart from many shared glycan compositions, primarily being oligo-/pauci-mannosidic glycans (Hex_2-9_HexNAc_2_) and fucosylated glycans (Hex_2-3_HexNAc_2-3_Fuc_1-2_), a major difference in glycoform between the two samples was the appearance of a new peak (*m/z* 1427.5) on the MS spectrum of GEA (**Table 1**, **Fig 6A**). This mass corresponded to a tri-fucosylated glycan structure (Hex_3_HexNAc_2_Fuc_3_), indicative of an altered glycosylation pattern due to the built-in glyco-module. This unique glycan composition was observed for each glycoengineered antigen examined individually, with a diverse abundance (**S7 Fig**). Separation of PA-glycans on RP-amide HPLC led to two distinct chromatograms (**Fig 6B**) and further structural analysis of glycans in HPLC fractions by MALDI TOF MS/MS revealed that the glycoengineered antigens carried a series of nematode-type N-glycan structures. Glycans of NEA displayed the typical insect-type structures carrying either a α1,3- or a α1,6-linked core fucose or both, as judged by key MS/MS fragments at m/z 446 and 592 (**Fig 7A-D and 7F**) or the loss of Y-ion (**Fig 7E**). In contrast, many fucosylated glycans of GEA were detected in late eluted fractions (>6.5 g.u.) and displayed *m/z* 608 and 754 MS/MS fragment ions, indicating the presence of the Galβ1,4Fuc epitope on those glycans [23]. The major tri-fucosylated glycan H3N2F3 of GEA (7.3 g.u.) also displayed such diagnostic ions and sequential losses of two terminal fucoses (**Fig 7L**). The other tri-fucosylated composition H2N2F3 (*m/z* 1265) represented two isomeric structures, as judged by distinct elution (6.6 and 7.5 g.u.) and fragmentation patterns (**Fig 7H and 7I**). PC-modified glycans were observed in both samples (**Table 1**) represented in minor portion. Anionic glycans were not detected in any of the examined samples.

**Fig 6.**
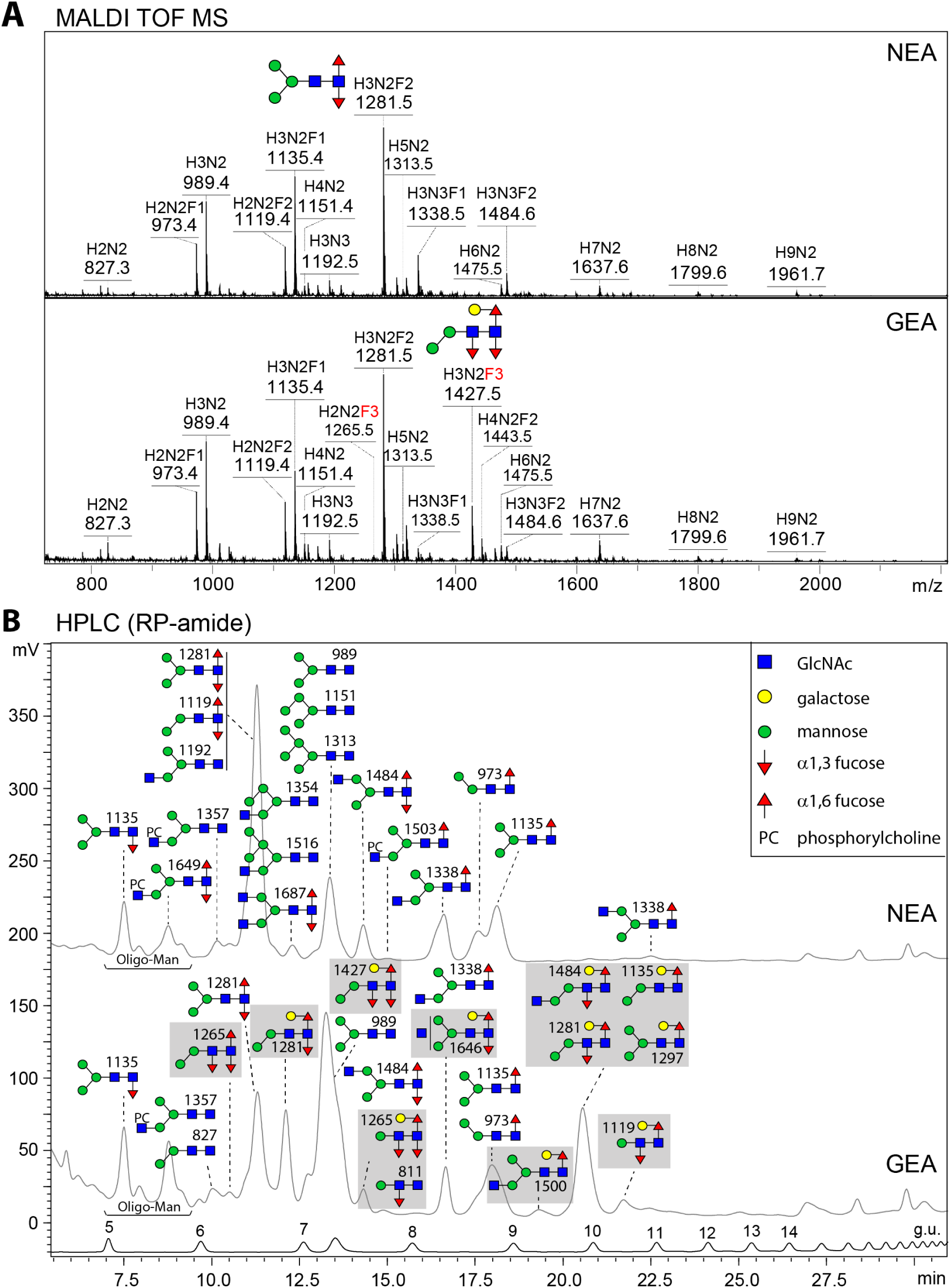
Distinctive N-glycoforms of non-glycoengineered antigens (NEA) and glycoengineered antigens (GEA). (A) MALDI-TOF MS spectra of 2-aminopyridine (PA)-labelled N-glycans released from the antigen cocktails, NEA and GEA. Tri-fucosylated glycan compositions are highlighted in red. (B) RP-amide HPLC chromatograms of PA-glycans of NEA and GEA. Peaks are annotated with major glycan structures (shown in SNFG format [57]), which were detected using MALDI TOF MS/MS. Nematode-type glycans are highlighted in gray boxes. H, hexose; N, N-acetyl-hexosamine; F, fucose; PC, phosphorylcholine; MS/MS, tandem mass spectrometry.

**Fig 7.**
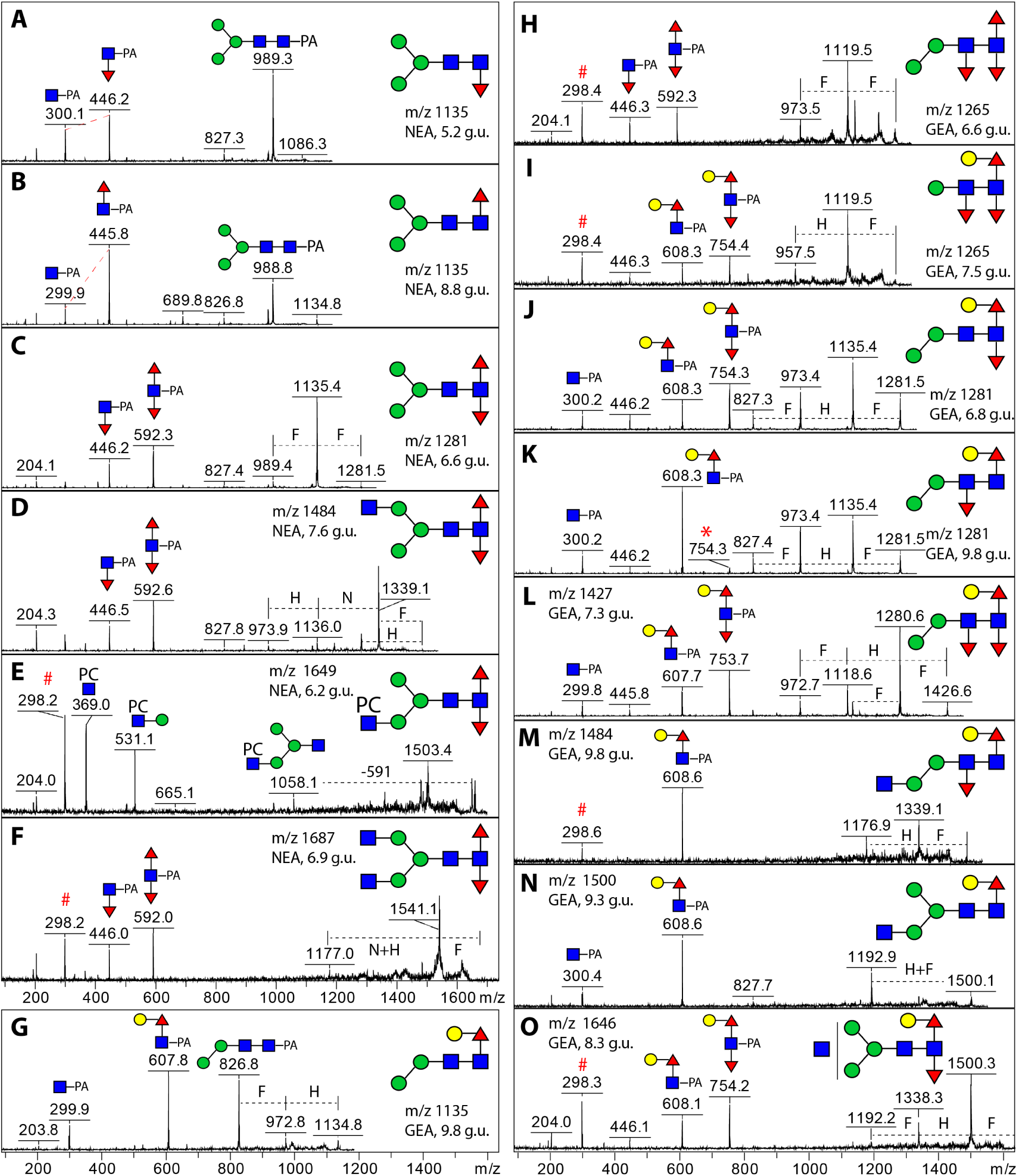
MALDI-TOF MS/MS spectra of HPLC-purified N-glycans. (A-F) Selected spectra of mono- and di-fucosylated N-glycans detected in NEA. (G-O) Nematode-type N-glycans were exclusively observed in GEA, possessing the Galβ1,4Fuc epitope and tri-fucosylated core. Masses (m/z) and glucose units (g.u.) are provided and key fragment ions are annotated. Asterisk (in K) indicates in-source rearrangement of core fucose; hashtags mark non-glycan noise signals.

**Table 1.** A comparison of N-glycans detected in the non-glycoengineered antigens (NEA) and glycoengineered antigens (GEA). Based on MALDI-TOF MS and MS/MS data, native and PA-labelled glycans before and after HPLC fractionation are summarised and their predicted compositions are provided. Checkmarks indicate the presence of a particular glycan mass in a sample. Key diagnostic MS/MS fragments of glycans containing one or more fucose (F) and/or phosphorylcholine (PC) residues are provided. Pauci-/oligomannosidic glycans are in bold. H, hexose; N, N-acetyl-hexosamine.

| Predicted composition | Observed m/z PA, [M+H]<sup>+</sup> | Calculated m/z PA, [M+H]<sup>+</sup> | NEA | GEA | Key diagnostic fragments |
| --- | --- | --- | --- | --- | --- |
| H1N2F1 | 811 | 811.35 |  | ✓ | trace 446 |
| **H2N2** | **827** | **827.34** |  | ✓ |  |
| H2N2F1 | 973 | 973.40 | ✓ | ✓ | 446 or 608 |
| **H3N2** | **989** | **989.39** | ✓ | ✓ |  |
| H2N3 | 1030 | 1030.42 |  | ✓ | 366/503/827 |
| H2N2F2 | 1119 | 1119.46 | ✓ | ✓ | 591 or 608 |
| H3N2F1 | 1135 | 1135.45 | ✓ | ✓ | 446 or 608 |
| **H4N2** | **1151** | **1151.45** | ✓ | ✓ |  |
| H3N3 | 1192 | 1192.47 | ✓ | ✓ |  |
| H2N2F3 | 1265 | 1265.51 |  | ✓ | 446/592 or 608/754 |
| H3N2F2 | 1281 | 1281.51 | ✓ | ✓ | 446/592 or 608/754 or 608/trace 754 |
| H4N2F1 | 1297 | 1297.50 |  | ✓ | 608 |
| **H5N2** | **1313** | **1313.50** | ✓ | ✓ |  |
| H3N3F1 | 1338 | 1338.53 | ✓ | ✓ | 446 |
| H4N3 | 1354 | 1354.53 | ✓ | ✓ | 366/503 |
| H3N3PC1 | 1357 | 1357.53 | ✓ | ✓ | 369/531 |
| H3N4 | 1395 | 1395.55 |  | ✓ | 503/1030 |
| H3N2F3 | 1427 | 1427.57 |  | ✓ | 608/754 |
| H4N2F2 | 1443 | 1443.56 | ✓ | ✓ | 592 |
| **H6N2** | **1475** | **1475.55** | ✓ | ✓ |  |
| H3N3F2 | 1484 | 1484.59 | ✓ | ✓ | 446/592 or 608/754 or 608 |
| H4N3F1 | 1500 | 1500.58 |  | ✓ | 608 |
| H3N3F1PC1 | 1503 | 1503.59 | ✓ | ✓ | 369/531/1058 |
| H5N3 | 1516 | 1516.57 |  | ✓ |  |
| H3N4F1 | 1541 | 1541.61 |  | ✓ |  |
| H3N5 | 1598 | 1598.63 | ✓ |  |  |
| **H7N2** | **1637** | **1637.60** | ✓ | ✓ |  |
| H4N3F2 | 1646 | 1646.64 |  | ✓ | 608/754 |
| H3N3F2PC1 | 1649 | 1649.64 | ✓ |  | 369/531/1058 |
| H3N4F2 | 1687 | 1687.66 | ✓ |  | 446/592/1541 |
| H3N4F1PC1 | 1706 | 1706.66 | ✓ | ✓ | 369, 531 |
| **H8N2** | **1799** | **1799.66** | ✓ | ✓ |  |
| **H9N2** | **1961** | **1961.71** | ✓ | ✓ |  |

### Recombinant antigens are capable of inducing cytokine secretion in vitro

To evaluate the immune response induced by NEA and GEA, their immunomodulatory effect was assessed on sheep peripheral blood mononuclear cells (PBMCs) *in vitro*. Supernatants were examined for chemokines/cytokines by Luminex assay (**Fig 8)**. Stimulation with both cocktails increased production of IL-1β, IL-6, IL-10, MIP-1α, MIP-1β, and TNF-α) at 48 and 72 hours of incubation compared to the negative control. Compared to ovine PBMC stimulated with antigens isolated from Barbervax (BVAX) in the same concentration, GEA, GEA(+) and NEA(+) induced higher levels of IL-1β, IL-10, MIP-1α and MIP-1β. Interestingly, the only cytokine for which BVAX induced a stronger response compared to the antigens prepared as part of this study was IL-6, while levels of IL-1β, IL-10, MIP-1α and MIP-1β induced by BVAX did not exceed those of the negative control. Quil A^®^, included as an adjuvant control, induced a marked cytokine response, underscoring its potent immunostimulatory properties. Con A, used as a positive technical control, stimulated the secretion of most chemokines/cytokines in a time dependent manner. For the remaining chemokines/cytokines included in the Luminex assay, either no induction was seen or they exhibited high variability, suggesting a lack of consistent biological relevance (**S8 Fig**).

**Fig 8.**
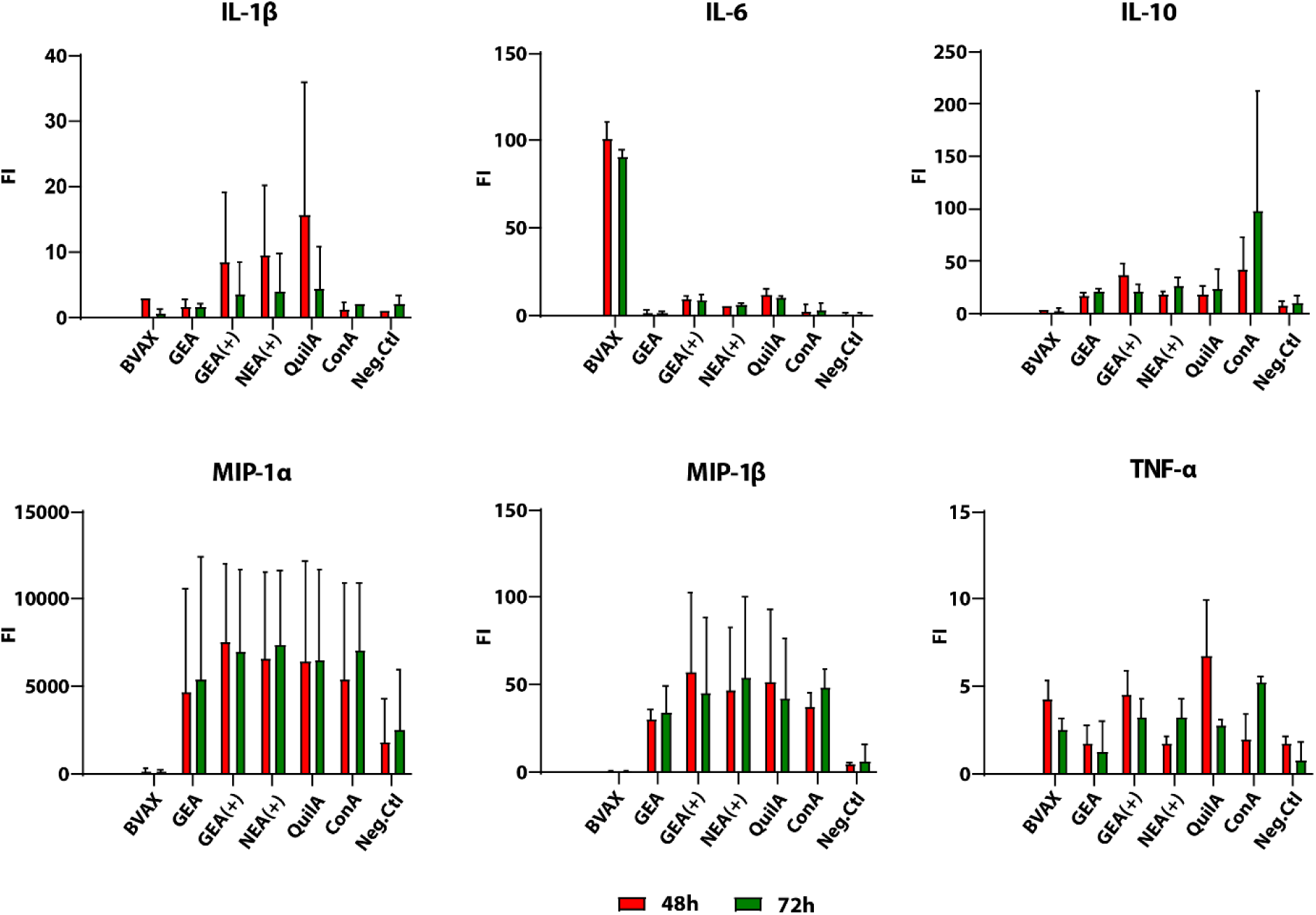
Biological activities of recombinant antigens. Cytokine secretion by ovine PBMCs was measured after 48 and 72 hours of stimulation with various antigen formulations using a Luminex Bio-Plex 200 system (Bio-Rad). Tested formulations included: BVAX (detergent-free antigens of Barbervax^®^); GEA, a glycoengineered antigen cocktail without adjuvant; GEA(+), a glycoengineered antigen cocktail with Quil A^®^ as adjuvant; and NEA(+), a non-engineered antigen cocktail with Quil A^®^. Concanavalin A (Con A) served as a technical control and unstimulated cells acted as the negative control. The data presented in GraphPad presented as mean fluorescence intensity (FI) of two animal replicates for six cytokines: IL-1β, IL-6, IL-10, MIP-1α, MIP-1β, and TNF-α. The FI values are displayed on a linear scale, providing a clear comparison of cytokine responses across different treatments. Error bars shown in the graphs represent the standard deviation (SD) of the data.

## Discussion

One of the major challenges in anti-parasitic vaccine development is the identification and subsequent production of effective antigens in a controlled and reproducible manner. In this study, we report the expression pattern of *H. contortus* H11 aminopeptidases and present a novel method for producing these helminth antigens in a soluble and properly glycosylated form using a glycoengineered baculovirus expression system. Following expression and purification, these recombinant proteins exhibited clear immunogenicity in host immune cells *in vitro*, making them promising vaccine candidates for efficacy evaluation in animals.

The native *H. contortus* H11 aminopeptidases are membrane-bound glycoproteins of 110 kDa in size which can be purified from the intestinal fraction of adult parasites using lectin and ion-exchange chromatography [24,25]. H11 expression is speculated to begin at late developmental stages, as indicated by a stage-specific expression pattern of *h11* genes with significantly higher transcription levels in adult worms compared to L3 larvae [9]. However, direct evidence at the protein level was still missing. In this study, we closed this gap by generating H11-specific polyclonal IgG antibodies targeting conserved peptide motifs shared by different H11 isoforms. Using an antibody specific to a surface-exposed motif, we detected intact H11 in adult worms but not in L3 larvae. This confirms a stage-specific expression pattern of H11 at the protein level, consistent with previously published LC-MS/MS-based somatic proteome data [26].

By far, various H11 sequences generated from different *H. contortus* isolates have been deposited in GenBank. Sequence alignment suggested that these fall into five major clusters, namely H11, H11-1, H11-2, H11-4 and H11-5 [9,27]. As a thorough understanding of H11 isoforms was necessary for the production of recombinant proteins, we re-assessed the last genome build of the *H. contortus* (MHco3(ISE) strain) [21], and identified seven gene loci encoding distinct H11 isoforms on chromosome 5. Sequence alignment indicated that three of the isoforms (6240, 6250 and 6260) shared a higher degree of homology (>83%) compared to the other isoforms, thereby belonging to the H11-5 branch (designated H11-5a, H11-5b, and H11-5c in this study). Our experimental data confirmed that all seven isoforms were transcribed in the adult *H. contortus* (KB strain), raising the question of whether each isoform is functionally active *in vivo*. The presence of multiple H11 isoforms is thought to provide redundancy of aminopeptidase activity, which is crucial for the parasite to efficiently process blood meals [28]. If all the seven isoforms are expressed in adult parasites, an effective vaccine should ideally target all or most of them to ensure broad protection.

To overcome this potential limitation of previous studies, in which usually only a single H11 isoform or pairwise combinations were targeted [9–11], we sought to produce all seven isoforms to be included in the envisaged vaccine cocktail. In addition, we included the polyprotein GA1 antigen in our envisaged cocktail due to the following considerations: GA1 is an immunogenic antigen expressed in the intestine of adult *H. contortus* [22], and, similar to H11, it is abundant in the intestinal protein extracts [29]. Moreover, GA1 is an N-glycoprotein which may carry similar glycan modifications as the native H11 antigens. Although recombinant baculoviruses for expressing all eight desired antigens were generated successfully, it remained difficult to produce the H11-5 isoforms. Nevertheless, five out the eight desired antigens could be recombinantly produced in sufficient yield and purity.

The glycoengineering approach employed in this study offered a solution for generating more effective H11-based recombinant vaccines against *H. contortus,* as the engineered antigens possessed glycan epitopes identical to those found in the native parasite antigens. This concept has been beneficial for improving the efficacy of vaccine antigens in another study, in which a glycoengineered ASP-1 antigen, recombinantly produced in *Nicotiana benthamiana*, induced an improved immune protection against the parasitic nematode *Ostertagia ostertagi* [30]. Tri-fucosylated N-glycans, originally found in adult *H. contortus* and on purified native H11 [12], have been reported in a few nematode species, including *O. dentatum* [31] and the free-living *C. elegans* and *Pristionchus pacificus* [32,33]. Although the biological role of such core modified glycans remains unclear, a recent study on *Schistosoma* glycans indicated that non-mammalian type core modifications (β1,2 xylose and α1,3 fucose) were highly immunogenic to the host and represented as ligands of the protective IgG antibodies raised during infection [34].

In this project, we demonstrated how *H. contortus* antigens carrying tri-fucosylated nematode-type glycans could be produced using glycoengineered insect cells. We chose *Trichoplusia ni*-derived Hi5 cell line instead of *Spodoptera frugiperda*-derived cell lines (Sf9 or Sf21) as expression host, because Hi5 cells are capable of producing N-glycosylated proteins with abundant core di-fucosylated glycans [16,17], a foundation for incorporating nematode glycoenzymes into the Hi5’s native glycan biosynthetic pathways. Baculoviruses carrying the complete open reading frames of three *C. elegans* genes (*aman-3*, *fut-6* and *galt-1*) were used to infect Hi5 cells, which led to the expression of three Golgi-localised glycoenzymes, AMAN-3, FUT-6 and GALT-1. The former two were required for introducing α1,3-linked fucose on the distal GlcNAc residue of N-glycans [18], whereas GALT-1 galactosylated the core α1,6-linked fucose [20]. Glycan analysis using HPLC and mass spectrometry provided evidence on the presence of nematode-type glycan structures, including tri-fucosylated structures and a set of mono-/di-fucosylated structures carrying Gal β1,4Fuc epitope, which only appeared on the glycoengineered forms. Moreover, to ensure proper glycosylation and folding of the recombinant products, we engineered the naturally membrane-bound H11 and GA1 antigens for extracellular production, including exclusion of their transmembrane domains and addition of a signal peptide and cleavable affinity-tags. To enhance expression efficiency, antigen-encoding DNAs were codon-optimised for Hi5 cells and chemically synthesised. As demonstrated in this study, the truncated recombinant antigens (except for H11-5) could be obtained in soluble form from cell culture supernatant. The H11 isoforms were enzymatically active between pH 5 and 8, strongly indicating correct protein folding.

Using the adjuvanted antigen cocktails, we were able to show that GEA(+) and NEA(+) are capable of eliciting enhanced immune responses compared to BVAX. This indicates that Hi5 cell-derived recombinant antigens are biologically active *in vitro* and possess ligands that can stimulate ovine immune cells to secrete a specific set of cytokines. *In vitro* cytokine profiling at 48- and 72-hour post-stimulation revealed distinct immune responses to the tested antigens, highlighting the impact of glycoengineering and adjuvant selection. Both NEA(+) and GEA(+) elicited a robust pro-inflammatory and chemoattractant response that was not observed in the supernatants of cells exposed to BVAX, and this response was sustained for the whole period. GEA, the antigen cocktail without the adjuvant, exhibited diminished cytokine production, underscoring the adjuvant’s essential role. Even though the assays were only done with PBMCs derived from two sheep, GEA still induced higher levels of IL-10, MIP-1α and MIP-1β compared to BVAX at 48 hours, and similar but prolonged IL-1β secretion. Among the tested cytokines IL-1β, a potent mediator of inflammation and neutrophil recruitment [35–37], plays a critical role in host defence [38,39], while MIP-1α and MIP-1β further enhance immune cell recruitment to infection sites [40]. IL-10 is a multifunctional cytokine that limits inflammatory responses and regulates the differentiation and function of various immune cells, including regulatory T cells, playing a key role in immune homeostasis and tolerance [41]. In the case of BVAX, in contrast to our antigen cocktails, a strong IL-6 response was observed, indicative of a potent but potentially imbalanced innate immune activation; IL-6, a pleiotropic cytokine, modulates immune responses through effects on lymphocyte differentiation and macrophage recruitment [42]. High IL-6 and reduced IL-10 levels, as seen with BVAX, suggest persistent inflammation and potential immune dysregulation; while this has not been reported for parasitic infection, it has been described for other infectious diseases on mucosal surfaces, such as mastitis in cattle [43]. This imbalance may impair inflammation resolution and disrupt immune homeostasis. Both technical controls, Quil A^®^ and Con A elicited either the expected inflammatory response, in case of Quil A^®^ or, stimulated IL-10 production, consistent with non-specific Th2 T-cell activation, in case of Con A [44,45]. In conclusion, GEA(+) demonstrated superior immune activation, attributed to its engineered glycoforms in combination with the adjuvant, highlighting the potential of this approach for formulating a highly bioactive and effective recombinant vaccine against *H. contortus*.

The glycoengineered H11 and GA1 antigens, carrying nematode-type N-glycan modifications, retained their enzymatic activity and stimulated cytokine secretion from ovine PBMCs *in vitro*, demonstrating the potential of this approach for producing vaccine antigens against *H. contortus*. Antigens produced using this approach were tested by the research team in a controlled vaccination trial in sheep, delivering promising results (Sajovitz-Grohmann *et al.* 2025). This work demonstrated the feasibility of modifying the native glycosylation machinery of Hi5 insect cells and introducing foreign glycan biosynthetic pathways to enable essential immunogenic glycan epitopes on recombinant products. As a robust expression system, Hi5 insect cells can be adapted for other engineering strategies and can be used for producing a broad spectrum of vaccine antigens, not only for anti-*H. contortus* vaccination but also for tackling other metazoan pathogens, such as *Schistosoma* and *Fasciola*.

## Materials and methods

### Preparation of antigen-specific polyclonal antibodies

Two H11-specific polyclonal antibodies (pAbs) were custom-produced by Eurogentec (Seraing, Belgium). To identify conserved peptide motifs, six major H11 antigen sequences (GenBank accession nos. CAB57357.1, CDJ83822.1, AGT9010.1, CAB57358.1, CAC39009.1, Q10737.2) were aligned using MAFFT [46] and visualised using UGENE [47]. Structural models of these sequences were generated using AlphaFold [48] and visualised in PyMOL [49]. Two polypeptides (C+GAMENWGLITYRE, 14 aa and C+EPEKYRHPT/KYGFKWD, 16 aa), each exhibiting identical or highly similar sequences across all isoforms, were chemically synthesised, conjugated through the N-terminal cystine to ovalbumin, and used as immunogens to generate polyclonal IgG antibodies in rabbits.

### Cultivation of E. coli cells

Different *E. coli* strains were used for the preparation of DNA constructs. NEB5α cells were purchased from New England Biolabs (Ipswich, MA, USA); chemical competent cells One Shot™ PIR1 were purchased from Fisher Scientific (Fair Lawn, New Jersey, USA) and DH10EMBacY from Geneva Biotech (Geneva, Switzerland). In brief, *E. coli* cells were cultivated in Lysogeny-Broth medium (Carl Roth, Karlsruhe, Germany) supplied with antibiotics of choice, including kanamycin (20 µg/ml), streptomycin (50 µg/ml), chloramphenicol (25 µg/ml), and tetracycline (10 µg/ml) purchased from Sigma-Aldrich (St. Louis, MO, USA) and gentamycin (10 µg/ml) from Fisher Scientific.

### Preparation of nematode cDNAs

Isolation of RNA from the N2 wild type *Caenorhabditis elegans* and adult *Haemonchus contortus* (KB strain, an isolate from Kathrinenhof in Bavaria, Germany) was performed according to the QIAamp^®^ RNA Blood Mini kit (QIAGEN, Hilden, Germany) protocol. For the purification of total RNA from tissue, worms were placed in a glass homogeniser (pre-treated with DEPC-treated water and autoclaved), and 600 µL of Buffer RLT (supplied with β-mercaptoethanol) was added. The tissue was then disrupted until it appeared homogeneous. DNase treatment was performed to avoid genomic DNA contamination. Purified RNA was quantified using a Nanodrop^®^ 2000 spectrophotometer and converted to cDNA using GoScript™ Reverse Transcriptase (Promega, Madison, WI, USA). Briefly, 4 µl of RNA was mixed with 1 µl of oligo(dT)_15_ primers (Promega), heated at 70°C for 5 min and immediately chilled on ice. In a separate tube, the reverse transcription mix was prepared on ice by combining 4 µl of GoScript™ 5x Reaction Buffer, 1.2 µl of 25 mM magnesium chloride, 1 µl of 10 mM (each dNTP) PCR nucleotide mix, 1 µl of GoScript™ reverse transcriptase, and nuclease free water to a final volume of 15 µl. Both mixes were combined, and cDNA synthesis was performed in a thermocycler as follows: annealing at 25°C for 5 min, extension at 24°C for 1 h, followed by inactivation of the enzyme at 70°C for 15 min. Obtained cDNAs were stored at -80°C prior to the PCR amplification of specific genes.

### PCR amplification of H11 isoforms and DNA sequencing

Predicted H11 encoding genes were PCR amplified with GoTaq® G2 Hot Start Green Master Mix (Promega, Madison, WI, USA) using the primers and optimised annealing temperatures specified in the **S1 Text**. Briefly, 10 µl reactions were set up containing GoTaq^®^ G2 Hot Start Green Master Mix, 0.2 µM of each primer, the appropriate amount of nuclease free water and 0.5 µl of 1:10 diluted cDNA as template. The thermocycler was set at 2 min of initial denaturation, followed by 30-40 cycles of 30 s at 95°C (denaturation), 30 s at the optimised annealing temperature (specified in **S1 Text**), and 3.5 min at 72°C (elongation), completed by a final 10 min elongation step. PCR products were verified by gel electrophoresis and either sequenced directly or subsequently cloned into a pCR4TOPO vector by TA cloning (Invitrogen TOPO™ TA Cloning Kit; Fisher Scientific, Fair Lawn, New Jersey, USA) and transformed into NEB5α cells. Positive transformants were identified by blue-white screening, followed by PCR screening to verify full length inserts with primers T3 (ATTAACCCTCACTAAAGGGA) and T7 (TAATACGACTCACTATAGGG) provided in the cloning kit. Selected clones were cultured as described above, the plasmids isolated, and the inserts sequenced with primers M13-21F (TGTAAAACGACGGCCAGT) and M13-rev2 (GAGTTAGCTCACTCATTAGG). Additionally, to rule out any point mutations potentially introduced by *Taq* polymerase, H11 encoding genes were PCR amplified from cDNA using Q5^®^ High-Fidelity DNA Polymerase (New England Biolabs) and sequenced directly. Sanger DNA sequencing of PCR products and DNA constructs was performed at LGC Genomics (Berlin, Germany). Additional sequencing primers were designed to cover the full-length sequences of H11 encoding genes (provided in the **S1 Text**).

### Protein sequence analysis and antigen design

Assembly of contigs was carried out using a SnapGene software (GSL Biotech LLC, San Diego, US) using the built-in assembly tool. When necessary, DNA sequences were inspected using raw sequencing chromatograms and mis-interpreted nucleotides were manually corrected. The obtained CDS of *h11* isoforms were compared with the chromosome 5 genomic data of *H. contortus* (GenBank: LS997566.1) by performing BLASTN searches. Deduced protein sequences (KB strain), translated by SnapGene, were compared with published H11 sequences (MHco3(ISE) strain) by performing BLASTP searches. Sequence alignment was performed using MAFFT [46] and a phylogenetic tree was constructed with FastTree [50], using Approximately Maximum Likelihood methods, and visualised with iTOL [51]. N-glycosylation sites were predicted using NetNGlyc - 1.0 server [52].

Antigens to be recombinantly expressed (H11s and GA1) were designed as follows: 1), transmembrane helixes (excluded) were predicted using the TMHMM 2.0 server [53]; 2), the truncated antigen sequences were codon-optimised using the GenSmart™ Codon Optimisation tool (expression host: High Five cells; GenScript, Leiden, NL); 3), a DNA sequence, encoding a melittin signal peptide and a HisFLAG duo-affinity tag coupled with a thrombin cleavage site to the upper stream of codon-optimised antigen sequences, was added. Synthetic DNAs (GenScript) were used for molecular cloning.

### Construction of recombinant baculoviruses

Gibson assembly approach was employed for all molecular cloning work using a NEBuilder^®^ HiFi DNA assembly cloning kit (New England Biolabs) and specific primers (**S1 Text**). A MultiBac™ cloning kit (Geneva Biotech, Geneva, Switzerland) was used to prepare recombinant baculovirus following the user manual (version 8.3). Synthetic antigen-coding genes were subcloned to the pACEBac1 acceptor vector and transformed to competent NEB5α cells, whereas *C. elegans* glyco-genes (*aman-3*, *fut-6* and *galt-1*) were subcloned to donor vectors (pIDC/K/S) and transformed to competent PIR1 cells (**Fig 4A**). Post antibiotic selection and PCR screening, plasmid DNAs were prepared and sequenced. Construction of the glyco-module (all three glyco-genes) was achieved by combining pIDC, pIDK and pIDS constructs via Cre-LoxP recombination. For expressing glycoengineered antigens, pACEBac1 constructs, each hosting one antigen-coding gene, were combined with a construct hosting the glyco-module. Resulting plasmids were verified by PCR and Nanopore sequencing (Microsynth AG, Balgach, Switzerland). Subsequently, the plasmids were transformed to DH10EMBacY competent cells and positive bacmids were obtained post blue-white screening and were PCR verified using specific primers in combination (**Fig 4B**).

### Insect cell culture, expression and purification of recombinant antigens

Sf9 insect cell line was obtained from ATCC (CRL-1711) via LGC Standards (Wesel, North Rhine-Westphalia, Germany) and the High Five^TM^ cell line (Hi5) was ordered from Fisher Scientific (Fair Lawn, New Jersey, USA). Both cell lines were adapted to grow in HyClone^TM^ SFM4Insect cell culture medium; for Sf9 cells, 2% Gibco foetal bovine serum was added. Cells were maintained in T-flasks at 27°C without CO_2_ and passaged twice a week when they reached ∼90% confluency. The Sf9 cell line, derived from *Spodoptera frugiperda*, was used for DNA transfection and viral propagation, whereas the Hi5 cell line, derived from the ovarian cells of *Trichoplusia ni*, was used for antigen expression.

Transfection of bacmid DNAs to Sf9 insect cells was carried out in 6-well cell culture plates (**Fig 4C**), using the FuGENE^®^ HD transfection reagent (Promega, Madison, WI, USA). For each construct, 1 µg of bacmid DNA was mixed with 10 µl of transfection reagent and added to Sf9 cells. Typically, recombinant baculoviruses (V_0_) were ready for harvesting 4 to 6 days post transfection, when YFP fluorescence could be detected in most of the cells. Subsequently high-titre viruses (V_1_) were obtained by infecting Sf9 cells with V_0_ viruses either in T75 cell culture flasks or in 100 ml Erlenmeyer shake flasks. Extracellular expression of *Haemonchus* antigens in Hi5 cells was initially carried out in 6-well plates and cell culture supernatants were examined by Western blotting using a custom-made H11(16 aa motif)-specific polyclonal antibody.

Large-scale suspension culture of Hi5 insect cells was carried out in 1 litre Erlenmeyer cell culture flasks at 27°C on a horizontal shaking platform (80 rpm) (**Fig 4D**). Typically, a 200 ml culture with a density of 1×10^6^ cells/ml was infected with 4 ml V_1_ viral stock (1:50) and harvested 3 days post infection. Cells and debris were removed by centrifugation and filtration (0.22 µm) prior to concentration and buffer-exchange steps using an ultrafiltration device (Amicon® Stirred Cell, Millipore) equipped with a 50 kDa molecular weight cut-off (MWCO) disc. Concentrated cell culture supernatants in binding buffer (20 mM sodium phosphate, 0.5 M NaCl, 20 mM imidazole, pH 7.4) were subject to affinity purification using an ÄKTA^TM^ Start protein purifier equipped with a fraction collector and a 1 ml prepacked HisTrap™ High Performance column (GE healthcare, Chicago, Illinois, USA). A 20-min linear gradient of elution buffer (20 mM sodium phosphate, 0.5 M NaCl, 500 mM imidazole, pH 7.4) was used to elute His-tagged proteins from the column. Purified recombinant antigens were desalted using Amicon^®^ Ultra-15 centrifugal devices, quantified using a Pierce™ Bradford Plus protein assay kit (Thermo Fisher Scientific) and stored in PBS at 4°C.

### Native antigens and recombinant antigen cocktails

Detergent-free native antigens were extracted from Barbervax^®^ (imported via Merlin Vet, Kelso, UK). 40 ml of Barbervax^®^ was concentrated and buffer-exchanged in PBS using Amicon^®^ Ultra-15 centrifugal filter units with a 3 kDa MWCO. Detergent in the sample was removed using a DetergentOUT™ GB-S10-3000 column (G-Biosciences) purchased from VWR (Vienna, Austria). The concentration of Barbervax^®^-derived native antigens (BVAX) was quantified using a Pierce™ BCA Protein Assay Kit (Thermo Fisher Scientific).

Adult *H. contortus* worms, resuspended in PBS supplemented with 1% Thesit^®^, 0.02% sodium azide, and a 1% protease inhibitor cocktail (Sigma-Aldrich), were transferred to a sterile glass homogeniser. Tissue disruption was performed on ice until a visually uniform homogenate was obtained. After centrifugation at 20,000 × *g* for 3 min at 4°C, supernatant containing soluble worm antigens was collected and stored at -20°C prior to subsequent assays. Homogenisation of third-stage larvae (L3) was performed using a porcelain mortar and pestle in liquid nitrogen. Supernatant of L3 worm lysate (in PBS supplemented with 1% Triton X-100, 1% protease inhibitor cocktail) was obtained by centrifugation at 15,000 × *g* for 20 min at 4°C. Protein concentrations were determined using the Pierce™ Bradford Plus protein assay kit (Thermo Fisher Scientific).

To assess the impact of different glycosylation patterns on immune response, two antigen cocktails were prepared, containing either non-glycoengineered antigens (NEA) or glycoengineered antigens (GEA). Each cocktail comprised equal amounts of five recombinant antigens (KB H11, KB H11-1, KB H11-2, KB H11-4, and US GA1), with the final protein concentration adjusted to 100 µg/ml. To formulate adjuvanted vaccines, Quil A® saponin was added to each antigen pool at a final concentration of 1 mg/ml.

### SDS-PAGE and Western blotting

To validate custom antibodies and verify the recombinant antigens, protein samples were dissolved in SDS-loading dye, denatured at 95°C and loaded on 12% SDS-PAGE gels. Precision Plus Protein™ Dual Color Standards (Bio-Rad Laboratories GmbH, Vienna, Austria) was used as protein ladder. Post electrophoresis at 200 V, proteins were either stained with ReadyBlue^®^ (Sigma-Aldrich) or transferred onto nitrocellulose membranes using a TransBlot Turbo™ transfer system (Bio-Rad). Immunodetection was performed using various antibodies, including rabbit anti-H11 (anti-H11 16 aa motif and anti-H11 14 aa motif, 1:1000), or anti-FLAG IgG monoclonal antibody (1:10,000, Sigma-Aldrich). Alkaline phosphatase-conjugated secondary antibodies, including goat anti-rabbit IgG (1:10,000, Sigma-Aldrich) and anti-mouse IgG (Fc-specific, 1:10,000, Sigma-Aldrich), were used for detection, with SIGMAFAST™ BCIP®/NBT as the substrate for colour development.

### Glycomic analysis of recombinant antigens

50 µg of GEA and NEA antigen cocktails were dissolved in 8 M urea and trypsinised overnight at 37°C. Tryptic peptides were purified using hand packed C18 cartridges prior to overnight deglycosylation by PNGase A (New England Biolabs). Released N-glycans were purified using two tandem cartridges, one filled with Dowex (50W×8) and C18, and another one with nPGC (Supelclean ENVI-Carb™ SPE, Sigma-Aldrich), following a protocol described by *Hykollari at al* [54]. Additionally, N-glycans were released from 25 µg of individual antigens following the same steps and examined subsequently. Lyophilised native GEA and NEA glycans were labelled with 2-aminopyridine (PA) to introduce a fluorescent tag at the reducing ends and the excess linker was removed by gel filtration (Sephadex^®^ G15, 0.5% acetic acid as solvent) as previously described [33]. PA-glycans were fractionated on an RP-amide column by a Shimadzu Nexera UPLC system equipped with an RF 20AXS fluorescence detector (excitation/emission: 320 nm/400 nm). Pooled and HPLC-fractionated N-glycans were analysed by MALDI TOF MS/MS (Bruker rapifleX, Bremen, Germany) in positive and negative ion mode using 6-aza-2-thiothymine as a matrix. MS and MS/MS spectra were manually interpreted based on the mass and fragmentation pattern. Glycan structures were proposed based on their key MS/MS fragment ions and glucose units (g.u.) in comparison with previously published glycomic data of Hi5 cells [17], *C. elegans* [55,18] and *O. dentatum* [31].

### LC-MS/MS

Aliquots of tryptic peptides were resuspended in 0.1% trifluoroacetic acid (TFA) and subjected to sequence identification using a Q Exactive Orbitrap LC-MS/MS system (Thermo Fisher Scientific). Briefly, peptides were separated on a nano-HPLC Ultimate 3000 RSLC system (Dionex) equipped with a 25 cm Acclaim PepMap C18 column (75 µm inner diameter, 2 µm particle size, and 100 Å pore size). Full-scan mass spectra were acquired in the ultrahigh-field Orbitrap mass analyser over the m/z range of 350–2000, with a resolution of 60,000. The maximum injection time (MIT) was set to 50 ms, and the automatic gain control (AGC) target was set to 3 × 10^6^. The ten most intense ions were selected for MS/MS fragmentation via high-energy collision dissociation (HCD) in the Orbitrap. Data were searched against a recombinant antigen sequence database using Proteome Discoverer software (version 2.4.1.15; Thermo Fisher Scientific). Further details are provided in the **S1 Table**.

### Enzymatic activity of H11 antigens

Aminopeptidase activity of recombinant H11 proteins expressed in Hi5 insect cells was verified following a standard protocol with small modification [9]. Purified recombinant H11 antigens (1 µg/well) were incubated in a 96-well plate with 0.5 mM L-Leucine-*p*-nitroanilide (Sigma-Aldrich) in the presence of 2 mM ZnCl_2_ in McIlvaine buffer, pH 7. Reactions were carried out in triplicate at 37°C. An equivalent amount of native antigens from Barbervax^®^ was included as positive control. After overnight incubation, reactions were stopped by adding 200 µl of 0.4 M glycine-NaOH buffer and OD_405_ was measured using a microplate reader (Filter Max F5, Molecular Devices). Enzymatic activity was also measured using McIlvaine buffer at pH 5 and pH 8.

### Ovine PBMC isolation and cytokine analysis

In a final experiment, the NEA and GEA cocktails were examined for their ability to induce an immune response in ovine cells. Ovine peripheral blood mononuclear cells (PBMCs) were isolated using peripheral blood obtained by venipuncture of the jugular vein of two clinically healthy, nematode-free, 5-month-old crossbreed (Jura x Lacaune) male lambs. Blood sampling was approved by the Ethics and Animal Welfare Committee of the University of Veterinary Medicine, Vienna in accordance with the University’s guidelines for Good Scientific Practice and authorised by the Austrian Federal Ministry of Education, Science and Research (ref BMBWF 2024-0.766.710) in accordance with current legislation. 100 ml of venous whole blood was aseptically collected and mixed with 50 IU/ml of heparin in a sterile measuring flask (Gilvasan Pharma GmbH, Vienna, Austria). PBMC isolation was carried out in a biosafety cabinet immediately after blood collection. First, 100 ml of whole blood was diluted in a 1:1 ratio with RPMI1640 medium (Gibco, Life Technologies, Austria) supplemented with 10% heat-inactivated foetal calf serum (Gibco, Life Technologies, Austria), and 1% penicillin, streptomycin, and amphotericin (Sigma-Aldrich, Germany, complete medium). PBMCs were isolated from blood using Lymphoprep™ density gradient medium (1.077 g/ml, STEMCELL Technologies, Germany) following the previously established protocol and optimised for our usage [56]. PBMCs were counted in trypan blue solution, shown to have > 95% viability and seeded into 6-well cell culture plates with a density of 1 × 10⁶ cells per well.

After 24-hour incubation at 37°C with 5% CO_2_, recombinant antigen cocktails GEA(+) and NEA(+) (1 µg/ml, supplemented with Quil A^®^ adjuvant), GEA (1 µg/ml, without adjuvant), Quil A^®^ adjuvant (10 µg/ml), native antigens of Barbervax^®^ (1 µg/ml), and Concanavalin A (Con A, 10 µg/ml) diluted in RPMI medium were added to the cells to stimulate cytokine production. After 48h and 72h incubation, cell culture supernatants were collected by centrifugation and kept at -20°C prior to cytokine assays.

Cytokine expression levels were analysed using an ovine cytokine/chemokine panel with a mixture of 14 probes (Merck KGaA, Vienna, Austria). According to the manufacturer’s instructions, following overnight incubation and washing steps, beads were analysed using a Luminex Bio-Plex 200 instrument (Bio-Rad). Median fluorescent intensities (FI) of the analytes were exported from the raw data to Excel. Data analysis was performed using GraphPad Prism 10 (GraphPad Software, LLC., USA).

## Supporting Information

S1 Text. Supplementary methods and results.

S1 Fig. Sequence and 3D-structure alignment of H11 antigens.

S2 Fig. Western blot analysis of protein samples using a rabbit anti-H11 polyclonal antibody.

S3 Fig. An overview of eight DNA constructs carrying the glyco-module (*aman-3*, *fut-6*, *galt-1*) and different *H. contortus* antigen encoding genes.

S4 Fig. Microscopic images of baculovirus-infected Sf9 cells.

S5 Fig. Western blot assay of recombinant antigens treated with or without thrombin.

S6 Fig. Aminopeptidase activity of recombinant glycoengineered (ge) H11 antigens at different pH conditions.

S7 Fig. MALDI-TOF MS spectra of native N-glycans released from individual recombinant antigens.

S8 Fig. Cytokine and chemokine expression profiles at 48h and 72h post-stimulation. S1 Table. Proteins identification by LC-MS/MS.

## Data availability

MALDI-TOF MS and MS/MS spectra are available on GlycoPost (GPST000569; Preview URL: https://glycopost.glycosmos.org/preview/109642856967ea88d8c3e9a; PIN CODE: 7082).

## Acknowledgement

This work has been funded by the Programme “Top Vet Science” financed by the Vetmeduni Vienna. The authors acknowledge Karin Hummel and Sarah Schlosser at VetCore Proteomics for providing LC-MS/MS services and the BOKU Core Facility Mass Spectrometry for the access to their MALDI TOF MS instrument.

## Author Contributions

Conceptualisation: S.Y., K.L., I.B.H.W., D.W.

Funding acquisition: S.Y., K.L.

Investigation: S.Y., K.L., D.W.

Methodology: I.A., L.N.W., F.SG., Z.D., H.W., S.B.R

Supervision: S.Y., K.L.

Visualisation: I.A., Z.D., S.Y.

Writing – original draft: I.A., L.N.W., S.Y.

Writing – review & editing: I.B.H.W., A.J., K.L., F.SG., T.W., D.W.,

