## Supplementary Text for "Glycoengineering of nematode antigens using insect cells: a promising approach for producing bioactive vaccine antigens of the barber’s pole worm *Haemonchus contortus*"

**S1 Text.** Supplementary methods and results.

### Materials and Methods

*Primers for DNA sequencing and cloning*

A set of DNA oligos were designed and employed in this study, serving as primers for different purposes (i-iv). Both the GoTaq® DNA polymerase (Promega) and Q5® High-Fidelity DNA polymerase (New England Biolabs) were used for the PCR amplification of DNA fragments. The former was generally used for DNA sequencing, TA cloning and PCR screening of *E. coli* clones, whereas the latter was only used for the molecular cloning of target genes for subsequent protein expression.

(i) Primers for amplifying the coding regions (CDS) of *h11* isoforms of the KB strain: Primers targeting all the seven identified CDS of *h11* isoforms were used for PCR amplification using *H. contortus* cDNAs as template. The goal was to deduce the protein sequences of H11s in the KB strain, which can be engineered and recombinantly produced in insect cells.

| Name | Sequence (5' to 3') | Related isoform | Annealing temperature used for Taq polymerase |
|---|---|---|---|
| H11_X94187.1_F | ATGACGTCGCAGGGGAGAAC | KB_H11 | 64°C |
| H11_X94187.1_R | CGAATTACAAGGTGGCTTTCTTGAAG | | |
| H11-1_AJ249941_F | ATGACAGCAGAGGAGAGTCAGG | KB_H11-1 | 62.2°C |
| H11-1_AJ249941_R | TTATGAATTAGATTTTTTGAAGAAAGCTGCTAG | | |
| H11-2_AJ249942.2_F | ATGACGGCGGAGTGGCAG | KB_H11-2 | 62.8°C |
| H11-2_AJ249942.2_R | CTATGATCTTGCTCTCTTGAAGAATTCCG | | |
| H11-4_AJ311316.1_F | ATGACGGCACGAGAGAGGAAAC | KB_H11-4 | 64°C |
| H11-4_AJ311316.1_R | TTACCAGGTAGCGTTCTTAAAGAATGAGG | | |
| HCON_6240_F | ATGACGGTACAATGGACTAAACGG | KB_H11-5a | 63°C |
| HCON_6240_R | TCATGAAGTAGATTTTTCGAAAAAATCCG | | |
| HCON_6250_F_TMD2 | GGCCTTTCAACCGGGCTTACG | KB_H11-5b (truncated) | 55 °C |
| HCON_6250_R | TCATTGAGTAGATTTTTCAAAGAAATCC | | |
| HCON00156260_F | ATGACGGTGGAATGGACTAAACGG | KB_H11-5c | 55°C |
| HCON00156260_R | TCATCGAGTAGATTTTTCGAAGAAATCCG | | |

(ii) Primers for DNA sequencing (synthesised by LGC Genomics)

| Name | Sequence (5' to 3') | Related isoform |
|---|---|---|
| Hc-H11-4.AJ311316_seqF1 | CCGCATATGGTCACGCCC | KB_H11, KB_H11-4 |
| Hc-H11-1.AJ249941_seqF3 | CATCTGGTGCTATGGAGAACTGG | KB_H11-1 |
| Hc-H11-2.AJ249942_seqR3 | GCATCGCTTATGATAGCGTTCC | KB_H11-2 |
| Hc-H11-2.AJ249942_seqR4 | CAATGCCGTTCGATGCAGCC | |
| HCON6240_F1 | TATGGTCACGACCAGAGGC | KB_H11-5a |
| HCON6240_R1 | TAGCCGATGTCGACTCCGGCTC | |
| HCON6240_F2 | GTGTCAGCATCGCCGCCCC | |
| HCON6240_R3 | GATATGATCCAGTCACCGCTGAC | |
| HCON6250_F1 | GGTGTTCGGTTCCGAATATGGTC AC | KB_H11-5b |
| HCON6250_F3 | CCTCTGTGGTATCAAGAAGGCG | |

| | | |
|---|---|---|
| HCON6250_R1 | CGCTGTCGGTTCCATATGAGTTG C | |
| HCON00156260_F1 | GATGATAGATACTACACACCG | KB_H11-5c |
| HCON00156260_F2 | CGACTCGCTATTCTTTGAAG | |

(iii) Primers for molecular cloning using the Gibson assembly approach

| primer name | Sequence (5' to 3') | Annealing temperature |
|---|---|---|
| GAsynt_ HII/1/2_F | GCGCGCGGAATTCAAAGGATGAAATTCTTAGTCAACGTTGC | 56.6-64.9°C |
| GAsynt_ HII/1/2_R | CAAGTGAGCTCGTCGACGTAGGGGGATTCCAAGTATCACGATC | |
| Polyhedrin forward | AAATGATAACCATCTCGC | 57°C |
| SV40pA-R | GAAATTTGTGATGCTATTGC | |

Synthetic DNA was amplified using GAsynt_HII/1/2_F and GAsynt_HII/1/2_R. Eight replicates of PCR reaction mix were prepared as follows: 0.25 µl Q5 High-Fidelity Polymerase, 0.5 µM of each primer, 1x Q5 Reaction Buffer, 1 µl of 10mM (each dNTP) PCR nucleotide mix, 1 µl of 1:10 diluted synthetic DNA, and nuclease-free water to a final volume of 25 µl. The thermocycler was set at 30 s initial denaturation (98°C), followed by 40 cycles of 10 s denaturation, 30 s annealing (linear gradient), and 2 min elongation (72°C), and 10 min of final elongation. The linear gradient was set at 60±1.5°C and each replicate assigned to a specific annealing temperature (56.6, 57.2, 58.8, 60.0, 62.7, 63.7, 64.3, 64.9). Aliquots of the PCR products were visualized on agarose gel, followed by DNA assembly of the selected product into the linearized pACEBac1 construct.

Following transformation into NEB® 5-alpha, positive clones were selected by PCR screening using primers Polyhedrin forward and SV40pA-R. Reaction mixes contained 5 µl of GoTaq® G2 Hot Start Green Master Mix, 0.2 µM of each primer, 4.3 µl of nuclease free water and 0.5 µl of template. The thermocycler was set at 2 min of initial denaturation, followed by 30 cycles of 30 s at 95°C, 30 s at 57°C and 3 min 50 s at 72°C, completed by a final 10 min elongation step. Additionally, Sanger sequencing of purified plasmids was performed at LGC genomics (Berlin), with standard sequencing primers pFASTBacF (ATTAAAATGATAACCATCTCGC) and pee13.4seqrev (TGCATTCATTTTATGTTTCAGGT).

(iv) Other primers

PCR screening primers to verify the large pACEBac1 constructs containing the glyco-module (PCR screening of transformants post CreLoxP recombination and transformation to Pir1)

| Primer name | Sequence (5' to 3') |
|---|---|
| Ce galt-1_full length_F | ATGCCTCGAATCACCGCC |
| Ce galt-1_full length_R | CTACAAGTCTAAAAGACCAACAACTG |
| GAsynt_ HII/1/2_F | GCGCGCGGAATTCAAAGGATGAAATTCTTAGTCAACGTTGC |
| GAsynt_ HII/1/2_R | CAAGTGAGCTCGTCGACGTAGGGGGATTCCAAGTATCACGATC |

| | |
|---|---|
| Cefut-6 full-length F_1 | TCAATGGAGGAAAGAATGGAA |
| HSV_Rev | CAACACCCGTGCGTTTTATT |
| Ce_aman-3Seq FOR_5 | GAGATTTCTATAAGTTGCCAGTTCAA |
| SV40pA-R | GAAATTTGTGATGCTATTGC |
| P1_H11_F | GATAAAGTCATGGAACTATACA |
| P1_H11-1_F | GTCAGCAGGCGACTGACTGT |
| P1_H11-2_F | CATTTTTTACGATGAAGTGA |
| P1_H11-4_F | GTAGTTCAACCTACTGTAATGG |
| P1_H11-5a_F | GAAGGAGGACAAGTAGCGTTCG |
| P1_H11-5b_F | GTTAAAGAAGGTGGCGAAGTAGCG |
| P1_H11-5c_F | CTTACTGTTATGGAGTGAAGGAG |
| P1_GA1_F | AGAGAAAGAGAAGTATCAGAAG |

Three PCR screening reactions were performed to verify the pACEBac1 constructs carrying the glyco-module. In a first step, a multiplex PCR was performed with primers Ce galt-1_full length_F, Ce galt-1_full length_R, GAsynt_ HII/1/2_F, and GAsynt_ HII/1/2_R (A). Primers Cefut-6 full-length F_1, HSV_Rev, Ce_aman-3Seq FOR_5, and SV40pA-R, were used in the second multiplex reaction (B). A third PCR screening was performed using the respective gene specific forward primer and SV40pA-R (C). Reaction mixes contained 5 µl of GoTaq® G2 Hot Start Green Master Mix, 0.2 µM of each primer, 0.5 µl of template and the appropriate amount of nuclease-free water to a final volume of 10 µl. The thermocycler was set at 2 min of initial denaturation, followed by 30 cycles of 30 s at 95°C, 30 s at 58°C (A) or 57°C (B, C) and 3 min 50 s (A) or 1 min 30 s (B, C) at 72°C, completed by a final 5 min elongation step.

PCR screening primers to verify the Bacmids

| **Primer name** | **Sequence (5' to 3')** |
|---|---|
| M13For | GTAAAACGACGGCCAG |
| M13Rev | CAGGAAACAGCTATGAC |
| P1_H11_F | GATAAAGTCATGGAACTATACA |
| P1_H11-1_F | GTCAGCAGGCGACTGACTGT |
| P1_H11-2_F | CATTTTTTACGATGAAGTGA |
| P1_H11-4_F | GTAGTTCAACCTACTGTAATGG |
| P1_H11-5a_F | GAAGGAGGACAAGTAGCGTTCG |
| P1_H11-5b_F | GTTAAAGAAGGTGGCGAAGTAGCG |
| P1_H11-5c_F | CTTACTGTTATGGAGTGAAGGAG |
| P1_GA1_F | AGAGAAAGAGAAGTATCAGAAG |
| GAsynt_ HII/1/2_R | CAAGTGAGCTCGTCGACGTAGGGGGATTCCAAGTATCACGATC |
| Ce galt-1_full length_F | ATGCCTCGAATCACCGCC |
| Ce galt-1_full length_R | CTACAAGTCTAAAAGACCAACAACTG |
| Cefut-6 full-length F | ATGTCTCAAATAGGCGGTGC |
| Cefut-6_M_Rev | CTGAACATCAGGGCAAGTCG |
| Ce_aman-3Seq FOR_5 | GAGATTTCTATAAGTTGCCAGTTCAA |

| | |
|---|---|
| aman-3 Seq REV_1 | ACGACCACCATCGTTCAAA |

Several PCR screenings were performed to verify the bacmids, two for bacmids without the glyco-module, and five for those carrying the glyco-module. In both cases, one reaction using the backbone specific primers M13For and M13Rev was performed. In a second screening step of bacmids without glyco-module, a triplex PCR using both backbone specific primers and the respective gene specific forward primer P1 was carried out. In the case of bacmids containing the glyco-module, the gene specific forward primers P1 were combined with M13For and GAsynt_HII/1/2_R in the second reaction. In the third, fourth, and fifth reaction, M13For was combined with a glyco-gene specific forward and revers primer for either *galt-1*, *fut-6*, or *aman-3*. All reaction mixes contained 5 µl of GoTaq® G2 Hot Start Green Master Mix, 0.2 µM of each primer, 0.5 µl of template, and nuclease-free water to a final volume of 10 µl. The thermocycler was set at 2 min of initial denaturation, followed by 30 cycles of 30 s at 95°C, 30 s at 55°C, and either 5 min (bacmids without a glyco-module) or 3 min (bacmids with glyco-module) at 72°C, completed by a final 5 min elongation step.

### Results

*DNA sequences of the native h11 CDS in KB strain*

Sequencing data were obtained by performing Sanger sequencing on either the selected positive plasmid DNAs post TA cloning, or directly on PCR amplicons in case of the *h11-5a* and *h11-5b*. In total, seven *h11* coding sequences were obtained by assembling overlapping DNA fragments using the SnapGene software. The assembled sequences below were used to deduce the native protein sequences of H11 antigens produced in the KB strain.

>*h11_KB strain*, full-length (2919 bp)
ATGACGTCGCAGGGGAGAACGCGGACATTGCTGAATCTGACTCCAATCCGTCTAATTTT
TGCATTATTTCTAGTAGCTGCTGCAGTCGGCCTCTCTATTGGTCTCACCTATTACTTTAC
TCGCAAAGCGTTCGATACCTCAGAAAAGCCAGGGAAGGATGATACTGGTGGCAAGGGC
AAAGACAATTCTCCCTCTGCGGCGGAACTACTTCTTCCAACCAATATAAAACCATTGTCT
TACGATTTGACGATCAAAACATATCTACCTGGTTATGTGAACTTCCCACCGGAGAAGAAT
CTCACATTCGATGGGCGTGTGGAAATATCAATGGTTGTAATTGAGCCAACAAAGAGTAT
CGTGCTCAATTCAAAGAAGATCTCTGTGATACCCCAAGAATGTGAACTGGTATCGGGCG
ATAAAAAACACGAAATTGAAAGTGTAAAGGAGCACCCAAGACTGGAAAAGGTCGAGTTT
CTTCTTAAGAACCAACTGGAAAAAGATCAACAAATCTTGCTCAAGGTCGGCTATATCGGC
CTCATCAGCAACAGTCTTGGAGGAATCTACCAGACCACTTACACCACCCCGAATGGCAC
CCCTAAGATCGCTGCAGTTTCACAAAATGAGCCCATAGATGCTCGTCGAATGGTACCAT
GCATGGATGAACCGAAATACAAAGCAAACTGGACCGTTACTGTCATTCACCCAAAAGGC
ACCAAAGCCGTCTCGAATGGAATCGAAGTGAACGGAGATGGAGAGATCAGTGGTGATT
GGATCACATCGAAGTTCTTGACTACTCCACGGATGTCATCCTACTTGTTGGCAGTTATG
GTTTCAGAATTTGAATACATTGAAGGTGAAACAAGGACGGGTGTCCGGTTCCGCATATG
GTCACGCCCAGAGGCCAAGAAGATGACAAAACTTGCTTTGGATTATGGTATCAAATGCA
TAGAGTTCTACGAAGATTTCTTTGATATCAAATTCCCTCTGAAAAAACAAGATATGATCGC
CCTTCCTGATTTCTCAGCAGGAGCCATGGAGAACTGGGGTCTTATCACTTACAGGGAAA
ACTCTTTGTTGTACGATGACAGATTCTATGCACCGATGAATAAACAGCGAATTGCTCGCA
TTGTTGCTCATGAGCTTGCCCATCAGTGGTTTGGGGACTTGGCTACAATGAAGTGGTGG
GATAATCTGTGGTTGAATGAAGGTTTTGCAAGATTCACGGAATTCATTGGAGCTGGTCA
GATAACTAAAGATGACGCCAGAATGAGGAACTACTTTCTGATTGATGTACTTGAACGCG
CTTTGAAAGCTGATTCGGTAGCGTCAAGCCATCCACTTTCCTTCAGAATCGACAAAGCT
GCAGAAGTTGAAGAAGCCTTTGATGATATCACATACGCCAAAGGAGCTTCTGTTCTTACT
ATGTTGAGAGCCTTGATTGGAGAAGAAAAACATAAGCATGCAGTATCGCAGTACCTCAA
GAAGTTCTCGTATAGCAATGCAGAAGCGACAGATCTATGGGCAGTTTTCGATGAAGTTG
TCACTGACGTCGAGGGTCCAGACGGCAAACCTATGAGAACCACGGAATTCGCAAGTCA
GTGGACGACTCAGATGGGCTTCCCAGTTATTTCCGTAGCAGAGTTTAACTCGACTACTT
TGAAATTAACGCAAAGTCGATATAAGGCGAATAAAGATGCTGTGGAGAAAGAGAAGTAC
CGTCATCCGAAATACGGATTTAAATGGGATATTCCACTGTGGTATCAGGAAGGCGATAA
GAAGGAGATAAAGCGAACATGGTTGAGAAGAGATGAACCGCTTTACTTGCATGTAAATG
ATCCTGGCGCTCCCTTTGTGGTGAATGCAGACCGCTATGGATTTTATCGACAAAATCAT
GACGCTAGTGGTTGGAAAAAGATAATCAAGCAGCTCAAGGATAATCATGAGGTTTACAG
TCCCCGGACAAGAAATGCCATCATTAGCGATGCGTTTGCTGCGGCTACAACTGACGCAA
TTGAGTATGAGACTGTATTTGAACTTCTGAAATATGCCGAAAAAGAAACGGAATATCTAC
CATTAGAAATCGCAATGTCCGGGATCTCTTCGATTTTGAAATACTTCGGTACCGAGCCA
GAGGCAAAGCCAGCTCAAACATACATGATGAACATATTGAAACCGATGTATGAAAAAG
CAGTATCGACTTCATTGCCAATAACTACAGAAATGACAAGCTGTTTTTCCAAATCAACCT
CCAAAAAGATGTCATTGATATGTTCTGCGCCCTCGGATCGCAAGACTGCAGGAAGAAAT
ATAAAAAACTTTTCGATGACGAAGTCATGAACAAATGCAGGGATGGTCAAGCAGCAACC
GAATGCGAAAGAATCGCCGCTCCTCTCCGATCAAGTGTTTATTGTTATGGTGTGAAGGA

AGGCGGTGATTATGCTTTCGACAAGGTGATGGAGCTTTATACGGCCGAAACACTCGCCC
TAGAAAAAGACTTCCTACGCCTAGCATTGGGATGTCATAAAGATGTTACTGCTTTGAAAG
GACTTCTCTTGCGGGCTCTGGACAGGAATTCGTCGTTCGTACGTATGCAGGATATCCCA
AGTGCTTTCAATGATGTAGCAGCAAATCCTATCGGCGGAGAATTCATTTTCAATTTCCTT
ATTGAAAGATGGCCAGATATCATTGAAAGTATAGGAACGAAGCACACATACGTTGAGAA
AGTGATACCAGCCTGCACTTCAGGAATCCGCTCACAACAGCAGATTGACCAGCTGAAGA
ATCTGCAGAAAAATGGCATGAACGCTCGTCAATTCGGTGCATTCGATAAAGCAATCGAA
CGAGCACAAAATAGGGTGGATTGGATTAAAAAACATTTCCAAAAATTAGCGGCTTTCTTC
AAGAAAGCCACCTTGTAA

>*h11-1_KB strain*, full-length (2934 bp)
ATGACAGCAGAGGAGAGTCAGGAGCAGGAGACGCAGCAACCACGAAAAAATACAGTGC
TACGGCTCACCCCAATCAAGTCTCTCTTTGCTTTGTTAGTGGTAGCTGCTGCCGTCGGC
CTCTCAATCGGTCTCACCTATTACTTTACAAGGAAAGCTTTTGATACTACTGGCGGAAAT
GGAAAAGAAGATCAACCTATTGTCGATGATAATTCCCCATCAGCTGAAGAATTACGTCTC
CCAACAACCATAAAACCTTTGACTTACGACTTAGTAATCAAAACGTATCTGCCAAACTAT
GTAAACTATCCACCTGAGAAAGATTTCGCTATTGATGGGACTGTGGTGATTGCTATGGA
AGTTGTGGAGCCAACAAAGTCCATAGTGCTCAACTCGAAAAATATTTCTGTAATTGCAGA
CCAGTGCGAACTGTTTTCTAACAACCAAAAACTCGACATCGAAAAGATTGTGGATCAGC
CAAGGCTGGAGAAAGTCGAATTCGTTTTGAAGAAAAAGCTGGAGAAAAATCAGAAAATC
ACGCTCAAGATTGTATACATTGGCCTTATCAACGACATGCTGGGAGGACTTTATCGAAC
AACCTATACAGATAAAGATGGTACAACTAAGATTGCTGCATGCACTCATATGGAACCGAC
AGACGCCCGTCTTATGGTCCCCTGTTTCGACGAGCCGACGTTTAAGGCAAACTGGACC
GTGACAGTAATTCATCCGAAGGGCACCAGTGCCGTGTCAAATGGAATAGAAAAGGGAG
AAGGAGAAGTCTCTGGCGATTGGGTCACAACCAGATTCGATCCAACCCCGCGAATGTCT
TCGTATTTGATTGCTCTTGTGATTTCCGAATTTAAGTACATTGAAAATTATACGAAAAGCG
GTGTTCGGTTCCGAATATGGGCTCGTCCGGAAGCTATGAAGATGACAGAATATGCCATG
ATAGCTGGAATCAAATGTTTGGATTACTATGAGGACTTCTTCGGGATCAAATTCCCACTT
CCAAAACAAGATATGGTTGCTCTTCCTGACTTCTCATCTGGTGCTATGGAGAACTGGGG
TCTCATCACATACAGGGAGGGTTCCGTGCTCTACGATGAAAACCTCTACGGACCAATGA
ATAAGGAGCGGGTTGCAGAAGTGATCGCGCACGAACTTGCACATCAGTGGTTCGGTAA
TTTGGTCACGATGAAGTGGTGGGATAACCTATGGCTGAACGAAGGATTCGCGTCATTCG
TGGAATACATCGGAGCCGACTTCATCAGCGATGGTCTATGGGAAATGAAAGATTTCTTC
CTGCTGGCACCGTACACAAGTGGTATTACGGCTGATGCAGTAGCTTCAAGCCATCCGCT
TTCCTTCAGAATAGATAAGGCTGCAGATGTATCAGAAGCGTTCGATGATATCACATACCG
TAAAGGAGCATCCGTTCTTCAAATGCTATTGAATTTAGTTGGGGACGAAAATTTCAAGCA
GTCTGTTTCGCGTTACCTCAAGAAGTTTTCATATGATAATGCGGCTGCTGAAGATTTATG
GGCAGCATTCGACGAAACCGTCCAAGGTATAACCGGACCTAATGGTGGACCATTGAAAA
TGTCCGAGTTTGCGCCACAATGGACAACTCAGATGGGGTTCCCTGTTCTTACTGTCGAG
TCGGTTAACGCAACGACTTTGAAAGTCACCCAAAAACGATACAGGCAGAACAAGGATGC
AAAGGAACCAGAGAAGTACCGTCATCCAACTTATGGGTTCAAATGGGATGTTCCTCTGT
GGTATCAGGAAGATGAACAGCAAGTGAAAAGAACTTGGTTAAAAAGAGAGGAACCGCTC
TATTTCCATGTAAGCAATTCTGATTCGTCAGTTGTGGTGAATGCCGAACGTCGTGCTTTT
TGCCGATCAAACTATGACGCTAACGGTTGGAGGAACATTATGAGAAGACTCAAGCAGAA
TCATAAGGTCTATGGTCCACGAACAAGAAACGCTCTCATAAGTGATGCGTTTGCAGCAG
CTGCAGTTGAGGAAATGAATTACGAGACCGTATTTGAAATGCTCAAATACACCGTGAAA
GAAGAGGATTACTTACCATGGAAGGAGGCAGTATCAGGATTCAATACAATTTTGGACTTT
TTCGGCAGCGAACCCGAATCTCAATGGGCTTCGGAATACATGCGAAAACTGATGAAGCC
AATTTATGACAAGAGTAGCATCAAGTTTATAGCGGAGAACTACAAAAAAGATTCGCTTTT
CTTCAAAAAATAATCTCCAAATAGCTGTTATTGACACATACTGTGGTCTTGGAGGCAAAGA
ATGTCTTGAAGAAATGAAAAAGCTTTTTGACAAGGAGGTCATGAAATGTCAACCTGGTCA

GCAAGCGACCGACTGCGTAAAGGTAACTGCTCCTCTCCGAAAAACTGTTTACTGCTATG
GGGTCCAGGAAGGCGGTGATGAGGCATTCGACAAGGTGATGGAACTATATAATGCGGA
ACAAGTGCAGTTGGAGAAAGACAGTCTACGTGAAGCATTGGGATGCCATAAAGACGTTA
CAGCTCTAAAGGGACTTCTTATGCTGGCTTTGGATCGGAATTCGTCATTTGTGCGTCTTC
AAGGTGCTCATGATGTGTTTAACATTGTGTCCAGAAATCCTGTTGGAAACGAACTGCTGT
TCAATTTCCTCACAGAGCGATGGGAAGAGATACTTGAAAGTTTGTCAATACGACACAGAT
CAGTTGATCGAGTGATCAGAGCCTGTACTCGAGGACTACGATCCAGGGAACAAGTACAA
CAGTTGAAGAATCTATACAAAAATGACAAACGTGCTCGCGAATACGGTGCATTTGGTGG
GGCAATAGAAAGATCGGAACACAGAGTCAAATGGATTGAGAAACATTTCCGGAAACTAG
CAGCTTTCTTCAAAAAATCTAATTCATAA

>*h11-2_KB strain*, full-length (2919 bp)
ATGACGGCGGAGTGGCAGAAGCGTAGAATCTTGGGCTTCTCACCGATCAGCCTGCTTT
GTACATTATTTGTATTAGCTGCTGCCGTTGGACTCTCCATTGGTCTGACCTATTACTTCA
CTCGTAAAGCATTCGATACCACACAAAAGGAACAGAAGGATGACACCGGTGGTAAAGAA
AAGGATAATTCTCCTTCTGCAGAAGAACTACTTCTTCCAACTAAAATAAAACCAGTGTCG
TACGACTTGAGCATTAAAACACATCTACCGGGTTACGTGAACATTCCACCAGAGAAGAA
TCTTACATTTGATGGCTACGTGGAGATTTCTATGGTTGTCGTTGAGCCAACAAATAGCAT
TGTGCTAAATTCAAAGAAAATCACTTTGGCACAAGGAGGATGCGAATTGTTTTCTGGTAA
TCAAAAACTTGACATCGAAAGTGTAAGGATGCGGGAAAGACTTGACAAGCTTGAGATTA
CCCTCAAAAATCAACTGCAGAAGGAACAGAAAATCCTGCTCAAGATCACTTACACCGGC
CTTATTAGCAACACTCTCGGTGGGCTCTACCAGTCTATCTATACCGATACGGACGGAAC
CACCAAGATCGTTGCTGTTTCACAAAATGAATCATCAGACGCTCGTCGTATAGCGCCAT
GCTTTGACGAACCGAAGTATAAGGCGAAATGGACTGTCACCGTTGTTCATCCCAAAGGT
ACAAAGGCTGCATCGAACGGCATTGAAGCGAATGGAAATGGGGAGCTCCAGGGTGATT
GGATAACATCCAAATTTAAAACTACCCCACCGATGTCGTCCTATTTATTAGCTATTATTGT
TTGTGAATTTGAATACATTGAAGGCAAAACCGAAACAGGTGTACGGTTCCGTATATGGTC
TCGACCAGAGGCGAAAGCAATGACGGCATACGCTTTGGATGCTGGCATCAGATGCCTG
GAGTTCTATGAGAAGTTCTTTGACATAAAATTCCCTCTGGAAAAACAAGATATGATTGCT
CTTCCTGATTTCACCGCTGGTGCCATGGAAAACTGGGGCCTTATCACTTATAGAGAGGA
TTCTCTCCTATACGATGAAAAAATTTATGCACCGATGAATAAACAGCGGGTTGCTCTCGT
AGTTGCTCACGAGCTTGCTCATCTGTGGTTCGGCAATCTGGTCACACTGAAGTGGTGGG
ATGATACGTGGTTGAACGAAGGTTTTGCAACATTTGTTGAGTATCTTGGAATGGACGAAA
TTAGCCACAACAATTTCAGAACGCAAGATTTCTTCTTGCTCGATGGAATGGATCGCGGA
ATGAGAGCTGACTCGGCAGCATCGAGCCATCCGCTTTCGTTTAGGGTTGACAAAGCGG
CAGAAGTTGCCGAAGCCTTTGACGATATTTCATACGCCAAGGGAGCGTCAGTTCTCACT
ATGCTACGGGCTTTGATTGGAGAGGACAATTACAGGAATGCTGTTGTGCAATACCTCAA
GAAGTTCTCCTACAATAATGCACAAGCAGCCGATCTGTGGAACGTCTTCAATGAAGTTG
TCAAAGGTGTTAAGGGTCCTGACGGCAACGTCATGAAAATCGACCAATTTACCGATCAG
TGGACGTATCAGATGGGTTATCCTGTGGTTAAAGTAGAAGAATTTAATGCGACCGCCCT
AAAGGTTACGCAGAGCCGGTACAAGACAAATAAAGACGCCTTGGAACCAGAGAAATATC
GTAATCCGAAATACGGGTTCAAGTGGGATGTTCCCCTATGGTATCAGGAAGGCAATAGC
AAAGAGGTGAAGCGAACATGGCTAAAAAGAGATGAACTGCTGTACTTAAACGTCAACAA
TCGGGATACATCCCTTGTGGTGAATGCTGATCGACATGGATTTTATCGACAAAACTATGA
TGCCAACGGTTGGAAAAAGATAATCAAGCAGCTCAAAGAAAATCACGAGGTCTTCGGTC
CAAGGACAAGGAACGCTATCATAAGCGATGCATTTGCTGCAGCTACGATTGACGCAATC
GACTATGAAACTGTATTCGAACTACTTGAATATGCCAAAAATGAAGAGGAATTCTTGCCT
TGGAAGGAAGCTCTGTCCGGCATGTTCGCAGTTTTAAAGTTCTTCGGTAATGAGCCGGA
GACAAAACCAGCTAGAGCTTACATGATGAGCATATTAGAACCGATGTATAATAAGAGCA
GCATTGATTACATCGTCAAGAATTATTTGGATGATACGTTATTCACAAAAATTAATACTCA
AAAGGATATCATTGATGCATATTGTTCCCTTGGATCAAAGGACTGTATAAAGCAATATAA

GGATATCTTCTACGATGAGGTTATGCCCAAGTGTAAGGCCGGGGAAGCAGCAACCAAAT
GCGTTAAGGTTTCCGCTCCTCTTCGAGCCAATGTTTACTGTTATGGTGTACAGGAAGGT
GGTGAAGAAGCTTTTGAAAAGGTGATGGGGCTGTATCTAGCAGAAGATATTCAACTGGA
GAAGGGTATCCTGTTCAAAGCCTTGGCATGCCACAAAGATGTTACAGCTCTAAAAGAAC
TTCTTTTGCGAGCCCTGGACCGTAAATCGTCGTTTGTGCGTCTTCAGGATGTCCCTACC
GCTTTCCGTGCTGTATCTGAAAACCCTGTGGGCGAAGAATTCATGTTCAATTTCCTAATG
GAGAGATGGGAGGAAATCACTGCGAGCTTGGAAACAGAACACAGAGCAGTTGATAAAG
TGGTCGGCGCTTGTTGCACAGGAATTCGCTCCCAACAACAAATAGATCAGCTGAAGAAT
CTACAGAAGAACAATGCGCAGGCTAAGAAGTTCGGCTCATTCACCCAGGAAATCGAAAA
AGGAGAACATAAAATTGCCTGGATCAAGAAACATTTTCACAGATTATCGGAATTCTTCAA
GAGAGCAAGGTCATAG

>*h11-4_KB strain*, full-length (2916 bp)
ATGACGGCACGAGAGAGGAAACGGTCACTGGTGGAATTCACACCATTCAGCCTGGCTT
TCGCAATGTTTGCAGTGGCTGCTGCAGTAGGTCTGGCCATTGGCCTCACTTATTTCTTTA
CTCGGAAAGCATTTGATCCCACCCAAAAAGACAAAAATCAACCTGGTGGTAAAGAAAAA
GACAATTCCCCGTCTGCAGCGGAACTACTTCTTCCGTCGAACATAAAGCCATTGTTGTA
CGACTTGACCATCAAAACATATCTACCTGGTTATGTGAACTTCCCACCAGAGAAAAATCT
CACATTTGATGGGCGTGTGGAAATTTCTATGGTTGTCGTTGAACCGACAAAAAGTATCGT
GCTCAACGCAAAGAAAATCACTGTAATACCAGCGGAATGCGAAGTGTTATCGGGCACCC
AAAAACTTGATATTGAAAGTGTGAAGGAGCATGAACGGCTCGAGAAGCTGGGGTTCCG
CCTCAAATCACGCTTGGAAAAAGATCAGAAAATTCTGTTAAAGATCACCTACGCCGGCC
TCATTAGTAACACCCTTGGAGGTATTTATCAGACAACTTACACTGATGCGAATGGAAACC
CAAAGATTGCTGCCGTATCTCAAAATGAACCGATAGATGGTCGTCGAATGGTACCATGC
ATGGACGAACCAAAGTACAAAGCGAATTGGACCGTTACTGTGATTCATCCAAAAGGCAC
CAAGGCTGCATCAAATAGCATCGAAATAAATGGAGAAGGAGATGTCAGCGGTGACTGG
ATTACGTCCAAATTCGAAACCACTCCACGAATGTCTTCCTATTTGCTGGCCGTTTTCATT
TCAGAATTCGATTTTGTCGAAGGTCGTACGAAACAAGACGTTCGATTCCGCATATGGTC
ACGCCCAGAGGCAAAGGGAATGACAAAATATGCTTTGGAATCTGGCATCAAATGCATAG
AGTTCTACGAAGATTTCTTTGATATCAAATTCCCACTGAAGAAACAGGATATGATTGCCC
TTCCCGATTTCTCAGCAGGAGCCATGGAGAACTGGGGTCTCATAACCTACAGAGAAAAC
TCCTTGTTGTACGATGAAAAATTCTATGGGCCAACAAATAAACGACGAGTTGCTGTTGTG
GTTGCTCATGAGCTTGCCCACCAGTGGTTTGGCGATTTGGTCACTATGAAATGGTGGGA
CGATCTATGGCTGAACGAAGGCTTTGCAACATTTGTCGAATACATTGGAGCGGATGAAA
TTGGAGACCATTACTTCAACATGCCGGACTTCTTCTTAATCGGTGCCCTTGAGCGTGCT
CTGAAAGCTGACTCGGCAGCTTCCAGTCACCCGCTTTCCTTCAGAATTGACAAAGCCGT
AGAAGTTGAAGAAGCCTTCGATGATATCTCGTACGCCAAAGGAGCGTCGATAATTACAA
TGCTACGAGCTCTGATTGGAGAAGACAAGCACAAACACGCAGTGACTCAATATCTCAAG
AAGTTCTCGTACAGTAATGCACAAGCGTCGGATCTATGGGAAGTATTCGATGAAGTAGT
CACGGATATCAAAGGCCCCGATGGCAAACCCATGAAAACTACAGCATTCGCTGACCAGT
GGACAACTCAGATGGGATTCCCATTGGTAACGGTGGAAGCGTTCAATGCCACTTCTGTA
AAAATATCGCAAAGTCGATTCAAGACCAATAAAGACGCTAAGGAACCAGAGAAATATCG
TCATCCAAAATATGGATTCAAATGGGATATCCCGCTATGGTATCAGGAAGGTGACAACA
AGGAAGTGAAGCAAACCTGGATAAGGAGAGAAGAACCGCTTTATCTGCACGTCAACGA
CCTTAGCAAACCCTTCGTGGTTAACGCCGATCGGCATGGATTTTATCGACAAAATTATGA
TGCTGACGGTTGGAGAAAGATAATCAAGCAGCTCAGGGACAATCATAAGGTTTTCAGCC
CACGGACAAGGAATGCTATCATAAGTGATGCATTTGCATTGGCTTCTGTCAATGCAATC
GAGTACGAAACTGTATTTGAACTACTTAAATACGCTGTTAACGAAGAGGAGTTCATACCT
TGGACAGAAGCAATTTCTGGAATATTCGCCGTCTTAAAATTCTTCGGTAACGAGCCGGA
GTCGAAGCCAGCCGAAGCCTATATGATGAAAATTTTGGAACCGATGTACAAGAAGAGCG
ATTTGGGTTACATTGCCGCTAAGTACAAGGTTGACCAATTATTTTCCAAAATCAATCTGC

AGAAAGATATCATTGATGCGTACTGTGCTCTTGGATCAAAGGACTGTATGAAGAAGTACA
AAGATATCTTTGATCGTGAAGTTATGAACAAATGTAATGACGGTGATGAGGCAACCAAAT
GCGTAAGTGTCGCTGCTCCTCTTCGATCGAGTACTTATTGCAATGGTGTGAAGGCAGGT
GGAACTTACGCATTCGAAAAGGTAAAGGCACTATATTATAAAGAAACTGTTCAACTGGAA
AAGGATATGCTCCTCCGAGCATTAGGCTGCCACAGGGATGTTACAGCTCTGAAAGGCCT
TCTTCTACTTGCGGTGGATCGCAATTCGTCATTTGTCCGTCTTCAGGATATCCCGAACG
CCTTCCAAGCTGTAGCCGCAAATCCTGTTGGCGAAGAATTCATGTTCAATTTCCTCATAG
AAAGATGGGGAGATATCATTGGAAGCATAGGATCAGAACCGACATATGTTGAAAGAGTG
ATACCACCATGTACCTCAGGCATTCGTTCCAAACAGCAGATAGATCAGCTGAGAAATCTT
CACAAAAAATGGCATTCATGCTCAAGAATACAGCACGTTTGTCAAAGAAATTGAACGAGCA
GAGCATAAAGTGGATTGGATTAAAAAACACTTCAAAAAATTAGCCTCATTCTTTAAGAAC
GCTACCTGGTAA

>*h11-5a_KB strain*, a truncated sequence missing the start and stop codons (2895 bp)
CAATGGACTAAACGGACAGTGTTAAAATTCACGCCTATTACCTTACTGCTTTTGTTATTTC
TAGTAGCTGCTTCAATTGGTCTTTCAATTGGGCTCACATACTACTTTACCAGGAAAGCTT
ATGATACTACTGAAAAAAACAAGGATCATGGTGCTGATGATAACTCACCTTCTGCAGAAG
AGCTACGTCTTCCAAAGAATATAGAACCGTTATTGTACGACCTGAGCATCAAAACTTATC
TGCCGGGTTACGTGAGCTTCCCACCTGAGAAGAATCTCACATTTGATGGACAAGTAGGA
ATCTCCTTAAGGGTAGTCGAACCAACCAAAAGTATTGTACTCAACTCAAAGAATATCACC
GTGATACCAGATAAATGCGAACTGTTTTCGGGCGACAAAAAACTTGAAATAGAAAGTATC
AAAGAACATGAAAGACTAGAAAAGTTGGAAATTCTACTGAAAAATCGGCTAGAGAAGGA
CCAAGAGGTCCTGCTGAAGATAATTTATACGGGTATCATCAGCAACACCCTTGGCGGAC
TCTATCAAGCCACCTATACAGATACTGATGGAACCGTCAAGATTGCCGCAGCCTCTCAA
AACGAGCCAACGGACGCTCGTCGTATGGTTCCGTGCTTGGATGAACCAAGTTTCAAAGC
CAATTGGACTGTGACTGTCATCCATCCGAAAGGAACAAAAGCTGTATCGAATGGCATTG
AAACAAATGGGGAAGGGGAAGTCAGCGGTGACTGGATCATATCAAAATTCGAGACAACA
CCTCGGATGTCTTCCTACTTGTTGGCTGTGGTTGTTTCAGAATTTGATTACATCGAAGGG
TTTACAAAAAGTGGTGTTCGGTTCCGGATATGGTCACGACCAGAGGCAATGAATATGAC
AGGATATGCAAAGGACGCCGGCATAAGATGTTTGGAATACTACGAAAACTTTTTCGATAT
CAAATTTCCTCTCAAAAAGCAAGATATGGTTGCTCTTCCTGACTTCTCCGCTGGTGCCAT
GGAAAATTGGGGTCTCATTACTTACAGAGAGAACCAGTTATTGTATGATGATAGATACTA
TGGACCGATCAACAAGCAGCGGGTTGCCCTTGTCGTTGCTCATGAACTTGCCCACCAGT
GGTTTGGCAATCTGGTCACACTGAAATGGTGGGACGACACGTGGCTAAACGAAGGCTT
TGCAACATTTGTGGAATACATTGGAACAGATGAAATTAGCTATAAAAACTTCAGGATGGA
CGACTTTTTCCTACCGGATGCACTTGCAATCGGCTTGGATGCTGATGCGGTATCTTCAA
CTCACCCACTCTCGTTTAGAGTTGATAAAGCTGCAGAAGTTGCTGAAGCTTTTGACGAAA
TTACCTACGCCAAGGGAGCATCTGTTTTGGTAATGCTACAGGCTGTAATTGGCGAGAAA
AACTACAAGCAAGCTGTTACGCAATACCTGAAAAAGTTTTCGTACAGCAACGCCCAAGC
TACTGACCTTTGGAACGTTTTTGACGAAGTTGTCAAGGACGTTAAAGGTCCAGACGGAA
ATCTCATGAAAACGACCGAATTCGCATCTCAATGGACTACTCAGTTGGGATTTCCTTTGG
TCACTGTGAAAGCTTTCAACGCAACTTCCTTGCAAATAACACAAACTCGGTATAAGACGA
ACAAAGACGCCCTGGAGCCGGAGAAGTATCGTCATCCAAAATACGGATTCAAGTGGGA
CATTCCTCTGTGGTATCAGGAAGGCGATAACAAAGATATAAAGTTAGCATGGTTGACAA
GAGAAGAACCGCTCTATTTGCACGTTAGCAATCCTGACACTTCCATTGTCGTGAATGCG
GATCGCCATGGATTTTATCGACAGAACTATGACGCTAACGGTTGGCGAAAGATCATCAA
GCAGCTTAAGGAAAATCATAAGGCATACAGTACTAGGACGAGAAATGCGATTATAGGCG
ATGCATTTGCAGCGGCTCTGATTGACCAACTTGAGTACGAAACTGTATTTGAACTGCTTG
AGTACGCTAAAAATGAAGAGGAATATTTGCCATGGACAGAGACAATATCTGGTTTTTATG
CCATCTTAGACTTCTTCGGAAATGAGCCGGAGTCGACATCGGCTAAAACTTTCATGATG
AATATATTGAAACCGATGTATGAGAAAACCAGTATGAAATATATCGCCGACAACTACAAA

AACGATTCGCTATTCTTTGAAATGAACCTCCAAAAATCTGTTATCAATGCCTATTGCTTCC
TCGGATCCATGGAATGTATCAAAAATTACACAGATTTGTTTGATAAAGAAGTTATGAAGA
AATGTAAAGACGATGACGAAGCAAGCAAATGTGTCAGCATCGCCGCCCCTCTTCGAGCA
AAAGCCTACTGCTATGGCGTGAAAGAAGGCGGACAAGTTGCTTTCGATAAGGTGATGAA
ATTATACTATGCAGAAAACGTTCAACTGGAAAAAGACGTGCTACTTCAAGGATTAGGATG
TCACAAAGATATTACAGCATTGAAGAGGCTCCTCCTCCTAGCTCTGGATCGAAATTCCTC
ATTTGTTCGACTTCAAGATGTGACTGATGTCTACGATGCTGTATCATCAAATCCTGTTGG
CGAAGAATTCATGTTCAACTTCCTCCTTGAAAGATGGGAAGAAATTCTTGAAAGCTTGAC
AACAGAGCACCGAGCTGTTGAACGAGTGATCGAAGCGTGCACTGCAGGTATTCGATCG
GAGCAGCAAATCGATCAGCTGAGGAGCCTTCAAAAAAATGGTGCACATGCTCGAGAATA
TGGAGCATTCGACGAACAAATCGAACGAGCGGAACATAAAATCAACTGGATCAAGAAAC
ATTTGCGAAAATTATCGGATTTTTTCGAAAAATCTACCTTC

>*h11-5b_KB strain,* a truncated sequence missing the start and stop codons (2756 bp)
ACGTACTTTACTCGGAGAGCTTATGATACTACTGAGAGGAATAAAGATTATTGTGCTGAT
TACAATTCACCTTCTGCACAAGCACTGCGTCTTCCAAGGAACATAGAACCGTTATTGTAT
GATTTGTGCATCAAAACTTATCTGCCAAATTACGTGGACTTTCCACCCGAGAAGAATCTC
ACATTTGATGGACAAGTAGGAATCTCTTTGAGGGTGCTTGAACCAACCAAAAGTATTGTA
CTCAACGCAAAGAATATCACGGTGATACCGAATAAATGCGAACTGTTCCTGGGCAACCA
AAAACTCAAAATAGAAAGTGTCAAAGAACATGAAAGACTAGAAAAGGTGGAGTTTCTACT
GAGAAATTGGCTACAGAAGGATCAAAAAGTCCTGTTAAAGATAGTTTATACGGGTCTCAT
TAGCAACACACTTTTCGGACTTTATCAAGCCACCTATACAGACCCGGATGGAACCGTCA
AGATTGCTGCAGCAACTCATATGGAACCGACAGCGGCTCGTCGTATGGTACCATGCTTA
GATGAGCCTAGTTTCAAAGCCAATTGGACTGTGACTGTCATACATCCAAAAGGGACAAA
AGCTGTGTCGAACGGCATCGAAACAAATGGAAAAGGAGAAGTCAGTGGTGACTGGATC
ATATCGAAATTTGAGACAACTCCACGGATGTCTTCCTATTTGTTGGCTGTAGTTGTTTCA
GAATTTGATTACATTGAAGGATTTACAAAAACTGGTGTTCGGTTCCGAATATGGTCACGA
CCGGAGGCAAAAAATATGACAGAATATGCAAGGGATGCCGGCATAAGATGTTTGGAATT
CTACGAAAATTTCTTCGAATCAAAATTTCCTCTCAAAAAACAAGATATGGTCGCTCTTCCT
GATTTCTCTTTTGGTGCTATGGAAAATTGGGGTCTCATCACTTACAGAGAGACTAGTCTG
TTGTATGATGATAGATACTATGGACCGATCAACAAGCAGCGGGTTGCCCTTGTCGTTGC
CCATGAACTTGCTCACCAGTGGTTTGGCAATTTGGTCACGCTGAAATGGTGGGATGATT
TGTGGCTGAATGAAGGCTTTGCAAGACGTGTTGAATACATTGGAACAGATGAAATCAAC
AATAAAACCATCAGGATGGACGACGTTTTCCTACCGAATGCACTTGTAAAAGCTTTGGAC
GCTGATGCGGTATCTTCAACCCATCCGCTCTCGTTCAGAATTGATAAAGCTGCAGAAGT
TCTTGAAGCCTTTGATACAATTACATATGAGAAAGGAGCGTCCGTTTTGGAAATGCTACG
GGCTGTAATTGGCGATAAAAACTTTAAGCAAGCAGTTACGAAATACCTGAAGAAGTTCTC
ATACAGAAACGCCAAGCCTTCTGATCTGTGGAACGTTTTCGACGAAGTTGTCAAGGATG
TCAGAGGCCCGGACGGCAACAGCATGAAAACCACCAAATTCGCACCTCAGTGGACTAC
GCAGTTGGGATTCCCTCTGGTTGCTGTGAAAGATTTCAACGGAACTTCTGTGAAAATAA
CTCAAACTCGTTACAAGGCCAATAAAGACGCCCTCGAGCCAGAGAAGTATCGCCATCCG
AAATACAGATTCAAGTGGGATATTCCTCTGTGGTATCAAGAAGGCGATAACAAAGATATA
AAGTTTGCATGGTTGACTAGAGAAAAACCGCTCTATTTGCACAAGACCAAGCCTGACAC
TTCCATTGTGGTGAATGCGGATCGCCATGGATTTTATCAACAGAACTATGATGCTAAGG
GATGGAGAAAGATCATCAAACAGCTCAAGAAAAATCATAAGGCATACAGTGCAAGGACG
AGAAATGCGATCATAGGCGATGCATTTGCAGCGGCTCTGATTGATGAACTTGAATACGA
AACAGTATTTAAACTGCTTGAGTACGCCAAGAATGAAGAGGAGTATTTGCCATGGACAG
AGACATTATCTGGTTTTGATGCCATTTTAAACTTCTTCGGAAATGAGCCGGAGTCGACAT
CGGCTAAAGCTTTCATGAAGAATATATTGAAACCGATGTATAAGAAAACCAGTATGAAAT
ATATTGCCGTCAACTACAAAAACGACTCGCTATTCTTTGAAGTGAACCTGCAAACATCTA
TTATCGATGCTTATTGCCACCTTGGAGCTAGGGAATGTATCAAAAACTACACAGATTTGT

TTGATAAAGAAGTTATGAAGAAATGTAGAGACGGTGACAAAGCAAGCAAATGTGTAAGC
ATCGCCGCCCCTCTTCGAGCGAAAGCCTACTGCTATGGCGTGAAAGAAGGCGGAGAAG
TTGCTTTCGATAAGGTAATGAAATTATGCTATGCAGAAAATGTTCAAGTGGAGAAAGACG
TGCTACTCAAAGGATTAGGATGTCACAGAGATATTACAGCATTGAAGAGGCTACTCCTC
CTGGCACTGGATCGAAATTCCACATTTGTCCGTCTTCAAGATGTGGCTGCTGTTTACGAT
GCTGTATCAGCGAATCCCATTGGCAAAGAATTCATGTTCAACTTCCTCCTAGAGAGATG
GGAAGAAATTCTCGAAAGCTTGACAGCGGACCACCGAACTGTTGAACGAGTGATCAAAG
CGTGCACTGCAGGTATTCGATTGGAGCAGCAGATAGATCAGCTGAGAAGCCTTCAGAAA
AATGGTGAACATGCTCGAGAATATGGAGCATTCGACGGACAAATCGAACGTGCGCAACA
TAAAATCAACTGGAACAAGAAACA

>*h11-5c_KB strain, a* full-length contig (2841 bp)
ATGACGGTGGAATGGACTAAACGGACAGTGTTAAAATCTGCGTCCATCAGCATACTGCT
TATATTATTTTCTGTAGCTGCTTCGATTGACCTCCCTGCTGATGATAGTTCACCTTCTGCA
CAAGAACTGCGTCTTCCAAGGAACGTAGAACCGTTACTGTACAATCTCAGCATCAAAAC
TTATTTGCCGAGTTACGTGAACTTCCCACCAGAGAAGAATCTCACATTTGATGGACAAGT
AGGAATCTCCTTGAGGGTGGTTGAACCAACCAAAAGTATTGTACTCAACGCAAAGAATA
TCACGGTGATACCGAATAATTGCGAACTGTTTTCGGGCGGCAGAAAACTTGAAATAGAA
AGTGTCGAAGTACATGAAAGACTAGAAAAGGTGGAGTTTCTACTGAGAAATCGGCTACA
GAAGGATCAAAAAGTCCTGTTGAAGATAATTTACACGGGTGTCATTAGCAACAGCCTTTT
CGGTCTTTATCAAGCCACCTATACAGATGCAGATGGAAACGTCAAGATTGCTGCAGCAA
CTCAGCTTTGCCCAAGTGATGCTCGTCGTTTGGTACCATGCTTAGATGAACCAAGTTTCA
AAGCCAGTTGGACTGTGACTGTCATACATCCAAAAGGGACAAAAGCTGTGTCGAACGGC
ATCGAAACAAATGGAAAAGGAGAAGTCAGTGGTGACTGGATCATATCGAAATTTGAGAC
AACACCACGGATGTCTTCCTATTTGTTGGCTATAGTTGCTTCAGAATTTGATTACGTCGA
AGGATTTACAAAAAGTGGTGTTCGTTTCCGAATATGGTCACGACCAGAGGCGAAAAATA
TGACAGCATATGCAAGGGATGCCGGTATAAGATGTTTGGAGTTCTACGAAAACTTCTTC
GATATCAAATTCCCTCTTAAAAAACAAGATATGGTTGTACTTCCTGATTTCTCTTTTGGCG
CCATGGAAAATTGGGGTCTCATCACTTACAGAGAGAACCGGCTGTTGTATGATGATAGA
TACTACACACCGATCAACAAGCAACTAGTTGCACTTGTCGTAGCTCATGAGCTTGCCCA
TCAGTGGTTTGGCGATTTGGTCACGCTGAAATGGTGGGACGATTTGTGGCTAAATGAAG
GCTTTGCAAGATTTGTTGAATACATTGGAACAGATGAAATTAACAATAAAACCATCAGGA
TGGACGACTTTTTCCTACCGAATGCGCTTGTAAAAGCTTTGGATGCTGATGCGGTATCAT
CAACCCATCCGCTCTCGTTTAGAGTTGATAAAGCTGCCGAAGTTGTTGAAGCCTTTGATA
GAATTACATATGAAAAAGGAGCGTCCGTTTTGAAAATGCTACAGGCTTTAATTGGACAGA
AGAACTACAAGCAAGCTGTTACGCAATACCTGAAGAAGTTCTCCTACAGCAACGCCCAA
GCTTCTGATCTCTGGAATGTTTTTGACGAAGTTGTCAAGGACGTCAAAGGTCCAGATGG
TAACCTCATGAAAACGACCGAATTCGCATCTCAGTGGACCACTCAGATGGGATTTCCTC
TGGTCACTGTGAAAGCTTTCAACGCAACTTCTTTGCAAATAACACAAACTCGGTATAAGA
CGAACAAAGACGCGCTGGAACCGGAGAAATATCGTCATCCAAAATACGGATTCAAGTGG
GATGTTCCTCTGTGGTATCAGGAAGGCGATAACAAAGATATAAAGTTTGCATGGTTGAC
AAGAGAAAAACCACTCTATTTGCACATGACCAAGCCTGACACCACCATTGTGGTGAATG
CGGATCGCCATGGATTTTATCGACAAAACTATGATGCGAACGGTTGGCGAAAGATCATC
AAACAGCTAAAGAAAAATCATAAGGCATACAGTGCAAGGACGAGAAATGCGATTATAGG
CGATGCATTTGCAGCGGCTCGGATTGATGAACTCGAGTACGAAACTGTATTTGAACTGC
TTGAGTACGCCAAGAATGAAGAGGAATATTTGCCATGGACAGAGGCAATATCTGGTTTT
TATGCCATCTTAGACTTCTTTGGAAATGAGCCGGAGTCGACATCGGCTAAAGCTTTCAT
GAAGAATTTATTGAAGCCGATGTATGACAAAACCAGTATGAAATATATCGCCGACAACTA
CAAAAACGACTCGCTATTCTTTGAAGTGAACCTCCAAACATCTATTATCGATGCTTATTG
CTTCCTTGGAGCTAGGGAATGCATCAAAAATTACGCAGATTTATTTGATAAAGAAGTTAT
GAAGAAATGTAAAGACGGTGACAAAGCAAGCAAATGTGTCAGCATCGCCGCCCCTCTTC

GAGCGAAAGCCTACTGCTATGGCGTGAAAGAAGGCGGAGAAGTTGCTTTCGAGAAGGT
AATGAAACTATGGTATGCAGAAAATGTTCAAGTGGAGAAAGACGTGCTACTCAAAGGAT
TAGGATGTCACAGAGATATTACAGCATTAAAGAGGCTTCTCCTACTAGCACTGGATCGA
AATTCCTCATTTGTTCGACTTCAAGATGTGGCTGCTGTTTATTATGCTGTATCATCGAATC
CCATTGGCAAAGAATTCATGTTCAACTTCCACCTTGAGAGATGGGAAGAAATTCTTGAG
GGCTTGACAACGGAGCACCGAGCTGTTGAACGAGTGATCAAAGCGTGCACTGCAGGTA
TTCGATTGGAGCAGCAGATAGATCAGCTGAGAAGCCTTCAGAAAAATGGTGAACATGCT
CGAGAATATGGAGCATTCGACGGACAAATCGAACGTGCGCAACATAAAATCAACTGGAT
CAAGAAACATATGCGAAAATTATCGGATTTCTTCGAAAAATCTACTCGATGA

*Codon-optimised synthetic DNAs used for molecular cloning*

Seven H11 antigens (KB strain, deduced from DNA sequences) and one apical gut membrane polyprotein GA1 antigen (US strain, AAB01192.1) without the predicted transmembrane domains were codon optimised using GenSmart (GenScript) for expression in *T. ni* cells (Hi5). The synthetic DNA sequences contain a 5'-end flanking region encoding functional motifs, including a melittin signal peptide, a HisFLAG affinity tag, a thrombin cleavage site (in green). To simplify molecular cloning, a 26 bp fragment of *fkp* gene was introduced into the 3'-end (in red).

>*h11_KB strain_codon-optimised*
ATGAAATTCTTAGTCAACGTTGCCCTTGTTTTTATGGTCGTATACATTTCTTACATCTATG
CGGCCCACCATCACCATCACCATGATTACAAGGACGACGATGACAAGCTGGTGCCGCG
CGGCAGCACCTATTACTTTACTCGCAAGGCCTTTGATACATCCGAGAAACCTGGCAAAG
ACGATACCGGAGGAAAGGGAAAAGATAACTCCCCTTCTGCAGCGGAATTACTCCTACCG
ACCAATATTAAGCCGCTATCATATGATCTCACAATAAAGACCTATCTCCCCGGGTATGTA
AACTTCCCACCCGAGAAAAATCTGACTTTCGACGGAAGGGTCGAGATAAGTATGGTTGT
GATCGAGCCTACAAAATCGATTGTTTTGAACTCGAAAAAGATAAGTGTAATCCCTCAGGA
GTGTGAATTAGTAAGCGGAGATAAGAAGCATGAAATTGAAAGTGTCAAAGAACATCCGA
GATTGGAGAAAGTTGAGTTTTTACTGAAGAACCAGCTGGAAAAAGATCAACAGATTTTGT
TGAAGGTAGGGTACATCGGATTGATAAGCAACTCACTTGGTGGAATTTATCAGACTACC
TATACCACTCCTAATGGCACTCCGAAAATCGCCGCTGTCTCCCAGAACGAGCCAATAGA
CGCAAGGCGGATGGTTCCCTGCATGGATGAACCTAAGTATAAAGCTAATTGGACCGTGA
CGGTAATTCACCCTAAGGGTACAAAAGCAGTATCCAATGGCATAGAAGTCAATGGGGAT
GGCGAGATCTCAGGTGACTGGATTACATCAAAATTCTTGACTACGCCTCGGATGTCCTC
TTATTTACTAGCGGTTATGGTTAGTGAGTTTGAATACATAGAGGGCGAAACCCGCACTG
GTGTACGCTTTAGAATTTGGTCGCGTCCAGAGGCCAAAAAGATGACTAAACTTGCTTTA
GATTATGGTATTAAATGTATAGAATTTTACGAGGACTTCTTTGACATTAAATTCCCGTTGA
AGAAACAGGACATGATCGCCCTACCAGACTTCAGTGCAGGGGCTATGGAGAATTGGGG
ACTGATAACGTACCGTGAAAACAGTCTGCTCTATGACGATCGATTCTATGCGCCAATGA
ATAAGCAGCGGATTGCCCGAATAGTTGCCCATGAGCTAGCCCATCAGTGGTTTGGGGA
CCTCGCTACTATGAAATGGTGGGACAACCTCTGGCTCAATGAGGGTTTTGCGAGATTTA
CTGAGTTTATCGGCGCTGGCCAAATAACCAAAGACGACGCACGGATGCGGAACTATTTT
TTAATAGATGTTCTCGAGAGAGCGCTCAAAGCGGACAGCGTCGCTTCTTCACACCCCCT
GTCTTTTAGGATAGATAAGGCGGCCGAGGTGGAAGAGGCTTTTGATGATATTACTTACG
CGAAAGGAGCTTCTGTCCTGACCATGCTGCGTGCATTGATTGGAGAGGAGAAGCACAA
ACATGCCGTATCACAATACCTTAAAAAATTCAGCTATTCCAATGCTGAGGCAACAGATCT
CTGGGCGGTTTTTGATGAGGTTGTAACCGATGTAGAGGGTCCCGATGGCAAGCCCATG
AGAACGACGGAATTTGCGTCGCAATGGACAACGCAGATGGGCTTCCCTGTGATAAGCG

TCGCAGAATTCAATTCTACTACACTTAAGTTAACGCAATCACGTTACAAGGCGAACAAGG
ACGCGGTCGAAAAGGAAAAGTATCGGCACCCGAAATACGGGTTCAAGTGGGACATACC
CCTCTGGTATCAAGAAGGGGATAAGAAAGAAATAAAGCGCACATGGCTACGAAGGGAC
GAACCGCTGTACCTTCATGTTAACGATCCGGGGGCCCCTTTCGTCGTGAACGCTGATAG
GTACGGGTTTTACCGCCAAAACCATGACGCCTCTGGTTGGAAAAAAATCATCAAACAGC
TTAAGGACAACCATGAAGTGTACTCACCACGAACGAGAAACGCAATCATCTCCGACGCA
TTTGCTGCTGCGACCACGGATGCTATTGAGTATGAAACGGTGTTCGAACTCTTGAAATA
CGCTGAAAAAGAGACTGAATACTTGCCCTTAGAAATTGCCATGAGCGGCATTTCAAGCA
TTTTAAAGTATTTCGGTACAGAGCCGGAAGCGAAACCAGCTCAGACGTATATGATGAAT
ATCTTAAAACCAATGTATGAGAAGAGCAGCATCGATTTTATCGCAAATAATTATCGTAAC
GATAAGCTTTTTTTCCAAATCAACCTACAGAAAGATGTCATAGACATGTTCTGTGCGCTT
GGCTCGCAAGACTGCAGAAAGAAATATAAGAAGCTTTTCGACGATGAGGTAATGAACAA
ATGTAGAGATGGGCAGGCTGCTACAGAGTGTGAACGAATTGCGGCCCCACTTAGGTCC
TCCGTGTACTGCTACGGAGTAAAAGAGGGAGGGGACTACGCGTTTGATAAAGTCATGG
AACTATACACAGCCGAAACGCTTGCACTAGAGAAGGATTTCCTCAGGTTGGCACTTGGT
TGCCACAAAGACGTGACTGCACTAAAAGGACTGCTGTTGCGTGCACTGGACCGGAACA
GTAGTTTCGTGCGAATGCAAGACATTCCGTCGGCTTTCAATGATGTGGCGGCGAATCCC
ATTGGTGGAGAGTTCATTTTTAACTTTCTTATCGAACGATGGCCAGACATTATAGAATCG
ATCGGTACAAAGCACACCTACGTTGAAAAGGTCATACCGGCCTGCACCTCGGGTATTCG
ATCTCAACAACAGATCGATCAGTTAAAGAATCTGCAAAAGAATGGCATGAACGCCCGTC
AGTTCGGGGCATTCGACAAAGCCATAGAAAGGGCCCAAAATCGCGTAGACTGGATCAA
GAAGCACTTCCAAAAGCTTGCCGCATTTTTTAAGAAAGCAACGTTGTGACTTATGATCGT
GATACTTGGAATCCC

*>h11-1_KB strain_codon-optimised*
ATGAAATTCTTAGTCAACGTTGCCCTTGTTTTTATGGTCGTATACATTTCTTACATCTATG
CGGCCCACCATCACCATCACCATGATTACAAGGACGACGATGACAAGCTGGTGCCGCG
CGGCAGCCGAAAAGCGTTTGATACGACTGGTGGTAATGGTAAGGAGGACCAACCGATT
GTAGATGATAATTCGCCTAGTGCTGAGGAATTGCGCTTGCCAACCACCATTAAGCCGCT
CACGTATGATTTAGTGATTAAGACTTATCTCCCTAACTACGTCAATTATCCCCCAGAAAA
GGATTTCGCTATAGATGGTACAGTTGTTATCGCAATGGAAGTTGTAGAACCCACTAAATC
TATAGTGTTAAACTCGAAAAATATTAGCGTAATAGCAGACCAATGCGAGCTTTTCAGTAA
CAATCAAAAACTGGACATAGAAAAGATTGTGGACCAGCCGAGGCTGGAGAAAGTAGAGT
TTGTGCTGAAGAAAAAGTTGGAAAAAAATCAGAAAATCACGCTCAAGATAGTCTATATTG
GATTGATCAATGATATGTTAGGGGGACTGTATAGGACCACATATACCGATAAGGACGGG
ACGACTAAAATCGCCGCCTGCACCCACATGGAACCAACCGACGCTCGCTTGATGGTAC
CGTGTTTCGATGAACCTACTTTCAAAGCCAATTGGACGGTCACGGTTATCCACCCCAAG
GGCACAAGTGCAGTCTCAAATGGCATAGAAAAAGGCGAGGGTGAGGTCTCCGGCGACT
GGGTCACCACACGATTTGACCCCACTCCAAGAATGTCCTCGTATCTAATTGCCCTCGTC
ATCTCAGAGTTCAAATATATAGAGAACTACACGAAGTCCGGAGTTCGATTCCGAATATGG
GCACGACCGGAAGCAATGAAGATGACAGAGTACGCTATGATCGCAGGGATAAAGTGTC
TTGACTACTACGAAGATTTCTTCGGAATAAAATTTCCTCTACCAAAGCAAGATATGGTCG
CGTTACCAGACTTCTCGAGCGGGGCAATGGAGAACTGGGGACTCATAACCTATAGAGA
GGGCTCGGTCCTATATGATGAGAACCTGTACGGACCGATGAATAAGGAGAGGGTTGCG
GAGGTCATCGCGCATGAATTGGCTCATCAGTGGTTTGGTAACTTAGTAACAATGAAATG
GTGGGACAACCTCTGGCTGAATGAGGGATTCGCGAGTTTTGTGGAATACATCGGCGCC
GATTTTATAAGTGATGGGTTATGGGAGATGAAGGACTTTTTTCTCTTGGCTCCTTACACA
TCTGGAATTACAGCTGACGCAGTCGCCAGTTCGCACCCGCTCTCCTTCCGTATCGACAA
AGCCGCGGATGTGTCAGAGGCTTTCGATGACATTACGTATCGGAAAGGGGCTAGTGTA
CTACAAATGCTGTTGAATCTAGTTGGGGACGAGAATTTCAAGCAGAGCGTGAGCCGCTA
CCTCAAGAAATTTAGCTACGATAACGCTGCCGCAGAAGATCTCTGGGCGGCTTTCGACG

AAACCGTGCAGGGTATCACCGGACCTAACGGGGGGCCCCTAAAAATGTCTGAATTTGC
TCCTCAATGGACCACCCAGATGGGCTTCCCTGTATTGACTGTAGAGAGCGTTAACGCCA
CGACTTTGAAAGTTACGCAAAAACGGTACCGGCAAAACAAAGATGCAAAAGAACCCGAG
AAATATCGGCACCCCACGTATGGTTTTAAATGGGACGTACCCCTCTGGTATCAGGAAGA
TGAACAGCAGGTGAAGAGAACATGGTTAAAACGGGAAGAGCCGCTATACTTCCATGTTA
GTAACTCAGATTCAAGTGTGGTGGTCAATGCGGAGCGGCGTGCGTTTTGTCGCTCAAAC
TATGATGCCAACGGATGGCGTAACATTATGAGACGTCTCAAGCAAAACCACAAGGTTTA
CGGTCCAAGGACACGCAATGCTTTAATTTCCGATGCCTTCGCTGCGGCAGCAGTAGAG
GAAATGAACTATGAGACAGTTTTCGAAATGTTAAAGTACACGGTGAAAGAGGAAGACTA
CCTACCGTGGAAAGAAGCGGTCTCTGGTTTTAATACCATTTTGGATTTCTTTGGTTCGGA
ACCTGAATCACAGTGGGCCAGCGAATATATGAGAAAACTAATGAAACCGATTTACGACA
AGTCTTCCATCAAATTTATCGCTGAAAACTATAAAAAGGACTCGCTGTTCTTTAAAAACAA
TCTACAAATAGCAGTAATCGATACGTATTGTGGACTAGGCGGTAAAGAGTGCCTTGAGG
AGATGAAAAAGTTGTTCGATAAGGAAGTTATGAAGTGCCAACCCGGTCAGCAGGCGACT
GACTGTGTGAAAGTGACTGCGCCCTTACGGAAGACTGTCTATTGTTACGGCGTTCAAGA
AGGAGGAGACGAGGCCTTTGACAAGGTCATGGAGTTATACAATGCGGAGCAGGTTCAA
CTTGAAAAGGATTCGCTTCGAGAGGCGCTTGGGTGCCATAAGGACGTTACAGCACTGA
AGGGCCTTCTCATGCTTGCCCTTGACCGAAATAGCTCGTTCGTTAGGTTACAAGGCGCC
CATGACGTGTTTAATATCGTATCTCGCAATCCAGTCGGGAACGAACTGCTCTTTAATTTT
CTTACGGAGCGTTGGGAAGAAATCTTGGAGTCTCTCTCCATAAGGCATCGTTCCGTAGA
TCGCGTGATTCGTGCATGCACTAGGGGACTGCGATCACGCGAGCAAGTACAACAACTT
AAGAATCTATACAAGAACGATAAAAGAGCCAGGGAATATGGCGCATTCGGGGGGGCAA
TAGAGAGAAGTGAACACAGAGTGAAATGGATTGAAAAACATTTTCGAAAGCTTGCGGCC
TTCTTTAAGAAGTCAAACTCTTGACTTATGATCGTGATACTTGGAATCCC

*>h11-2_KB strain_codon-optimised*
ATGAAATTCTTAGTCAACGTTGCCCTTGTTTTTATGGTCGTATACATTTCTTACATCTATG
CGGCCCACCATCACCATCACCATGATTACAAGGACGACGATGACAAGCTGGTGCCGCG
CGGCAGCACGTATTACTTTACCCGAAAGGCTTTCGACACCACACAAAAAGAGCAGAAGG
ACGACACTGGAGGTAAGGAGAAAGATAACAGTCCGTCCGCTGAAGAGCTACTTCTACCA
ACCAAGATCAAACCAGTTTCTTACGACTTAAGCATCAAAACTCATTTGCCGGGCTATGTC
AATATCCCCCCGGAAAAAAATTTAACCTTTGATGGCTACGTAGAGATAAGTATGGTAGTG
GTCGAACCGACGAACTCTATTGTTCTGAATAGCAAGAAAATTACATTGGCACAAGGCGG
GTGTGAGCTATTCTCAGGTAACCAGAAACTCGATATAGAGTCTGTCAGGATGCGGGAAC
GCCTTGACAAATTAGAAATTACCTTAAAGAATCAACTTCAAAAAGAGCAAAAAATACTGTT
AAAAATCACATATACCGGTTTGATATCGAATACACTCGGTGGTCTCTACCAAAGTATATA
CACGGACACAGATGGTACGACTAAGATTGTCGCGGTTTCCCAAAACGAGTCCTCGGAT
GCCCGTCGCATTGCCCCATGTTTTGATGAGCCTAAGTATAAGGCCAAGTGGACGGTAAC
AGTAGTGCACCCAAAAGGAACAAAAGCCGCCTCGAACGGAATAGAAGCGAATGGCAAC
GGTGAGCTGCAGGGTGATTGGATAACCTCCAAATTCAAGACTACCCCCCCCATGTCCTC
GTACCTGCTAGCGATTATCGTCTGTGAGTTCGAATACATCGAGGGAAAAACTGAGACCG
GGGTCAGGTTTAGGATATGGTCTCGCCCGGAGGCGAAAGCGATGACGGCCTATGCGCT
CGACGCCGGGATAAGGTGCCTCGAATTTTATGAAAAATTCTTCGACATCAAATTCCCTCT
AGAAAAGCAAGATATGATAGCTCTACCAGACTTTACAGCTGGCGCAATGGAGAACTGGG
GACTAATCACTTATCGAGAGGATAGCCTCCTTTATGACGAGAAGATTTATGCGCCTATGA
ATAAGCAACGGGTAGCTCTAGTGGTCGCCCATGAACTCGCCCATCTATGGTTTGGGAAT
TTGGTCACATTGAAATGGTGGGATGACACATGGTTAAACGAAGGTTTCGCCACGTTTGT
AGAGTATCTCGGTATGGACGAAATTAGTCACAACAACTTTCGTACTCAGGATTTTTTCCT
TTTAGACGGCATGGATCGTGGGATGCGAGCAGATAGCGCGGCGTCATCACACCCATTG
TCCTTCAGAGTAGACAAGGCAGCGGAAGTTGCGGAGGCTTTCGACGACATATCTTACG
CCAAGGGAGCTTCGGTTCTCACGATGTTGCGAGCATTAATCGGTGAAGATAACTACAGG
AATGCAGTCGTACAGTATCTGAAAAAATTCAGTTACAATAATGCGCAAGCTGCGGACCTT

TGGAATGTATTCAATGAGGTTGTCAAGGGCGTCAAGGGCCCTGATGGGAATGTGATGAA
GATTGACCAGTTCACCGACCAATGGACTTACCAAATGGGGTATCCCGTGGTTAAAGTCG
AAGAATTCAACGCGACAGCTCTTAAAGTGACACAGTCGAGATATAAAACTAATAAGGAC
GCTTTAGAGCCAGAGAAGTATAGAAACCCGAAATATGGCTTCAAATGGGACGTACCCCT
ATGGTATCAGGAGGGAAATAGCAAGGAGGTTAAGAGAACTTGGCTCAAGCGGGATGAA
TTATTGTACCTGAACGTGAACAACAGGGACACTTCGCTGGTTGTGAATGCCGATCGGCA
CGGATTCTATCGTCAGAACTATGACGCCAACGGATGGAAGAAGATCATCAAGCAACTAA
AGGAAAATCACGAGGTATTTGGCCCTAGAACAAGGAACGCAATTATTTCAGATGCATTC
GCGGCAGCTACGATAGATGCCATAGATTATGAAACTGTCTTCGAGTTACTCGAATACGC
GAAAAACGAGGAAGAGTTTCTTCCGTGGAAGGAAGCTTTGTCCGGGATGTTCGCAGTTC
TGAAATTTTTTGGGAATGAGCCCGAAACTAAACCTGCACGCGCATATATGATGTCCATTC
TTGAGCCTATGTACAACAAAAGCAGCATAGATTACATCGTTAAGAATTACTTGGATGACA
CACTGTTTACGAAGATCAACACCCAGAAGGATATCATTGACGCTTACTGCTCTCTAGGTA
GCAAAGACTGCATTAAGCAATACAAGGACATTTTTTACGATGAAGTGATGCCAAAATGCA
AAGCGGGAGAAGCTGCAACGAAATGCGTGAAGGTATCCGCTCCTCTAAGAGCAAACGT
TTACTGTTATGGGGTGCAGGAGGGCGGGGAGGAAGCATTTGAAAAGGTGATGGGGCTG
TACCTCGCCGAGGATATACAATTAGAAAAGGGCATCCTTTTTAAGGCGCTCGCATGTCA
TAAGGATGTAACGGCTTTGAAAGAACTTCTTCTTCGTGCGTTGGATCGTAAGTCATCTTT
TGTCCGCCTACAGGATGTCCCGACCGCGTTCCGAGCAGTTAGTGAGAATCCCGTGGGC
GAGGAATTTATGTTTAATTTTTTAATGGAGCGATGGGAAGAAATTACGGCCTCGCTGGAA
ACTGAACATCGAGCAGTAGATAAAGTAGTAGGAGCATGTTGTACCGGGATCCGGTCTCA
ACAGCAGATCGACCAGCTTAAGAACCTTCAGAAAAATAATGCCCAGGCTAAAAAATTCG
GTAGTTTCACGCAAGAAATCGAGAAGGGAGAACACAAAATTGCATGGATAAAGAAGCAC
TTCCATCGGTTGAGCGAATTCTTTAAACGGGCTCGTAGTTGACTTATGATCGTGATACTT
GGAATCCC

*>h11-4_KB strain_codon-optimised*
ATGAAATTCTTAGTCAACGTTGCCCTTGTTTTTATGGTCGTATACATTTCTTACATCTATG
CGGCCCACCATCACCATCACCATGATTACAAGGACGACGATGACAAGCTGGTGCCGCG
CGGCAGCACATACTTTTTTACACGCAAAGCTTTCGACCCGACGCAGAAGGATAAGAACC
AGCCCGGCGGGAAAGAGAAAGACAACTCCCCCTCGGCTGCCGAGTTGTTACTTCCGAG
CAATATAAAGCCATTGTTGTATGATCTGACAATCAAGACATATTTGCCAGGGTATGTAAA
CTTTCCACCGGAAAAAAATCTAACTTTCGACGGGCGTGTGGAAATATCGATGGTGGTCG
TCGAACCCACGAAATCTATAGTCCTAAATGCGAAAAAGATTACAGTTATTCCCGCAGAAT
GTGAGGTGTTGTCAGGTACTCAAAAGCTTGACATAGAGTCCGTCAAGGAGCATGAACG
GCTGGAGAAATTAGGATTCCGACTAAAAAGCCGCCTGGAAAAAGACCAAAAGATACTCC
TTAAGATCACTTATGCAGGGCTGATATCGAACACACTTGGTGGAATATACCAAACCACGT
ACACCGACGCCAATGGTAATCCAAAAATAGCGGCTGTCAGCCAAAATGAGCCAATAGAT
GGGCGCAGGATGGTCCCCTGTATGGATGAACCTAAGTACAAAGCTAATTGGACCGTTAC
GGTTATTCACCCCAAGGGAACCAAGGCTGCCTCCAACTCTATCGAAATCAATGGCGAGG
GTGACGTGTCTGGTGATTGGATAACTTCGAAATTCGAAACTACGCCTAGAATGTCCAGT
TATCTATTAGCGGTTTTCATATCAGAATTTGATTTTGTAGAGGGACGCACCAAGCAGGAC
GTACGGTTCCGAATTTGGTCACGGCCCGAGGCGAAGGGGATGACGAAATACGCTCTGG
AGAGTGGGATCAAATGCATCGAATTCTACGAAGATTTCTTCGACATAAAGTTTCCTTTAA
AAAAGCAGGATATGATAGCTCTTCCTGACTTCAGCGCGGGTGCCATGGAGAACTGGGG
TTTGATCACCTATCGTGAGAATTCGCTTTTATATGACGAAAAGTTCTATGGTCCAACAAA
CAAGCGTCGCGTGGCCGTCGTGGTTGCTCATGAATTGGCACACCAGTGGTTTGGCGAC
TTAGTCACCATGAAATGGTGGGACGATTTATGGTTGAATGAAGGATTTGCCACATTCGTA
GAATATATTGGGGCGGATGAAATCGGCGATCACTACTTTAACATGCCTGATTTCTTCCTC
ATTGGCGCACTGGAACGCGCTCTTAAAGCCGACAGTGCGGCATCCTCCCATCCGCTAT
CCTTCAGAATTGATAAAGCCGTTGAAGTCGAGGAGGCCTTTGACGACATTAGCTATGCC
AAAGGAGCATCCATAATTACTATGCTCCGAGCGCTAATAGGCGAAGACAAGCACAAACA

TGCTGTTACTCAATATTTGAAAAAGTTCTCCTATTCTAACGCGCAAGCGTCTGATCTCTG
GGAGGTATTCGACGAGGTCGTTACCGATATCAAGGGACCTGATGGTAAACCTATGAAGA
CGACCGCATTTGCTGACCAATGGACGACCCAAATGGGTTTCCCCCTCGTGACTGTTGAA
GCTTTTAATGCTACCTCGGTAAAAATCTCTCAGAGTCGTTTTAAGACTAATAAAGATGCT
AAAGAGCCGGAGAAGTACCGGCATCCTAAATATGGTTTCAAATGGGACATACCTCTATG
GTATCAAGAGGGCGACAACAAGGAGGTCAAGCAAACGTGGATACGAAGAGAAGAGCCA
CTGTACCTCCACGTGAATGATCTCAGTAAACCTTTTGTAGTCAATGCAGACCGGCATGG
GTTCTATCGGCAGAATTACGATGCTGACGGTTGGCGTAAGATCATTAAACAGCTCAGGG
ACAACCACAAGGTGTTTAGTCCAAGGACAAGAAATGCAATTATCAGTGACGCATTTGCT
CTGGCGTCGGTTAACGCGATTGAATACGAAACAGTGTTCGAATTGCTAAAATACGCAGT
GAACGAGGAGGAGTTTATACCGTGGACGGAAGCCATTTCAGGCATCTTTGCTGTGTTAA
AGTTTTTCGGAAACGAGCCGGAAAGCAAACCGGCCGAGGCGTACATGATGAAGATTCTT
GAGCCAATGTACAAGAAAAGCGATCTGGGATACATTGCCGCGAAGTACAAAGTCGACCA
GCTATTCTCGAAGATTAATCTGCAGAAGGATATAATCGACGCGTACTGCGCACTTGGGA
GCAAAGACTGTATGAAAAAATATAAAGACATCTTTGATAGGGAAGTAATGAACAAGTGCA
ATGATGGAGATGAGGCAACTAAGTGCGTAAGTGTAGCAGCTCCACTACGTAGTTCAACC
TACTGTAATGGAGTGAAGGCGGGCGGGACGTATGCATTTGAGAAAGTTAAGGCACTTTA
CTACAAAGAAACAGTTCAGCTTGAAAAAGATATGTTACTTCGAGCCCTGGGGTGTCACA
GAGACGTAACTGCACTCAAGGGATTATTGCTGCTCGCCGTTGATAGGAACTCATCTTTT
GTACGCTTGCAAGATATACCAAACGCCTTCCAAGCAGTAGCGGCTAACCCGGTCGGTG
AAGAATTCATGTTTAACTTTCTTATCGAGCGATGGGGCGATATTATTGGTTCAATTGGGT
CAGAACCCACCTATGTTGAACGAGTCATTCCGCCCTGCACTTCAGGCATCAGGAGCAAG
CAGCAAATCGACCAATTACGTAACCTACACAAAAATGGCATCCATGCCCAGGAATATTC
GACATTTGTAAAAGAGATTGAGAGAGCCGAGCACAAAGTGGATTGGATCAAAAAACATT
TCAAGAAGCTAGCCTCTTTTTTCAAGAATGCGACGTGGTGACTTATGATCGTGATACTTG
GAATCCC

*>h11-5a_KB strain_codon-optimised*
ATGAAATTCTTAGTCAACGTTGCCCTTGTTTTTATGGTCGTATACATTTCTTACATCTATG
CGGCCCACCATCACCATCACCATGATTACAAGGACGACGATGACAAGCTGGTGCCGCG
CGGCAGCCAGTGGACTAAACGAACAGTTTTAAAGTTTACCCCAATAACCCTACTGCTAC
TTTTGTTTTTGGTAGCCGCTTCTATAGGTCTATCTATTGGCCTGACCTATTACTTCACTCG
TAAGGCCTACGACACAACAGAGAAGAATAAGGATCACGGGGCCGATGATAACAGCCCC
AGCGCCGAGGAGCTTCGTCTCCCAAAAAATATCGAGCCGCTACTGTACGATCTTTCTAT
TAAGACATACCTACCGGGGTATGTTTCATTTCCGCCTGAGAAAAATTTAACATTTGATGG
CCAAGTTGGGATTAGTTTGCGGGTGGTGGAACCTACCAAGTCCATTGTTCTCAACTCCA
AAAATATCACCGTCATTCCCGACAAATGCGAGTTATTCTCGGGAGATAAGAAGTTAGAAA
TCGAGTCGATTAAGGAACACGAAAGACTCGAAAAGCTAGAGATCCTGCTGAAGAACAGA
TTGGAGAAAGACCAAGAGGTGCTACTCAAGATAATATACACTGGTATCATATCCAACAC
GCTCGGAGGACTTTACCAAGCTACGTATACTGACACAGACGGGACCGTGAAGATTGCC
GCTGCTTCCCAAAATGAGCCAACTGACGCAAGACGTATGGTTCCCTGTCTGGATGAACC
TTCTTTCAAAGCGAACTGGACAGTCACTGTTATACATCCCAAGGGGACGAAGGCAGTAA
GTAACGGCATTGAAACAAATGGAGAGGGTGAGGTCTCCGGGGACTGGATCATTAGTAA
GTTTGAGACGACCCCTAGGATGTCGAGTTATTTACTAGCGGTTGTCGTCTCTGAATTTGA
CTATATCGAGGGTTTTACTAAATCGGGTGTACGATTTCGAATATGGTCCCGGCCCGAAG
CAATGAACATGACGGGGTATGCAAAAGACGCGGGAATCAGGTGCCTGGAATATTACGA
GAATTTTTTCGACATAAAATTCCCTCTCAAGAAGCAGGATATGGTGGCATTGCCCGACTT
TTCTGCGGGGGCTATGGAAAACTGGGGGCTGATCACCTATAGGGAAAATCAATTGTTGT
ACGACGATCGATATTACGGCCCAATCAACAAGCAGCGCGTTGCATTAGTGGTTGCGCAC
GAGCTGGCGCACCAGTGGTTTGGCAACTTAGTGACTTTAAAATGGTGGGATGATACGTG
GCTCAACGAGGGGTTCGCGACGTTCGTGGAATATATTGGCACGGACGAAATATCATATA
AGAATTTCCGGATGGACGACTTTTTCCTCCCAGACGCATTGGCGATAGGCCTTGACGCC

GACGCTGTAAGTTCAACTCATCCTTTGAGTTTCCGAGTGGATAAGGCGGCGGAGGTGG
CGGAAGCCTTCGATGAGATTACATATGCTAAGGGGGCATCAGTATTGGTCATGCTTCAG
GCGGTAATTGGTGAAAAAAATTACAAACAGGCCGTGACCCAGTATTTGAAAAAATTTAGC
TATTCAAATGCTCAAGCTACAGATCTATGGAACGTTTTCGATGAGGTGGTCAAGGACGT
AAAAGGTCCGGATGGCAACTTAATGAAAACGACCGAGTTTGCCTCCCAGTGGACCACAC
AGTTAGGATTCCCTTTGGTTACGGTGAAAGCGTTCAATGCAACGAGCTTGCAGATCACG
CAAACCCGCTATAAAACCAATAAGGATGCCCTTGAACCAGAGAAATACCGACACCCTAA
ATATGGATTTAAGTGGGATATTCCGTTGTGGTACCAAGAGGGCGATAATAAAGATATCAA
ACTAGCTTGGCTAACACGTGAAGAACCACTTTACTTACATGTATCGAATCCGGACACAA
GCATAGTTGTTAATGCCGACAGGCATGGTTTTTACAGGCAAAACTACGACGCAAACGGG
TGGCGTAAGATAATAAAGCAGCTCAAAGAAAACCACAAGGCCTACAGTACTAGAACTCG
GAACGCTATTATCGGAGACGCTTTTGCTGCAGCACTGATCGACCAGCTGGAATATGAAA
CGGTCTTCGAGCTACTCGAGTACGCGAAAAACGAGGAAGAATATCTGCCATGGACTGAA
ACAATTTCAGGTTTCTACGCAATACTCGACTTTTTTGGTAACGAGCCCGAATCAACCAGT
GCTAAGACGTTTATGATGAATATCCTCAAACCGATGTATGAAAAAACGTCTATGAAGTAC
ATCGCTGACAACTACAAGAACGATTCGCTGTTCTTCGAAATGAATTTGCAAAAATCTGTC
ATAAACGCATATTGTTTTTAGGCTCAATGGAGTGTATTAAAAATTACACGGATCTGTTTG
ATAAGGAGGTAATGAAAAAGTGTAAAGATGATGACGAAGCGTCGAAATGCGTATCAATA
GCGGCGCCGTTACGCGCTAAGGCTTATTGCTATGGAGTCAAGGAAGGAGGACAAGTAG
CGTTCGATAAGGTAATGAAGCTGTACTATGCAGAAAATGTCCAATTAGAGAAGGATGTC
CTCCTTCAAGGGCTCGGCTGCCACAAAGATATAACAGCTCTAAAAAGGCTTCTCCTTCT
AGCACTCGACAGAAATTCTTCTTTTGTACGCCTACAGGATGTAACCGACGTATATGATGC
CGTGTCGTCGAACCCCGTTGGTGAGGAGTTCATGTTCAATTTCCTTCTGGAGCGTTGGG
AAGAAATTCTTGAAAGCTTAACAACTGAGCATCGGGCAGTCGAACGCGTAATTGAGGCC
TGTACTGCCGGCATTCGGTCCGAACAGCAAATAGACCAGCTTCGAAGCCTCCAGAAAAA
CGGTGCCCACGCCAGAGAATACGGTGCCTTCGACGAGCAAATCGAACGCGCAGAGCAT
AAGATCAATTGGATAAAAAAACATCTTCGCAAGTTAAGTGATTTTTTCGAAAAATCCACTT
TCTGACTTATGATCGTGATACTTGGAATCCC

*>h11-5b_KB strain_codon-optimised*
ATGAAATTCTTAGTCAACGTTGCCCTTGTTTTTATGGTCGTATACATTTCTTACATCTATG
CGGCCCACCATCACCATCACCATGATTACAAGGACGACGATGACAAGCTGGTGCCGCG
CGGCAGCACCTACTTTACAAGGCGCGCTTATGATACTACGGAGAGGAACAAAGATTATT
GCGCTGATTATAACTCCCCCTCTGCTCAGGCTCTTAGGCTGCCGAGGAATATTGAGCCG
TTGCTGTACGATTTATGCATTAAGACTTATCTACCTAACTATGTAGATTTCCCCCCAGAAA
AAAATTTAACTTTCGATGGTCAAGTCGGCATCAGTCTGAGAGTCTTAGAGCCTACGAAGT
CCATTGTGCTCAATGCGAAAAATATTACAGTGATACCTAATAAGTGTGAGCTCTTCCTAG
GAAACCAGAAGTTGAAAATTGAGAGCGTGAAAGAGCACGAGAGACTGGAAAAAGTTGAA
TTTCTTCTTCGCAACTGGTTGCAGAAAGACCAGAAAGTCCTCCTCAAAATCGTTTACACG
GGCTTAATATCGAACACCCTGTTTGGCCTTTACCAGGCAACGTATACCGACCCAGATGG
GACAGTAAAGATAGCTGCCGCAACCCATATGGAGCCAACAGCGGCAAGACGAATGGTC
CCGTGTCTGGACGAACCTTCTTTCAAGGCGAATTGGACGGTTACCGTGATTCATCCCAA
GGGGACTAAGGCAGTATCAAATGGAATAGAGACAAATGGAAAAGGCGAGGTGAGCGGG
GACTGGATTATATCGAAATTTGAGACTACACCACGTATGTCCTCGTATCTTCTAGCCGTT
GTAGTCTCAGAGTTCGATTACATAGAGGGCTTTACTAAGACTGGCGTACGATTCCGCAT
CTGGTCCCGGCCAGAGGCCAAAAACATGACCGAGTATGCCCGGGACGCGGGGATAAG
GTGCCTGGAATTCTACGAAAACTTTTTTGAATCAAAGTTTCCACTGAAGAAGCAGGATAT
GGTAGCCCTTCCAGACTTTTCGTTTGGGGCGATGGAGAATTGGGGTCTCATCACGTACC
GCGAGACTAGTTTGCTATACGACGACCGTTACTATGGTCCCATCAATAAACAAAGAGTC
GCATTGGTTGTTGCGCACGAGTTGGCTCATCAGTGGTTCGGGAACCTGGTTACGCTGA
AGTGGTGGGATGATTTATGGCTAAATGAAGGCTTCGCTAGACGGGTGGAATATATTGGT
ACCGATGAGATCAATAACAAAACGATTCGTATGGATGATGTATTTCTACCGAATGCCCTC

GTAAAGGCGTTGGATGCTGACGCGGTTAGCTCGACTCACCCACTTTCCTTTCGCATAGA
TAAAGCTGCGGAGGTATTAGAAGCGTTCGATACGATTACTTACGAAAAAGGCGCCAGTG
TCCTAGAGATGTTGCGAGCAGTCATAGGAGACAAGAATTTCAAACAAGCCGTGACGAAG
TATTTGAAAAAATTTTCATATCGAAATGCTAAGCCCAGCGACCTCTGGAATGTTTTTGAC
GAAGTCGTGAAGGATGTACGAGGGCCGGATGGTAACTCTATGAAGACAACCAAATTTGC
CCCTCAGTGGACCACTCAGTTGGGGTTTCCCCTGGTCGCGGTGAAGGATTTCAATGGG
ACCTCTGTTAAAATCACACAAACCAGGTACAAAGCCAACAAGGATGCGCTTGAACCCGA
GAAATACCGTCACCCTAAATATCGATTTAAATGGGACATTCCGCTGTGGTATCAGGAGG
GTGACAACAAGGACATTAAATTCGCGTGGCTAACACGAGAGAAGCCTTTGTACCTACAT
AAGACTAAGCCTGACACGAGCATTGTGGTGAACGCTGATCGCCATGGTTTCTACCAACA
AAATTACGACGCAAAGGGATGGCGGAAGATAATCAAGCAGCTTAAGAAAAACCACAAGG
CATATAGCGCCCGGACTCGCAATGCTATTATAGGAGATGCATTTGCAGCCGCGCTCATA
GACGAGCTTGAATACGAGACAGTTTTTAAACTACTTGAATATGCCAAGAATGAGGAAGAA
TATTTACCCTGGACGGAAACACTCAGTGGGTTCGACGCCATACTGAACTTCTTTGGTAAT
GAGCCAGAATCTACCTCTGCCAAGGCATTCATGAAAAACATTCTCAAACCTATGTACAAA
AAGACCTCCATGAAGTATATAGCAGTTAACTACAAAAATGACAGTCTGTTCTTTGAAGTC
AACCTTCAAACGTCTATCATCGACGCATACTGTCATTTAGGTGCTAGAGAATGCATAAAA
AACTATACAGATCTCTTCGATAAAGAGGTAATGAAAAAGTGCCGTGACGGCGACAAGGC
TAGCAAATGCGTGTCCATTGCTGCACCGCTACGGGCAAAAGCCTACTGTTACGGAGTTA
AAGAAGGTGGCGAAGTAGCGTTCGACAAGGTTATGAAGCTGTGTTACGCGGAGAACGT
GCAAGTCGAAAAAGACGTGTTGTTAAAAGGATTGGGGTGCCATCGTGATATCACTGCTT
TAAAACGGCTATTGCTATTAGCATTAGATAGAAATTCAACCTTCGTAAGGCTTCAGGACG
TCGCCGCCGTCTATGACGCGGTATCGGCAAACCCGATCGGAAAGGAGTTCATGTTCAA
TTTTCTATTAGAAAGATGGGAGGAAATCCTTGAATCGCTCACCGCGGATCACAGGACGG
TTGAAAGGGTGATCAAAGCCTGTACAGCAGGCATCCGTCTAGAGCAGCAAATAGACCAA
CTCCGTAGTCTCCAAAAAAATGGAGAACATGCTCGAGAGTATGGTGCATTTGATGGACA
AATAGAACGGGCGCAACACAAGATCAACTGGAACAAGAAGTGACTTATGATCGTGATAC
TTGGAATCCC

*>h11-5c_KB strain_codon-optimised*
ATGAAATTCTTAGTCAACGTTGCCCTTGTTTTTATGGTCGTATACATTTCTTACATCTATG
CGGCCCACCATCACCATCACCATGATTACAAGGACGACGATGACAAGCTGGTGCCGCG
CGGCAGCGCAGATGACTCGTCACCCTCAGCCCAAGAATTACGCCTACCGCGAAATGTT
GAGCCTCTGCTTTATAATCTATCGATTAAGACTTATTTGCCGAGTTACGTCAACTTCCCG
CCGGAGAAAAATTTGACATTTGATGGGCAAGTCGGCATTAGCCTTAGGGTAGTTGAACC
GACCAAGTCCATTGTCCTTAACGCTAAGAATATTACCGTGATACCTAATAATTGCGAGCT
CTTCTCGGGCGGTAGGAAATTAGAAATTGAATCAGTTGAAGTGCACGAGCGTCTAGAGA
AAGTGGAATTCCTCCTACGGAATCGTCTCCAAAAAGATCAGAAGGTTCTACTAAAGATAA
TATATACTGGCGTAATATCCAACAGTCTTTTTGGTCTATATCAGGCAACGTATACAGATG
CGGACGGAAATGTCAAGATCGCAGCAGCGACTCAGTTGTGTCCCAGCGATGCCAGAAG
GCTTGTTCCATGTCTGGACGAACCTAGTTTTAAAGCGTCATGGACCGTGACCGTGATTC
ATCCCAAGGGTACCAAAGCTGTATCGAACGGAATAGAGACTAACGGGAAGGGGGAGGT
GTCTGGTGATTGGATAATCTCGAAGTTTGAAACCACTCCTCGGATGTCTTCCTACCTGTT
GGCCATAGTAGCCAGTGAGTTCGATTATGTCGAAGGGTTCACCAAAAGCGGGGTAAGA
TTCCGAATTTGGTCCAGACCTGAAGCTAAGAACATGACGGCTTATGCGCGTGACGCGG
GCATTCGGTGCCTTGAGTTCTACGAAAATTTTTTCGACATCAAGTTTCCATTGAAAAAAC
AGGACATGGTAGTGCTACCCGACTTTTCTTTCGGAGCTATGGAGAATTGGGGCTTAATA
ACCTATCGGGAAAATAGGCTTCTCTACGACGATCGGTATTACACCCCCATAAACAAACA
ACTAGTCGCCCTCGTTGTGGCACATGAGTTGGCTCACCAATGGTTTGGAGATTTAGTTA
CACTTAAATGGTGGGACGATCTATGGTTGAACGAAGGATTCGCGCGATTCGTAGAATAT
ATAGGAACTGATGAAATTAATAATAAGACGATACGTATGGACGATTTTTTCCTGCCAAAT
GCACTTGTCAAGGCTCTTGACGCTGACGCTGTAAGCTCTACACACCCACTATCGTTTAG

GGTAGACAAGGCTGCCGAGGTAGTTGAGGCGTTTGACAGAATCACCTACGAAAAGGT
GCGTCAGTCCTCAAGATGCTGCAGGCGTTAATAGGCCAAAAGAACTATAAGCAAGCCGT
AACTCAATATCTAAAAAAGTTTTCCTACTCAAACGCGCAGGCAAGTGATCTCTGGAACGT
TTTCGACGAAGTCGTCAAAGACGTCAAAGGCCCGGATGGTAACTTAATGAAGACGACCG
AGTTCGCGAGTCAGTGGACGACTCAAATGGGTTTTCCACTTGTCACTGTGAAAGCATTC
AACGCAACGAGCCTACAGATCACGCAGACACGTTACAAAACGAACAAAGACGCACTGG
AGCCTGAGAAATATCGCCATCCTAAGTATGGCTTTAAGTGGGACGTACCATTATGGTAC
CAAGAAGGAGATAACAAAGATATTAAATTTGCATGGTTAACGCGCGAAAAGCCCTTGTA
CTTACATATGACAAAGCCGGACACCACAATCGTGGTCAACGCGGATAGACACGGTTTTT
ATCGCCAGAACTATGATGCGAACGGTTGGCGCAAAATCATCAAACAGTTGAAAAAGAAC
CATAAGGCCTATTCGGCCCGAACACGCAATGCAATAATAGGTGATGCGTTTGCCGCGG
CGCGCATTGACGAACTCGAATATGAGACAGTTTTCGAGCTGTTGGAGTATGCCAAGAAT
GAGGAAGAGTACCTTCCATGGACAGAAGCGATCAGCGGGTTCTACGCCATCTTGGATTT
TTTTGGGAACGAACCAGAATCTACTTCCGCAAAAGCTTTTATGAAAAACCTGCTCAAGCC
TATGTACGATAAAACTAGCATGAAATACATAGCAGACAATTACAAAAATGACTCCCTCTT
CTTTGAGGTTAATTTACAGACATCGATCATCGATGCCTACTGCTTCCTAGGCGCAAGGG
AATGTATTAAGAATTATGCTGATCTTTTTGATAAAGAGGTGATGAAGAAATGCAAGGACG
GCGATAAAGCTTCTAAGTGTGTCTCTATCGCAGCACCGCTCCGGGCTAAGGCTTACTGT
TATGGAGTGAAGGAGGGGGGAGAGGTAGCTTTCGAAAAGGTTATGAAATTATGGTACG
CGGAAAATGTTCAAGTAGAAAAAGATGTTCTGCTCAAAGGATTAGGGTGCCACAGAGAT
ATTACCGCACTTAAGAGATTGCTGCTCCTCGCACTAGACCGGAATTCGAGCTTTGTTCG
CCTGCAGGACGTGGCTGCCGTGTACTACGCCGTTTCCAGTAACCCCATCGGAAAGGAG
TTCATGTTCAACTTCCATCTGGAGCGATGGGAGGAAATTCTGGAGGGCTTAACTACAGA
GCATAGAGCGGTAGAGCGAGTCATTAAGGCCTGTACGGCTGGGATCCGTCTAGAACAA
CAAATAGATCAGCTCCGTTCACTCCAGAAGAACGGGGAGCACGCCCGAGAGTACGGTG
CCTTTGACGGCCAAATTGAGCGGGCACAACACAAAATAAATTGGATCAAAAAGCACATG
AGGAAACTGTCAGACTTCTTCGAAAAATCTACGAGGTGACTTATGATCGTGATACTTGGA
ATCCC

*>GA1_US strain_codon-optimised*
ATGAAATTCTTAGTCAACGTTGCCCTTGTTTTTATGGTCGTATACATTTCTTACATCTATG
CGGCCCACCATCACCATCACCATGATTACAAGGACGACGATGACAAGCTGGTGCCGCG
CGGCAGCGAATTGCATTTCCCCGGAGAAACTCACTTGAAAAATGTTAGGCAACTGACTT
TTGAACATCGTAACACCGAGGCTTACTTTAACTCAGACGACACATACCTTGTCTTTCAAG
CTACTGGATATGGGGTGGACTGTGAACAGATTTACCGACTCGAGCTCAGCCGTCCAGT
CGAAACTTTAGCGAAGATATCGACGGGTATTGGGACATCGGCGGCCTCGTTTTTCTACC
CTAACTCAGTGGACATCCTATACTCAGGAAATTTTCACAAGACTCGTGTGAATGCCAAGA
AACTCACCAATGACAGCAGTTGCCCTCAGAAGGTCTGCGACTCGAGTAAAGCTAAAACG
GATCCCATCATAAAGAAGATGTGCGAATCAGGTCATGCCTGGGATGTTTTCCCAACATA
CGATATCTTCAAAGTAAACGAATATGGTAACGTCTTAGACCAAATTACCAAGAATGATGT
ATACGATTCGGAAGCATTTATCAGTCCCGATGGCAAAAAAATAGTTTATACCTCGAAACA
AAGCGGTGATTTGGACTTGTGGATCACAGATATAAATGGAGCCCTTAAAATCCAGCTTA
CGAAGGCAAAGGGTTACGACGGGGGTGCATCATTTAGCCCAGACGGTGAGAAGATTGT
ATTTCATGCTTCCCGCCCGACCACAAAAGACAAGATTGACATCTATGATCACTTATTAGA
AAACGACTTGGTGGCAATGTCTGAGATGGAACTCTACGTCATGAACGCGGATGGTATTG
ATAAACGGTCTGTATTTAAACAGCCGCGGGGGGGCCTGAACTGGACTCCTTATTATCAT
CCGGATAACAAGAGAATAATCTTCTCTTCTAATACTAATAGCACTAAGCCAAGTGAGTTC
CATTTATATATCGTGAACGAAGACGGTACCGGACTTGAGAGAGTTACGTTTGGATCAGG
ATATTTTAATGCTTTTCCGGTATTTAGTCACGATGGCAAGAAAGTTGTTTGGTCTTCAAAT
AGGGGCAGCACTAAGAAAGGCGATCTGAACCTATTCATAGCTGACTGGGTTGATCCAG
GAAGAGACACAGATGATGAGTCTGATGCCAAAGAAGAGCTTAGAACAAATCGGATTCCC
AAAAAGGCCACGCAAAAAAACGCGATGAGGTGGCAAGACACCGTGACAACGCCATACG

ACAACGTAGTCCACTATACGGGCGAAAGACGACTAAAGAACGTCAAACAATTGACATTT
CGGGGCCAAAATGCCGAAGGGTACTTCTCCTATGACGATAGTAAGATTATCCTTCAGGC
AACCGGATACGGAACGACTTGCGACCAAATTTATGAACTCGATTTGAATGTCGACCCGC
GAAAGCAGATAATGAAGCGAATGTCCACCGGGCTGGGCGGCACGACATGTAGTTTCTT
TTTCAATGAGCCCGACAATAACCATCGCCTCTACGCAGGCGATTTCTGGGCGCTGAACG
ATACAGTGCGAGATACGATCATAACTCACACGTGTCCTGCGAAGAAATGCGAAAACAGA
AAGGCTATAAAGGACCCGGTGTTGAAAGAACTCTGTAATACCGTATACACATGGACAT
TTACCCCGAGTTCGACATATTCATGGTGAATAAATACGGGAATATCGTGAAACAACTCAC
CGATGAGCCTGGTTATAATGCGGAAGCTGTTCTCTCCCCGGATGGGAAGACTATCGCG
TTCACTTCCATCAGGACAGGGGACCTAGAACTCTGGACCATGAATACAGATGGGACTAA
CTTACACCAGGTAACGAAGGAGTATGGGTATGATGGTGGTTGTTTTTTTTCTCCCGATG
GGAAACGTCTAGTGTTCCGCGCCTCCCGTCCCAAGACCCAGAAAGAGAAAGAGAAGTA
TCAGAAGCTTTTAGACTATGGATTAGTCGAGCCTACACTAACGGAGATATACGTGGTCG
ATGTCGACGGGAAGAACCTAAAGCAGGTAACAAACTTCGGTGTTGCCTCGTGGGCACC
GTATTACCTTCCAGACAACAAACGCATTATTTTCTCCTCCAATTATAACATGAGCGCAAA
GCAGTTCGGATCGTTTGCCTTGTATGTTATAAACGAGGACGGTACCGGCCTGGAAAGG
GTAACCTTTGGGGAGGGGTACCAATTCAACGCATTTCCTATGTTCAATCGAGCAGGTAA
TAAGCTGGTTTGGGGCAGCTCACGGAATCGCACGAAAGGAGCGTCTCTGAATCTATTCA
TTGCTGATTGGGTTGACAAAATAGATGATGATAACATTGGAGGAGGCGGAGGTGGCGA
CGATAACAAAAAAAAAGACGAGAAGGGCAAAGAAAAAGAGGAAAAAAAGAGTTGACTTA
TGATCGTGATACTTGGAATCCC

*Protein sequences of recombinant products*

Based on our design, the recombinant proteins should have the following sequences and corresponding theoretical isoelectric points (pI) and molecular weights (Mw). Protein primary sequences were verified by LC-MS/MS using purified antigens (**S1 Table**).

>Recombinant product H11_KB strain
MKFLVNVALVFMVVYISYIYAAHHHHHHDYKDDDDKLVPRGSTYYFTRKAFDTSEKPGKDD
TGGKGKDNSPSAAELLLPTNIKPLSYDLTIKTYLPGYVNFPPEKNLTFDGRVEISMVVIEPTKS
IVLNSKKISVIPQECELVSGDKKHEIESVKEHPRLEKVEFLLKNQLEKDQQILLKVGYIGLISNS
LGGIYQTTYTTPNGTPKIAAVSQNEPIDARRMVPCMDEPKYKANWTVTVIHPKGTKAVSNGI
EVNGDGEISGDWITSKFLTTPRMSSYLLAVMVSEFEYIEGETRTGVRFRIWSRPEAKKMTKL
ALDYGIKCIEFYEDFFDIKFPLKKQDMIALPDFSAGAMENWGLITYRENSLLYDDRFYAPMNK
QRIARIVAHELAHQWFGDLATMKWWDNLWLNEGFARFTEFIGAGQITKDDARMRNYFLIDV
LERALKADSVASSHPLSFRIDKAAEVEEAFDDITYAKGASVLTMLRALIGEEKHKHAVSQYLK
KFSYSNAEATDLWAVFDEVVTDVEGPDGKPMRTTEFASQWTTQMGFPVISVAEFNSTTLKL
TQSRYKANKDAVEKEKYRHPKYGFKWDIPLWYQEGDKKEIKRTWLRRDEPLYLHVNDPGA
PFVVNADRYGFYRQNHDASGWKKIIKQLKDNHEVYSPRTRNAIISDAFAAATTDAIEYETVFE
LLKYAEKETEYLPLEIAMSGISSILKYFGTEPEAKPAQTYMMNILKPMYEKSSIDFIANNYRND
KLFFQINLQKDVIDMFCALGSQDCRKKYKKLFDDEVMNKCRDGQAATECERIAAPLRSSVY
CYGVKEGGDYAFDKVMELYTAETLALEKDFLRLALGCHKDVTALKGLLLRALDRNSSFVRM
QDIPSAFNDVAANPIGGEFIFNFLIERWPDIIESIGTKHTYVEKVIPACTSGIRSQQQIDQLKNL
QKNGMNARQFGAFDKAIERAQNRVDWIKKHFQKLAAFFKKATL*
**Theoretical pI/Mw: 6.79 / 109653.12**

>Recombinant product H11-1_KB strain
MKFLVNVALVFMVVYISYIYAAHHHHHHDYKDDDDKLVPRGSRKAFDTTGGNGKEDQPIVD
DNSPSAEELRLPTTIKPLTYDLVIKTYLPNYVNYPPEKDFAIDGTVVIAMEVVEPTKSIVLNSKN
ISVIADQCELFSNNQKLDIEKIVDQPRLEKVEFVLKKKLEKNQKITLKIVYIGLINDMLGGLYRTT
YTDKDGTTKIAACTHMEPTDARLMVPCFDEPTFKANWTVTVIHPKGTSAVSNGIEKGEGEVS

GDWVTTRFDPTPRMSSYLIALVISEFKYIENYTKSGVRFRIWARPEAMKMTEYAMIAGIKCLD
YYEDFFGIKFPLPKQDMVALPDFSSGAMENWGLITYREGSVLYDENLYGPMNKERVAEVIA
HELAHQWFGNLVTMKWWDNLWLNEGFASFVEYIGADFISDGLWEMKDFFLLAPYTSGITAD
AVASSHPLSFRIDKAADVSEAFDDITYRKGASVLQMLLNLVGDENFKQSVSRYLKKFSYDNA
AAEDLWAAFDETVQGITGPNGGPLKMSEFAPQWTTQMGFPVLTVESVNATTLKVTQKRYR
QNKDAKEPEKYRHPTYGFKWDVPLWYQEDEQQVKRTWLKREEPLYFHVSNSDSSVVVNA
ERRAFCRSNYDANGWRNIMRRLKQNHKVYGPRTRNALISDAFAAAAVEEMNYETVFEMLK
YTVKEEDYLPWKEAVSGFNTILDFFGSEPESQWASEYMRKLMKPIYDKSSIKFIAENYKKDS
LFFKNNLQIAVIDTYCGLGGKECLEEMKKLFDKEVMKCQPGQQATDCVKVTAPLRKTVYCY
GVQEGGDEAFDKVMELYNAEQVQLEKDSLREALGCHKDVTALKGLLMLALDRNSSFVRLQ
GAHDVFNIVSRNPVGNELLFNFLTERWEEILESLSIRHRSVDRVIRACTRGLRSREQVQQLK
NLYKNDKRAREYGAFGGAIERSEHRVKWIEKHFRKLAAFFKKSNS*
**Theoretical pI/Mw: 6.12 / 108932.96**

>Recombinant product H11-2_KB strain
MKFLVNVALVFMVVYISYIYAAHHHHHHDYKDDDDKLVPRGSTYYFTRKAFDTTQKEQKDD
TGGKEKDNSPSAEELLLPTKIKPVSYDLSIKTHLPGYVNIPPEKNLTFDGYVEISMVVVEPTNS
IVLNSKKITLAQGGCELFSGNQKLDIESVRMERLDKLEITLKNQLQKEQKILLKITYTGLISNT
LGGLYQSIYTDTDGTTKIVAVSQNESSDARRIAPCFDEPKYKAKWTVTVVHPKGTKAASNGI
EANGNGELQGDWITSKFKTTPPMSSYLLAIIVCEFEYIEGKTETGVRFRIWSRPEAKAMTAYA
LDAGIRCLEFYEKFFDIKFPLEKQDMIALPDFTAGAMENWGLITYREDSLLYDEKIYAPMNKQ
RVALVVAHELAHLWFGNLVTLKWWDDTWLNEGFATFVEYLGMDEISHNNFRTQDFFLLDG
MDRGMRADSAASSHPLSFRVDKAAEVAEAFDDISYAKGASVLTMLRALIGEDNYRNAVVQY
LKKFSYNNAQAADLWNVFNEVVKGVKGPDGNVMKIDQFTDQWTYQMGYPVVKVEEFNAT
ALKVTQSRYKTNKDALEPEKYRNPKYGFKWDVPLWYQEGNSKEVKRTWLKRDELLYLNVN
NRDTSLVVNADRHGFYRQNYDANGWKKIIKQLKENHEVFGPRTRNAIISDAFAAATIDAIDYE
TVFELLEYAKNEEEFLPWKEALSGMFAVLKFFGNEPETKPARAYMMSILEPMYNKSSIDYIVK
NYLDDTLFTKINTQKDIIDAYCSLGSKDCIKQYKDIFYDEVMPKCKAGEAATKCVKVSAPLRA
NVYCYGVQEGGEEAFEKVMGLYLAEDIQLEKGILFKALACHKDVTALKELLLRALDRKSSFV
RLQDVPTAFRAVSENPVGEEFMFNFLMERWEEITASLETEHRAVDKVVGACCTGIRSQQQI
DQLKNLQKNNAQAKKFGSFTQEIEKGEHKIAWIKKHFHRLSEFFKRARS*
**Theoretical pI/Mw: 6.12 / 109704.83**

>Recombinant product H11-4_KB strain
MKFLVNVALVFMVVYISYIYAAHHHHHHDYKDDDDKLVPRGSTYFFTRKAFDPTQKDKNQP
GGKEKDNSPSAAELLLPSNIKPLLYDLTIKTYLPGYVNFPPEKNLTFDGRVEISMVVVEPTKSI
VLNAKKITVIPAECEVLSGTQKLDIESVKEHERLEKLGFRLKSRLEKDQKILLKITYAGLISNTL
GGIYQTTYTDANGNPKIAAVSQNEPIDGRRMVPCMDEPKYKANWTVTVIHPKGTKAASNSIE
INGEGDVSGDWITSKFETTPRMSSYLLAVFISEFDFVEGRTKQDVRFRIWSRPEAKGMTKYA
LESGIKCIEFYEDFFDIKFPLKKQDMIALPDFSAGAMENWGLITYRENSLLYDEKFYGPTNKR
RVAVVVAHELAHQWFGDLVTMKWWDDLWLNEGFATFVEYIGADEIGDHYFNMPDFFLIGAL
ERALKADSAASSHPLSFRIDKAVEVEEAFDDISYAKGASIITMLRALIGEDKHKHAVTQYLKKF
SYSNAQASDLWEVFDEVVTDIKGPDGKPMKTTAFADQWTTQMGFPLVTVEAFNATSVKISQ
SRFKTNKDAKEPEKYRHPKYGFKWDIPLWYQEGDNKEVKQTWIRREEPLYLHVNDLSKPFV
VNADRHGFYRQNYDADGWRKIIKQLRDNHKVFSPRTRNAIISDAFALASVNAIEYETVFELLK
YAVNEEEFIPWTEAISGIFAVLKFFGNEPESKPAEAYMMKILEPMYKKSDLGYIAAKYKVDQL
FSKINLQKDIIDAYCALGSKDCMKKYKDIFDREVMNKCNDGDEATKCVSVAAPLRSSTYCNG
VKAGGTYAFEKVKALYYKETVQLEKDMLLRALGCHRDVTALKGLLLLAVDRNSSFVRLQDIP
NAFQAVAANPVGEEFMFNFLIERWGDIIGSIGSEPTYVERVIPPCTSGIRSKQQIDQLRNLHK
NGIHAQEYSTFVKEIERAEHKVDWIKKHFKKLASFFKNATW*
**Theoretical pI/Mw: 6.70 / 109405.84**

>Recombinant product H11-5a_KB strain
MKFLVNVALVFMVVYISYIYAAHHHHHHDYKDDDDKLVPRGSQWTKRTVLKFTPITLLLLLFL
VAASIGLSIGLTYYFTRKAYDTTEKNKDHGADDNSPSAEELRLPKNIEPLLYDLSIKTYLPGYV
SFPPEKNLTFDGQVGISLRVVEPTKSIVLNSKNITVIPDKCELFSGDKKLEIESIKEHERLEKLEI
LLKNRLEKDQEVLLKIIYTGIISNTLGGLYQATYTDTDGTVKIAAASQNEPTDARRMVPCLDEP
SFKANWTVTVIHPKGTKAVSNGIETNGEGEVSGDWIISKFETTPRMSSYLLAVVVSEFDYIEG
FTKSGVRFRIWSRPEAMNMTGYAKDAGIRCLEYYENFFDIKFPLKKQDMVALPDFSAGAME
NWGLITYRENQLLYDDRYYGPINKQRVALVVAHELAHQWFGNLVTLKWWDDTWLNEGFAT
FVEYIGTDEISYKNFRMDDFFLPDALAIGLDADAVSSTHPLSFRVDKAAEVAEAFDEITYAKG
ASVLVMLQAVIGEKNYKQAVTQYLKKFSYSNAQATDLWNVFDEVVKDVKGPDGNLMKTTEF
ASQWTTQLGFPLVTVKAFNATSLQITQTRYKTNKDALEPEKYRHPKYGFKWDIPLWYQEGD
NKDIKLAWLTREEPLYLHVSNPDTSIVVNADRHGFYRQNYDANGWRKIIKQLKENHKAYSTR
TRNAIIGDAFAAALIDQLEYETVFELLEYAKNEEEYLPWTETISGFYAILDFFGNEPESTSAKTF
MMNILKPMYEKTSMKYIADNYKNDSLFFEMNLQKSVINAYCFLGSMECIKNYTDLFDKEVMK
KCKDDDEASKCVSIAAPLRAKAYCYGVKEGGQVAFDKVMKLYYAENVQLEKDVLLQGLGC
HKDITALKRLLLLALDRNSSFVRLQDVTDVYDAVSSNPVGEEFMFNFLLERWEEILESLTTEH
RAVERVIEACTAGIRSEQQIDQLRSLQKNGAHAREYGAFDEQIERAEHKINWIKKHLRKLSDF
FEKSTF*
**Theoretical pI/Mw: 5.48 / 113073.47**

>Recombinant product H11-5b_KB strain
MKFLVNVALVFMVVYISYIYAAHHHHHHDYKDDDDKLVPRGSTYFTRRAYDTTERNKDYCA
DYNSPSAQALRLPRNIEPLLYDLCIKTYLPNYVDFPPEKNLTFDGQVGISLRVLEPTKSIVLNA
KNITVIPNKCELFLGNQKLKIESVKEHERLEKVEFLLRNWLQKDQKVLLKIVYTGLISNTLFGLY
QATYTDPDGTVKIAAATHMEPTAARRMVPCLDEPSFKANWTVTVIHPKGTKAVSNGIETNG
KGEVSGDWIISKFETTPRMSSYLLAVVVSEFDYIEGFTKTGVRFRIWSRPEAKNMTEYARDA
GIRCLEFYENFFESKFPLKKQDMVALPDFSFGAMENWGLITYRETSLLYDDRYYGPINKQRV
ALVVAHELAHQWFGNLVTLKWWDDLWLNEGFARRVEYIGTDEINNKTIRMDDVFLPNALVK
ALDADAVSSTHPLSFRIDKAAEVLEAFDTITYEKGASVLEMLRAVIGDKNFKQAVTKYLKKFS
YRNAKPSDLWNVFDEVVKDVRGPDGNSMKTTKFAPQWTTQLGFPLVAVKDFNGTSVKITQ
TRYKANKDALEPEKYRHPKYRFKWDIPLWYQEGDNKDIKFAWLTREKPLYLHKTKPDTSIVV
NADRHGFYQQNYDAKGWRKIIKQLKKNHKAYSARTRNAIIGDAFAAALIDELEYETVFKLLEY
AKNEEEYLPWTETLSGFDAILNFFGNEPESTSAKAFMKNILKPMYKKTSMKYIAVNYKNDSLF
FEVNLQTSIIDAYCHLGARECIKNYTDLFDKEVMKKCRDGDKASKCVSIAAPLRAKAYCYGV
KEGGEVAFDKVMKLCYAENVQVEKDVLLKGLGCHRDITALKRLLLLALDRNSTFVRLQDVAA
VYDAVSANPIGKEFMFNFLLERWEEILESLTADHRTVERVIKACTAGIRLEQQIDQLRSLQKN
GEHAREYGAFDGQIERAQHKINWNKK*
**Theoretical pI/Mw: 8.62 / 108201.77**

>Recombinant product H11-5c_KB strain
MKFLVNVALVFMVVYISYIYAAHHHHHHDYKDDDDKLVPRGSADDSSPSAQELRLPRNVEP
LLYNLSIKTYLPSYVNFPPEKNLTFDGQVGISLRVVEPTKSIVLNAKNITVIPNNCELFSGGRKL
EIESVEVHERLEKVEFLLRNRLQKDQKVLLKIIYTGVISNSLFGLYQATYTDADGNVKIAAATQ
LCPSDARRLVPCLDEPSFKASWTVTVIHPKGTKAVSNGIETNGKGEVSGDWIISKFETTPRM
SSYLLAIVASEFDYVEGFTKSGVRFRIWSRPEAKNMTAYARDAGIRCLEFYENFFDIKFPLKK
QDMVVLPDFSFGAMENWGLITYRENRLLYDDRYYTPINKQLVALVVAHELAHQWFGDLVTL
KWWDDLWLNEGFARFVEYIGTDEINNKTIRMDDFFLPNALVKALDADAVSSTHPLSFRVDKA
AEVVEAFDRITYEKGASVLKMLQALIGQKNYKQAVTQYLKKFSYSNAQASDLWNVFDEVVK
DVKGPDGNLMKTTEFASQWTTQMGFPLVTVKAFNATSLQITQTRYKTNKDALEPEKYRHPK
YGFKWDVPLWYQEGDNKDIKFAWLTREKPLYLHMTKPDTTIVVNADRHGFYRQNYDANGW
RKIIKQLKKNHKAYSARTRNAIIGDAFAAARIDELEYETVFELLEYAKNEEEYLPWTEAISGFY
AILDFFGNEPESTSAKAFMKNLLKPMYDKTSMKYIADNYKNDSLFFEVNLQTSIIDAYCFLGA

RECIKNYADLFDKEVMKKCKDGDKASKCVSIAAPLRAKAYCYGVKEGGEVAFEKVMKLWYA
ENVQVEKDVLLKGLGCHRDITALKRLLLLALDRNSSFVRLQDVAAVYYAVSSNPIGKEFMFN
FHLERWEEILEGLTTEHRAVERVIKACTAGIRLEQQIDQLRSLQKNGEHAREYGAFDGQIER
AQHKINWIKKHMRKLSDFFEKSTR*
**Theoretical pI/Mw: 7.64 / 107517.72**

>Recombinant product GA1_US strain
MKFLVNVALVFMVVYISYIYAAHHHHHHDYKDDDDKLVPRGSELHFPGETHLKNVRQLTFEH
RNTEAYFNSDDTYLVFQATGYGVDCEQIYRLELSRPVETLAKISTGIGTSAASFFYPNSVDIL
YSGNFHKTRVNAKKLTNDSSCPQKVCDSSKAKTDPIIKKMCESGHAWDVFPTYDIFKVNEY
GNVLDQITKNDVYDSEAFISPDGKKIVYTSKQSGDLDLWITDINGALKIQLTKAKGYDGGASF
SPDGEKIVFHASRPTTKDKIDIYDHLLENDLVAMSEMELYVMNADGIDKRSVFKQPRGGLNW
TPYYHPDNKRIIFSSNTNSTKPSEFHLYIVNEDGTGLERVTFGSGYFNAFPVFSHDGKKVVW
SSNRGSTKKGDLNLFIADWVDPGRDTDDESDAKEELRTNRIPKKATQKNAMRWQDTVTTP
YDNVVHYTGERRLKNVKQLTFRGQNAEGYFSYDDSKIILQATGYGTTCDQIYELDLNVDPRK
QIMKRMSTGLGGTTCSFFFNEPDNNHRLYAGDFWALNDTVRDTIITHTCPAKKCENRKAIKD
PVLKELCNTVYTWDIYPEFDIFMVNKYGNIVKQLTDEPGYNAEAVLSPDGKTIAFTSIRTGDL
ELWTMNTDGTNLHQVTKEYGYDGGCFFSPDGKRLVFRASRPKTQKEKEKYQKLLDYGLVE
PTLTEIYVVDVDGKNLKQVTNFGVASWAPYYLPDNKRIIFSSNYNMSAKQFGSFALYVINED
GTGLERVTFGEGYQFNAFPMFNRAGNKLVWGSSRNRTKGASLNLFIADWVDKIDDDNIGG
GGGGDDNKKKDEKGKEKEEKKS*
**Theoretical pI/Mw: 6.04 / 91034.70**

MKFLVNVALVFMVVYISYIYA: melittin signal peptide (cleaved from mature proteins)

HHHHHHDYKDDDDK: HisFLAG duo-affinity tag

LVPRGS: thrombin recognition and cleavage site

E-P/K-EKYRHP-T/K-YGFKWD: anti-H11(16aa) binding site
