## Supplementary Figures for "Glycoengineering of nematode antigens using insect cells: a promising approach for producing bioactive vaccine antigens of the barber’s pole worm *Haemonchus contortus*"

**S1 Fig.** Sequence and 3D-structure alignment of H11 antigens. (A) Multiple sequence alignment of H11 antigens (accession numbers: CAB57357.1, CDJ83822.1, AGT79010.1, CAB57358.1, CAC39009.1, Q10737.2) showing five highlighted conserved regions. (B) AlphaFold-predicted 3D-structure of H11 antigens; (I) Surface representation highlighting two exposed motifs (shown in yellow and magenta). (II) Ribbon representation showing internal motifs (indicated in green, red, and blue).

**A**

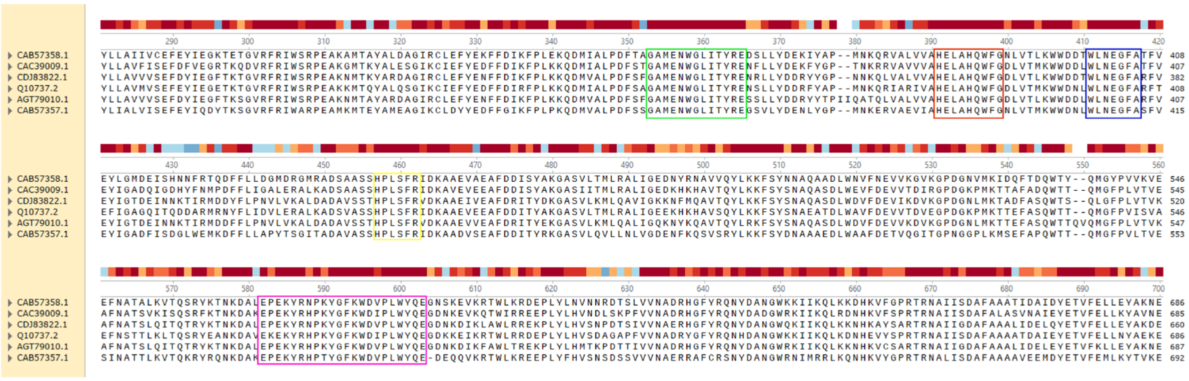


**B**

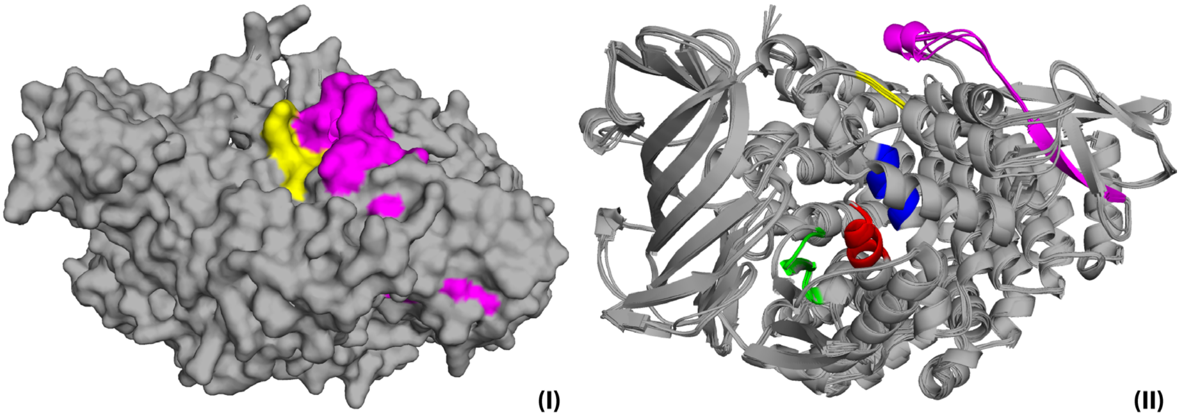

**S2 Fig.** Western blot analysis of protein samples using a rabbit anti-H11 polyclonal antibody. Samples, including a recombinantly expressed H11-1 protein, *C. elegans* lysate (N2), *Ascaris suum* lysate (extract A and B), *O. dentatum* lysate (Ösi) and *H. contortus* lysate (L3 and adult), were sequentially incubated with either purified IgG (A) or pre-immune serum (B), and an anti-rabbit IgG secondary antibody prior to colour development.

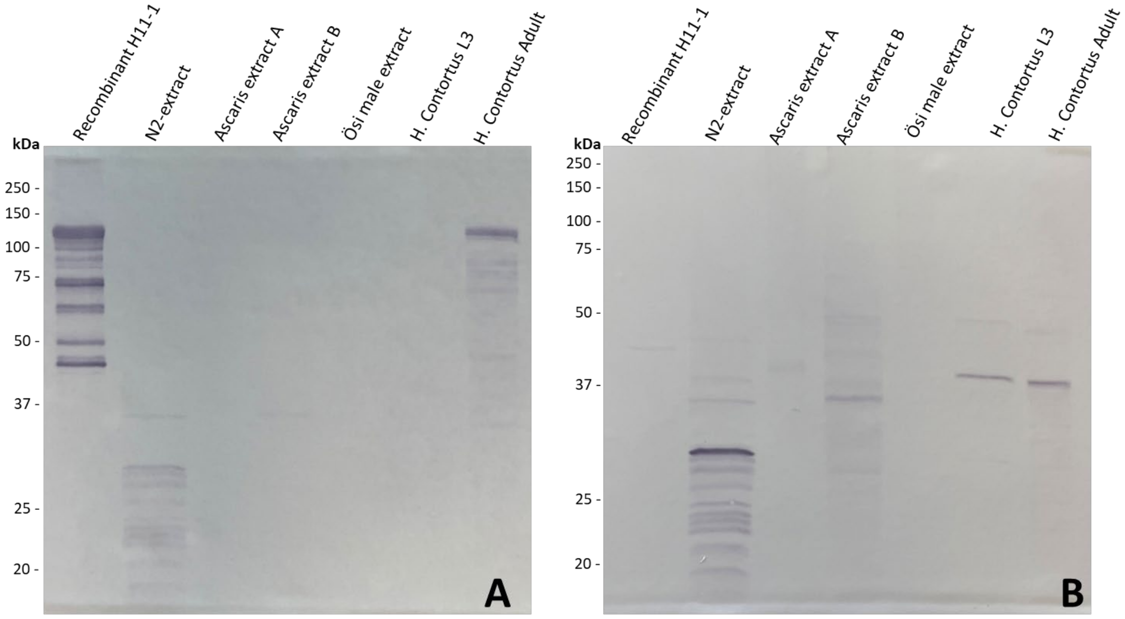

**S3 Fig.** An overview of eight DNA constructs carrying the glyco-module (*aman-3*, *fut-6*, *galt-1*) and different *H. contortus* antigen encoding genes. Constructs were verified by Nanopore DNA sequencing.

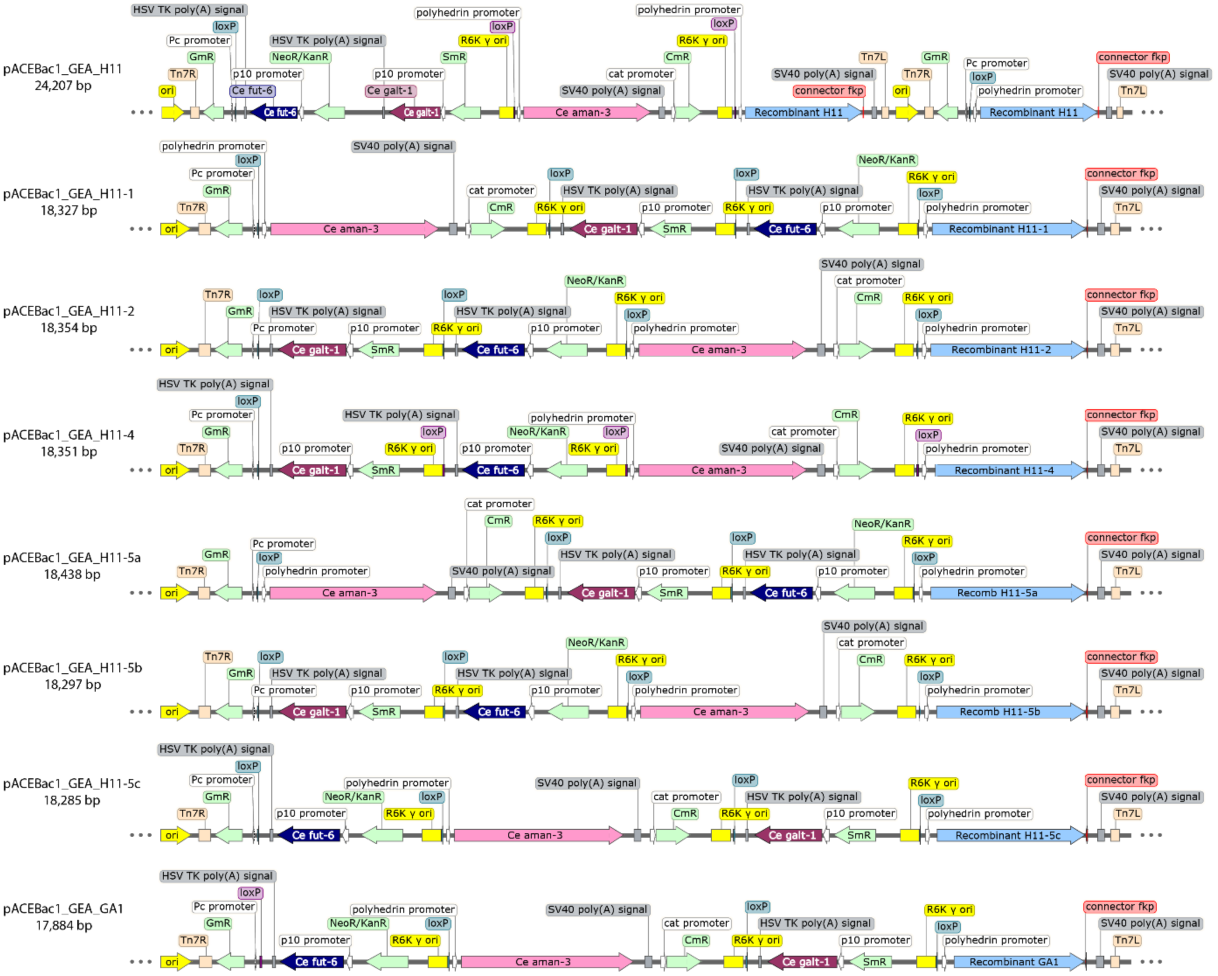

**S4 Fig.** Microscopic images of baculovirus-infected Sf9 cells. The example shown here indicated that cells infected with viruses carrying the glyco-module (geH11, left panel) have more floaters than cells infected with viruses without the glyco-module (H11, right panel). Cells infected with $V_0$ baculoviruses displayed bright fluorescent signals (lower panel) four days post infection.

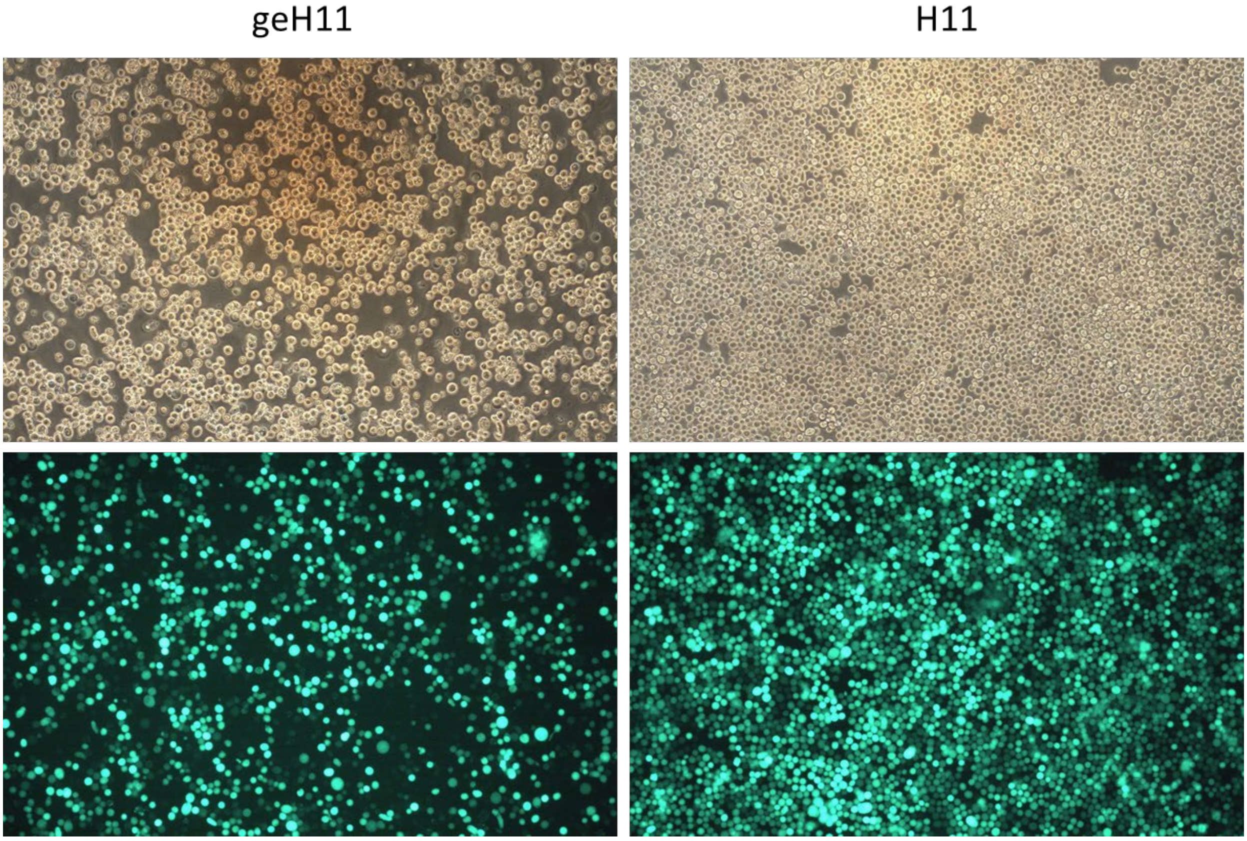

**S5 Fig.** Western blot assay of recombinant antigens treated with or without thrombin. As an example to demonstrate that it is feasible to remove the HisFLAG duo-affinity tag from recombinant antigens, the glycoengineered H11-1 antigen (KB geH11-1) was incubated with thrombin-agarose (Sigma-Aldrich, St. Louis, MO, USA) overnight at 37°C, under two conditions (b and d, at 300 rpm on a thermomixer; c and e, at 5 rpm on an overhead shaker). Untreated sample (a) was loaded as a control. Post blotting, the NC membrane was split into two (dash line), which were separately probed by either a FLAG-specific monoclonal antibody (WB 1) or a H11(16-aa motif)-specific polyclonal antibody (WB 2). The altered recognition pattern in WB 1 indicated a partial (b and d) and complete removal (c and e) of the affinity tag.

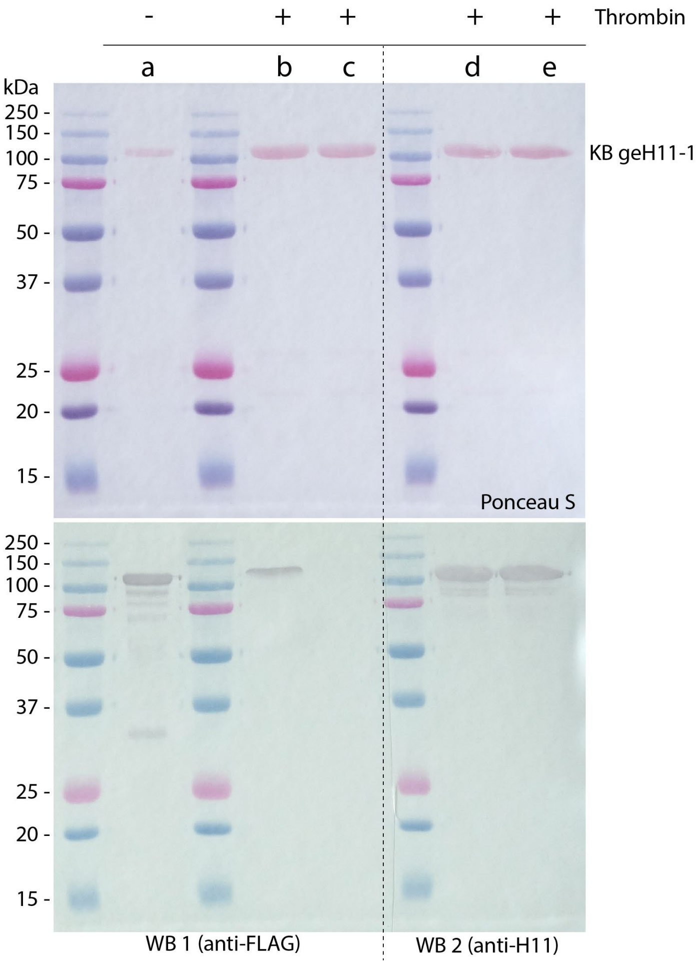

**S6 Fig.** Aminopeptidase activity of recombinant glycoengineered (ge) H11 antigens at different pH conditions. Enzymatic activities were assessed at pH 5, pH 7, and pH 8 and absorbance at OD450 nm was measured after 24-hour incubation at 37°C. The negative controls, included for assays at pH 7, were reaction mixtures contained either no substrate (Neg. Ctl 1) or no recombinant antigen (Neg. Ctl 2). Activities measured at pH 5 and pH 8 displayed lower OD values as compared to the ones at pH 7.

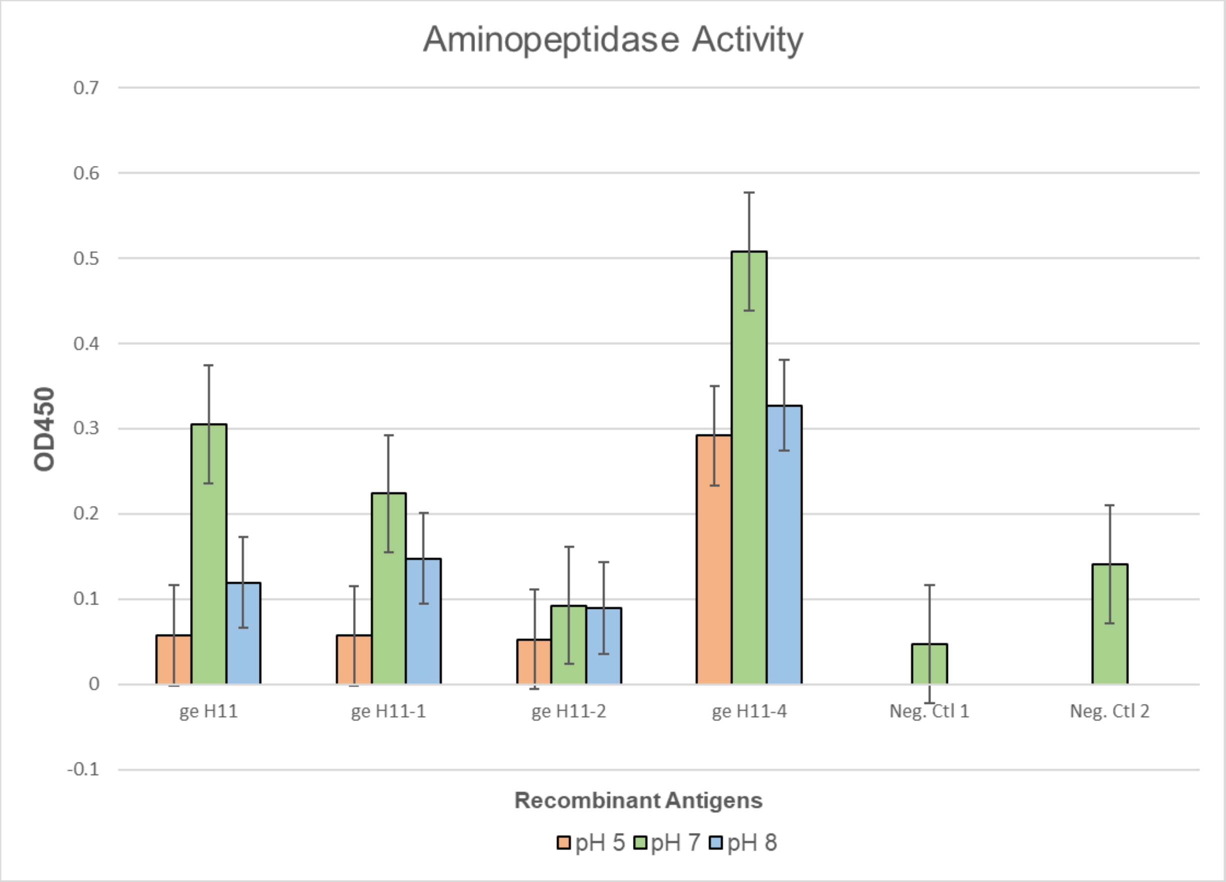

**S7 Fig.** MALDI-TOF MS spectra of native N-glycans released from individual recombinant antigens. (A) Spectra of native glycans from non-glycoengineered antigens: H11, H11-1, H11-2, H11-4, and GA1. (B) Spectra of native glycans from glyco-engineered (ge) variants: geH11, geH11-1, geH11-2, geH11-4, and geGA1. MALDI-TOF MS spectra indicated that only the glyco-engineered antigens (panel B) possess the desired tri-fucosylated N-glycan (m/z 1387), which is absent in the non-glycoengineered antigens (panel A).

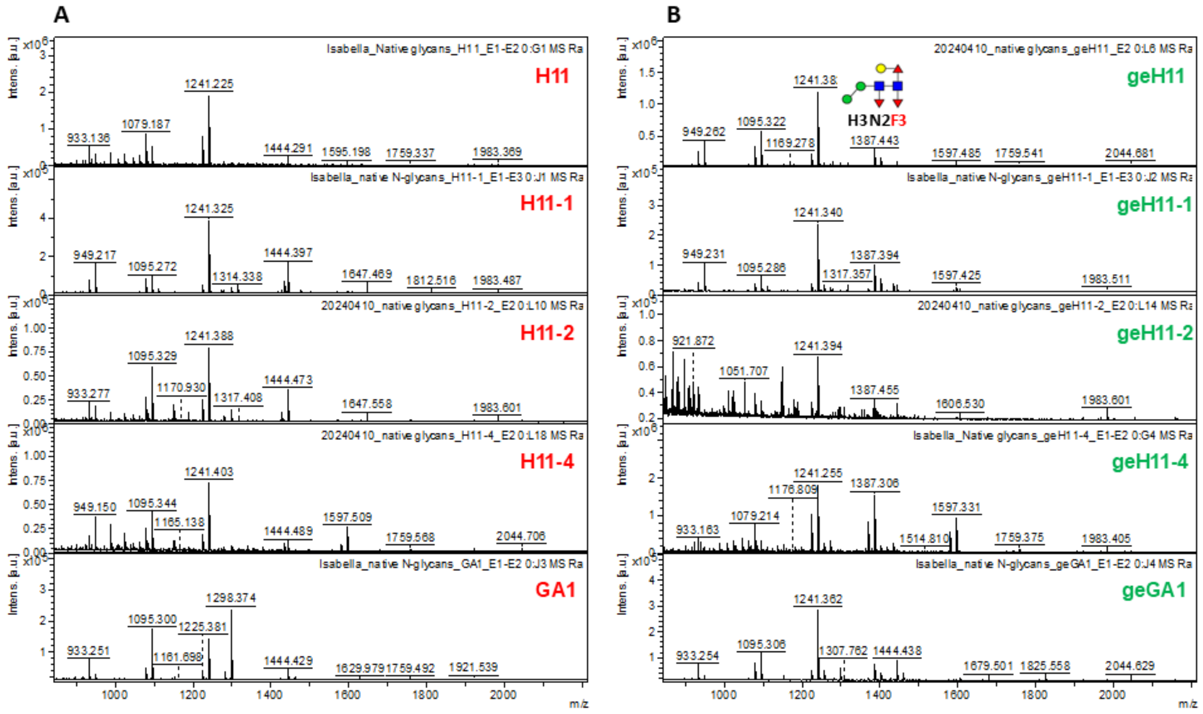

**S8 Fig.** Cytokine and chemokine expression profiles at 48h and 72h post-stimulation. Fluorescence intensity (FI) levels of IL-1α, IFNγ, IL-17A, IL-36RA, IL-4, IL-8, IP-10, and VEGF-A were quantified in response to BVAX, GEA, GEA (+), NEA (+), Quil A, Con A, and a negative control (Neg-Ctl). Data are presented as mean ± standard deviation (SD) from PBMCs of two animals. Graphs were generated using GraphPad software.

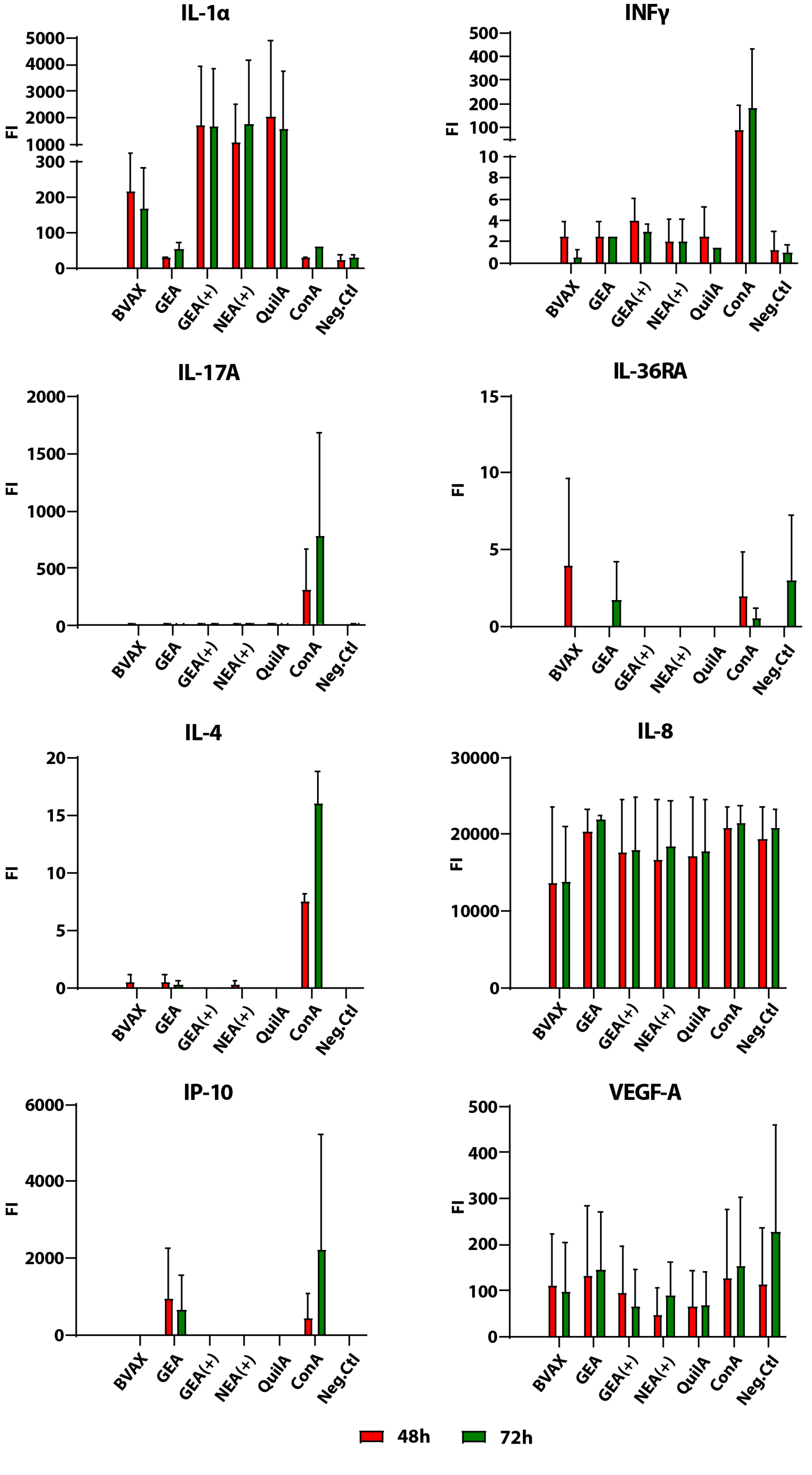
